# Biologically informed NeuralODEs for genome-wide regulatory dynamics

**DOI:** 10.1101/2023.02.24.529835

**Authors:** Intekhab Hossain, Viola Fanfani, Jonas Fischer, John Quackenbush, Rebekka Burkholz

## Abstract

Modeling dynamics of gene regulatory networks using ordinary differential equations (ODEs) allow a deeper understanding of disease progression and response to therapy, thus aiding in intervention optimization. Although there exist methods to infer regulatory ODEs, these are generally limited to small networks, rely on dimensional reduction, or impose non-biological parametric restrictions — all impeding scalability and explainability. PHOENIX is a neural ODE framework incorporating prior domain knowledge as soft constraints to infer sparse, biologically interpretable dynamics. Extensive experiments - on simulated and real data - demonstrate PHOENIX’s unique ability to learn key regulatory dynamics while scaling to the whole genome.

## Background

Biological systems are complex with phenotypic states, including those representing health and disease, defined by the expression states in the entire genome. Transitions between these states occur over time through the action of highly interconnected regulatory processes driven by transcription factors. Modeling molecular mechanisms that govern these transitions is essential if we are to understand the behavior of biological systems, and design interventions that can more effectively induce a specific phenotypic outcome. But this is challenging since we want not only to predict gene expression at unobserved time points but also to make these predictions in a way that explains any prior knowledge of transcription factor binding sites. Models that accurately encode such interactions between transcription factors (TFs) and target genes within gene regulatory networks (GRNs) can provide insights into important cellular processes, such as disease-progression and cell-fate decisions [1–4].

Given that many dynamical systems can be described using ordinary differential equations (ODEs), a logical approach to modeling GRNs is to estimate ODEs for gene expression using an appropriate statistical learning technique [3–6]. Although estimating gene regulatory ODEs ideally requires time-course data, obtaining such data in biological systems might be difficult. One can instead use pseudotime methods applied to cross-sectional data to order samples and subsequently estimate ODEs that capture the regulatory structure [7, 8].

While a variety of ODE estimation methods have been proposed, most suffer from critical issues that limit their applicability in modeling genome-wide regulatory networks. Some systems biology models (such as those built using COPASI [9]) formulate ODEs based solely on biochemical principles of gene regulation and use the available data to parameterize these equations. However, such methods impose several restrictions on the ODEs and cannot flexibly adjust to situations where the underlying assumptions do not hold; this increases the risk of model misspecification and hinders scalability to large networks, particularly given the enormous number of parameters necessary to specify a genome-scale model [10, 11]. Other methods including Dynamo, PRESCIENT, and RNA-ODE are based on non-parametric approaches to learning regulatory ODEs, using tools such as sparse kernel regression [3], random forests [5], variational auto-encoders [7, 12, 13], diffusion processes [4], and neural ordinary differential equations [6, 14], but these fail to include biologically relevant associations between regulatory elements and genes as constraints on the models.

These latter models can be broadly placed into two classes based on the inputs required to estimate the gradient *f* of the gene regulatory dynamics, where 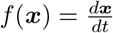. The first class consists of methods like PRESCIENT [4] and RNAForecaster [6] that can learn *f* based only on time series gene expression input 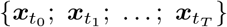 without additional steps or consideration of other regulatory inputs [4, 6, 15]. In the process of learning transitions between consecutive time points, these “one-step” methods **implicitly** learn the local derivative (in the context of single cell sequencing often referred to as “RNA velocity” [16]) 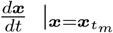, as an intermediary to estimating *f*. One significant issue with these approaches is scalability, and studying meaningfully large dynamical systems (ideally those describing the entire genome) has so far been hindered by a large performance loss and missing interpretability [4, 6, 17, 18]. This leads to potential issues with generalizability as regulatory processes operate genome-wide and even small perturbations can have wide-ranging regulatory effects.

A second class of approaches consists of “two-step” methods such as Dynamo [3], RNA-ODE [5], and DeepVelo [12] that instead only require snapshot expression data for learning *f* with two separate steps [3, 5, 12], which allows for much broader applicability to standard RNA-seq data at the cost of performance as true timecourse data is required for reliable causal inference. These approaches first **explicitly** estimate the RNA velocity 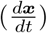 for each data point in a preprocessing step, requiring spliced and unspliced transcript counts, and one of many available velocity estimation tools [3, 16, 19–23]. In the next step, the original task of learning *f* is reduced to learning a vector field from expression-velocity tuples 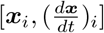 and a suitable learning algorithm is deployed. Apart from needing additional inputs that may not always be available (for example, spliced and unspliced counts are not available for microarray data), these “two-step” methods are also sensitive to the velocity estimation tool used, many of which suffer from a multitude of weaknesses [19]. Still, the Jacobian of the estimated vector field can help inform whether the learned dynamics are biologically meaningful [2, 3, 5, 8].

While the flexibility of both classes of models helps estimate arbitrary dynamics, they are “black-box” methods whose somewhat opaque nature not only makes them prone to over-fitting but also creates challenges in teasing out interpretable mechanistic insights into regulatory control [1, 6]. These models are optimized solely to predict RNA velocity or gene expression levels and so the predictions are not explainable in the sense that most cannot be related back to a sparse causal GRN. Another major issue is the scalability of these methods; because of their computational complexity, they have not yet been shown to feasibly scale up to tens of thousands of genes – and definitely not to the entire genome [3, 4, 7, 12–14]. Consequently, most of these methods either restrict themselves to a small set of highly variable genes [4, 6, 7, 12, 14] or resort to dimension-reduction techniques (PCA, UMAP, latent-space embedding, etc.) [3, 4, 7, 13] as a preprocessing step. Although dimensional reduction for feature selection has proven useful in some instances, such as in developing predictive biomarkers, such use of “metagenes” suffers from a lack of interpretability or apparent mechanistic association. Dimensional reduction results in certain biological pathways being masked in the dynamics and impedes the recovery of causal GRNs. Further, there is no obvious way to incorporate biological constraints and prior knowledge to guide model selection and prevent over-fitting when using dimensionally reduced data [1, 24]. Given that there is a strong desire among biological scientists to understand the dynamic properties of individual genes in health and disease, we focused our efforts on developing a method capable of scaling to the genome, on the original gene expression space.

We developed **PHOENIX** (**P**rior-informed **H**ill-like **O**DEs to **E**nhance **N**euralnet **I**ntegrals with e**X**plainability) as a scalable method for estimating dynamical systems governing gene expression through an ODE-based machine learning framework that is flexible enough to avoid model misspecification and is guided by insights from systems biology that facilitate biological interpretation of the resulting models [25, 26]. At its core, PHOENIX models temporal patterns of gene expression using neural ordinary differential equations (NeuralODEs) [27, 28], an advanced computational method commensurate with the scope of human gene regulatory networks – with more than 25,000 genes and 1,600 TFs – and a limited number of samples. We implement an innovative NeuralODE architecture that inherits the universal function approximation property (and thus the flexibility) of neural networks while resembling Hill–Langmuir kinetics, which have been used to model dynamic transcription factor binding site occupancy [10, 29, 30]. It can hence reasonably describe gene regulation by modeling the sparse yet synergistic interactions of genes and transcription factors. Importantly, PHOENIX operates on the original gene expression space and performs without any dimensional reduction, thus preventing information loss, especially for lowly-expressed genes that are nonetheless important for cell fate [4].

In the optimization step of PHOENIX, we introduce user-defined prior knowledge in the form of a “network prior.” Here, we derive a prior based on TF binding motif enrichment that is predicated on the understanding that the direct regulators of gene expression are transcription factors that bind in the region around a gene’s transcription start site (TSS). Because most transcription factors bind to distinct sequence motifs, each transcription factor has the potential to regulate only a fixed subset of genes. This regulatory constraint can be expressed as a network prior by mapping transcription factors to the promoter sequence of regulated genes, using tools such as FIMO[31] and GenomicRanges [32].

The incorporation of user-defined prior knowledge of likely network structure ensures that a trained PHOENIX model is explainable – it not only predicts temporal gene expression patterns but also encodes an extractable GRN that captures key mechanistic properties of regulation such as activating (and repressive) edges and strength of regulation.

## The PHOENIX model

Given a time series gene expression data set, the NeuralODEs of PHOENIX implicitly estimate the local derivative (RNA velocity) at an input data point with a neural network (NN). We designed activation functions that resemble Hill-kinetics and thus allow the NN to sparsely represent different patterns of transcriptional co-regulation by combining separate additive and multiplicative blocks that operate on the linear and logarithmic scales respectively. An ODE solver then integrates the estimated derivative to reconstruct the steps taken from an input *x*_*i*_ at time *t*_*i*_ to a predicted output 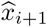 at time *t*_*i*+1_ [27]. The trained neural network block thus encodes the ODEs governing the dynamics of gene expression and hence encodes the underlying vector field and GRN. An important advantage of incorporating an ODE solver is that we can predict expression changes for arbitrarily long time intervals without relying on predefined Euler discretizations, as is required by many other methods [4, 12, 18]. We further augmented this framework by allowing users to include prior knowledge of gene regulation in a flexible way, which acts as a domain-knowledge-informed regularizer or soft constraint of the NeuralODE [24] (**Figure 1**). By combining the mechanism-driven approach of systems biology-inspired functional forms and prior knowledge with the data-driven approach of powerful machine learning tools, PHOENIX scales up to full-genome data sets and learns meaningful models of gene regulatory dynamics.

**Figure 1.**
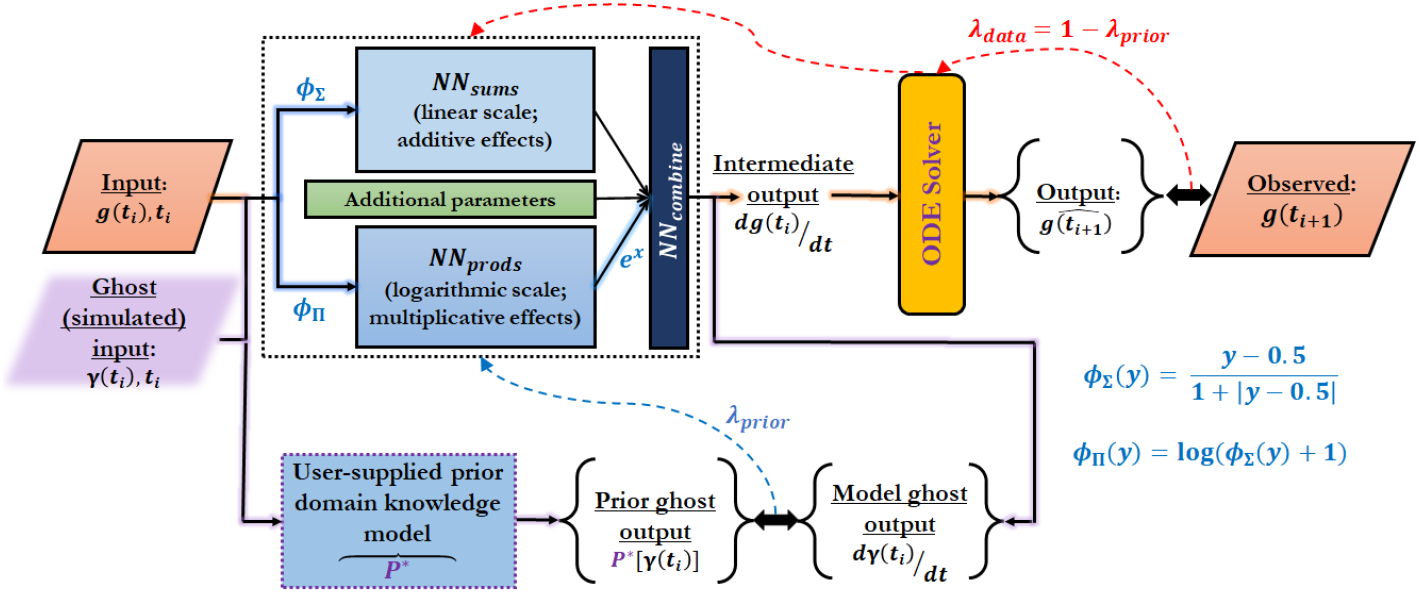
PHOENIX is powered by a NeuralODE engine. Given an expression vector ***g***(*t*_*i*_) *∈* R^#genes^ at time *t*_*i*_, a neural network (dotted rectangle) estimates the local derivative *d****g***(*t*_*i*_)*/dt* and an ODE solver integrates this value to predict expression at subsequent time points ***ĝ*** (*t*_*i*+1_). The neural network is equipped with activation functions (*ϕ*_Σ_ and *ϕ*_Π_) that resemble Hill-Langmuir kinetics, and two separate single-layer blocks (NN_*sums*_ and NN_*prods*_) that operate on the linear and logarithmic scales to model additive and multiplicative co-regulation respectively. A third block (NN_*combine*_) then flexibly combines the additive and multiplicative synergies. PHOENIX incorporates two levels of back-propagation to parameterize the neural network while inducing domain knowledge-specific properties; the first (red arrows with weight *λ*_data_) aims to match the observed data, while the second (blue arrow with weight *λ*_prior_) uses simulated expression vectors ***γ***(*t*_*i*_) *∈* ℝ^#genes^ to implement soft constraints defined by user-supplied prior models (𝒫^*∗*^) of putative regulatory interactions. Since the ***γ***(*t*_*i*_)s were simulated expression values, we also refer to them as “ghost inputs.” More details about their operationalization can be found in Supp. Methods 1.2.

### Neural ordinary differential equations (NeuralODEs)

NeuralODEs [27] learn dynamical systems by parameterizing the underlying derivatives with neural networks:

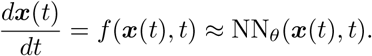

Given an initial condition, the output at any given time-point can now be approximated using a numerical ODE solver 𝒮 of adaptive step size

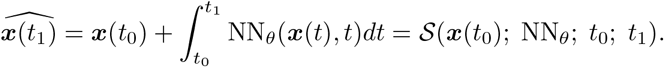

This is the basic architecture of a NeuralODE [27], and it lends itself to loss functions *L* (here, *l*_2_ loss) of the form

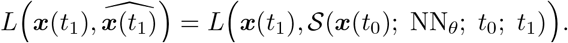

To perform back-propagation, the gradient of the loss function with respect to all parameters *θ* must be computed, which is done using the adjoint sensitivity method [27]. Building off of the NeuralODE author’s model implementation in PyTorch [28], we made biologically motivated modifications to the architecture and incorporated user-defined prior domain knowledge, as described below.

### Model formulation and neural network architecture

Most models for co-regulation of gene expression are structured as a simple feedback process [29]. Given that gene regulation can be influenced by perturbations across an entire regulatory network of *n* genes, the gene expression of all genes *g*_*j*_(*t*) can affect a specific *g*_*i*_(*t*) at time point *t*:

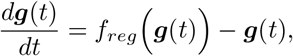

where 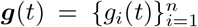, *f*_*reg*_ : ℝ^*n*^ *→* ℝ^*n*^, and *f*_*reg*_ is approximated with a neural network. To model additive as well as multiplicative effects within *f*_*reg*_, we used an innovative neural network architecture equipped with activation functions that emulate - and can thus sparsely encode - Hill-kinetics (see **Figure 1**). The Hill–Langmuir equation *H*(*P*) was originally derived to model the binding of ligands to macromolecules [30], and can be used to model transcription factor occupancy of gene regulatory binding sites [10]:

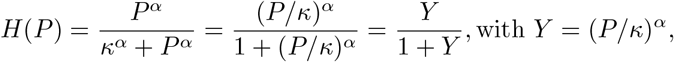

which resembles the softsign activation function *ϕ*_soft_(*y*) = 1*/*(1 + |*y*|). For better neural network trainability, however, we shifted it to the center of the expression values. To approximate suitable exponents *α*, we further log-transformed *H*, since composing additive operations in the log-transformed space with a Hadamard exp *○* function can represent multiplicative effects.

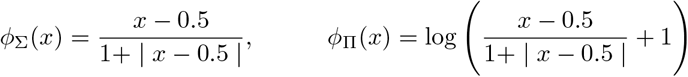

were employed as activation functions to define two neural network blocks (NN_*sums*_ and NN_*prods*_), representing additive and multiplicative effects

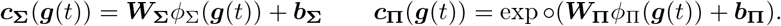

The concatenated vectors ***c***_**Σ**_(***g***(*t*)) *⊕* ***c***_**Π**_(***g***(*t*)) served as input to a third block NN_*combine*_ (with weights ***W*** _*∪*_ *∈* ℝ^*n×*2*m*^) that flexibly combined these additive and multiplicative effects. We found that a single linear layer was sufficient for this purpose. Given that ODEs have difficulty with learning zero gradients, we found it necessary (see Supp. Fig. 5) to introduce gene-specific multipliers ***υ*** *∈* ℝ^*n*^ for modelling steady states of genes that do not exhibit any temporal variation 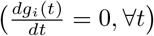.

Accordingly, the output derivative for each gene *i* was multiplied with ReLU(*υ*_*i*_) = max{(*υ*_*i*_, 0)}. We expressed this using the Hadamard product (*⊙*) of the previous output and the elementwise ReLU of ***υ*** As

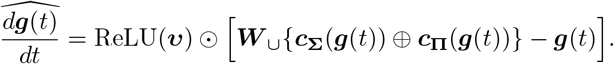

The trainable parameters ***θ*** = (***W***_**Σ**_, ***W***_**Π**_, ***b***_**Σ**_, ***b***_**Π**_, ***W*** _*∪*_, ***υ***) were learned based on observed data and prior domain knowledge (details in Supp. Methods 1).

### Structural domain knowledge incorporation

One challenge we found in interpreting PHOENIX is that NeuralODEs have multiple solutions [33], of which many are inconsistent with our understanding of the process by which specific transcription factors (TFs) regulate the expression of other genes within the genome. Most solutions accurately represent gene-gene correlations, but do not necessarily reflect biologically established TF-gene regulation processes. Inspired by recent developments in physics-informed deep learning [24], we introduced biologically motivated soft constraints to regularize the search for a parsimonious approximation. We started with the NeuralODE prediction for the gene expression vector

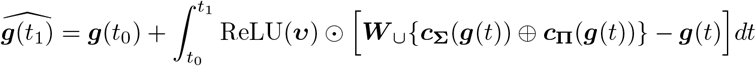

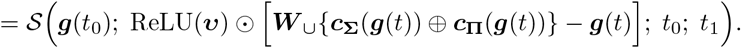

We found that the unregularized PHOENIX provides an observed gene expression-based approximation for the local derivative 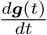, but often we have additional structural information available about which TFs are more likely to regulate certain target genes. Hence, one could also formulate a domain knowledge-informed 𝒫^*∗*^ (***g***(*t*)) that is a prior-based approximation as

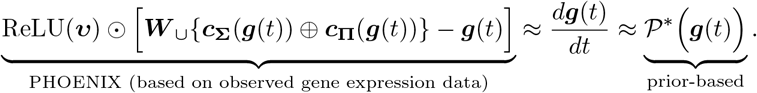

By promoting our NeuralODE to flexibly align with such structural domain knowledge, we automatically searched for biologically more realistic models that still explained the observed gene expression data. To this end, we designed a modified loss function ℒ_mod_ that incorporated the effect of prior model 𝒫^*∗*^ using a set of *K* simulated (“ghost”) expression vectors 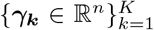. This induced a preference for consistency with prior domain knowledge.

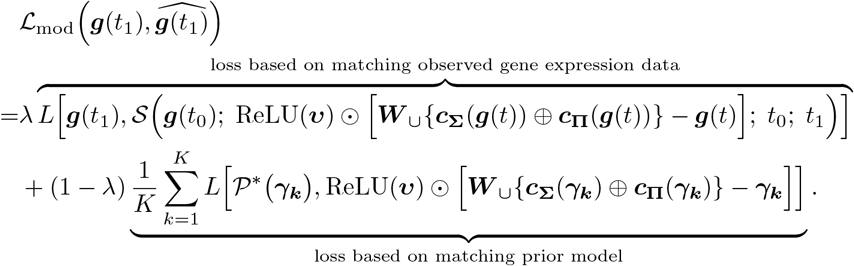

Here, *λ* is a tuning parameter for flexibly controlling how much weight is given to the prior-based optimization, which we tuned with cross-validation, and 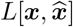 is the primary loss function, set to the *L* = *l*_2_ loss in our experiments.

While our modeling framework is flexible regarding the nature of the prior model 𝒫^*∗*^, we incorporated a simple linear model, a common choice for chemical reaction networks or simple oscillating physical systems [34] as

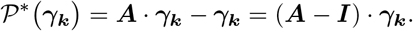

Here, ***A*** is the adjacency matrix of likely network structure based on prior domain knowledge, such as experimentally validated interactions, TF—gene binding information derived from motif scans, etc., with ***A***_*ij*_ ∈ {+1, *−*1, 0} representing an activating, repressive, or no prior interaction, respectively. For cases where the signs (activating/repressive) of prior interactions were unknown, we found that formulating ***A*** simply based on *prior interaction existence*, or ***A***_*ij*_ ∈ {1, 0}, would suffice (see Supp. Table 2).

## Results

We demonstrated the utility of PHOENIX for estimating gene expression dynamics by performing a series of *in silico* benchmarking experiments, where PHOENIX exceeded even the most optimistic performance of popular black-box RNA dynamics estimation methods. We demonstrated the scalability of PHOENIX by applying it to genome-scale breast cancer microarray samples ordered in pseudotime and investigated how scaling to the complete data set improves a representation of key pathways. We further applied PHOENIX to yeast cell cycle microarray data to show that it can capture oscillatory dynamics by flexibly deviating from Hill-like assumptions when necessary. Finally, to test PHOENIX with a different data modality, we investigated avalaible genome-scale timecourse RNASeq data of Rituximab treated B-cells, where PHOENIX is able to capture key molecular changes in the main mechanism of action of Rituximab.

### PHOENIX accurately and explainably learns temporal evolution of *in silico* dynamical systems

We began our validation studies with simulated gene expression time-series data so that the underlying dynamical system that produced the system’s patterns of gene expression was known. We adapted SimulatorGRN [29, 35] to generate time-series expression data from two synthetic *S. cerevisiae* gene regulatory systems (SIM350 and SIM690, consisting of 350 and 690 genes respectively). The activating and repressive interactions in each *in silico* system were used to synthesize noisy expression “trajectories” for each gene across multiple time points (see Methods 1.1 and 1.2). We split up the trajectories into training (88%), validation (6% for hyperparameter tuning), and testing (6%), and compared PHOENIX predictions on the test set against the “known”/ground truth trajectories. Since PHOENIX uses user-defined prior knowledge as a regularizer, we also corrupted the prior model at a level commensurate with the “experimental” noise level (see Supp. Methods 4.2), reflecting the fact that transcription factor-gene binding is itself noisy.

We found that PHOENIX accurately learned the temporal evolution of the SIM350 and SIM690 systems (**Figure 2**) and was able to recover the **true** test set trajectories (that is, test set trajectories pre-noise) with a reasonably high accuracy even when the training trajectories included high levels of noise and the prior knowledge model was strongly corrputed. Furthermore, the shapes of the predicted trajectories (and hence the predicted steady-state levels) obtained from feeding initial values (expression at *t* = 0) into the trained model remained robust to noise, suggesting that a trained PHOENIX model could be used to estimate the temporal effects of cellular perturbations.

Since the primary prediction engine of PHOENIX is a NeuralODE, we wanted to benchmark its performance relative to “out-of-the-box” (OOTB) NeuralODE models (such as RNAForecaster [6]) to understand the contributions of our modifications to the NeuralODE architecture. We tested a range of OOTB models where we adjusted the total number of trainable parameters to be similar to that of PHOENIX (see Supp. Methods 3.2). Because PHOENIX uses a domain prior of likely gene-regulation interactions in its optimization scheme, we also tested a version (PHX_0_) where the weight of the prior was set to zero (*λ*_prior_ = 0). For each of SIM350 and SIM690, we observed that PHOENIX outperformed OOTB NeuralODEs on the test set in noiseless settings (Supp. Fig. 2 and Supp. Table 3). When we added noise, the PHOENIX models still generally outperformed the OOTB models, especially PHX_0_. The test MSEs were more comparable between all the models in very high noise settings. The consistently strong performance of PHOENIX suggests that using a Hill-kinetics-inspired architecture better captures the dynamics of the regulatory process, in part because it models the binding kinetics of transcription factor-gene interactions.

**Figure 2.**
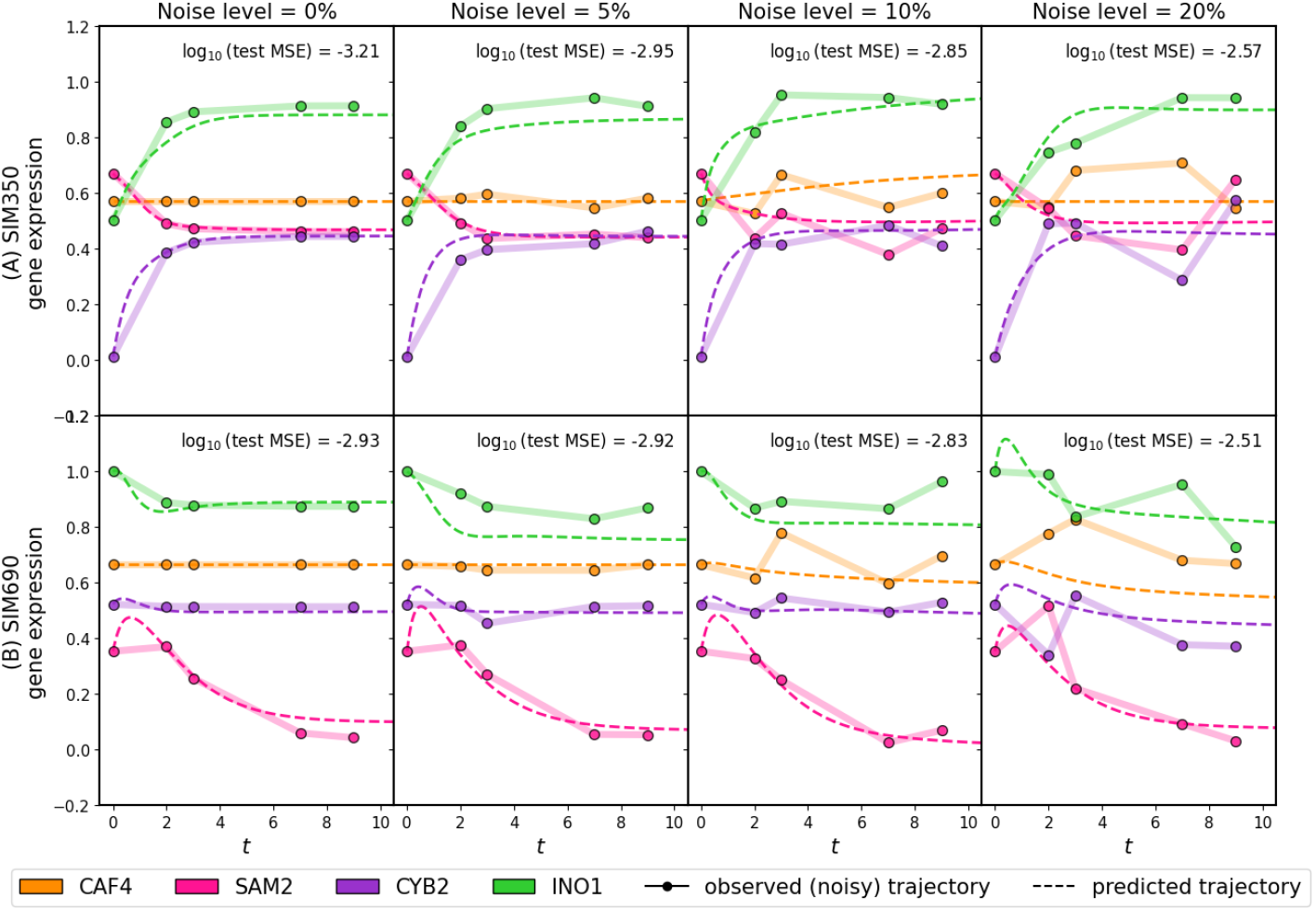
We applied PHOENIX to simulated gene expression data originating from *in silico* dynamical systems SIM350 (A) and SIM690 (B) that simulate the temporal expression of 350 and 690 genes respectively. Each simulated trajectory consisted of five time points (*t* = 0, 2, 3, 7, 9) and was subjected to varying levels of Gaussian noise (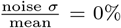, 5%, 10%, 20%, higher noise settings in Supp. Fig. 1). Since PHOENIX uses a user-defined prior network model as a regularizer, we also corrupted the prior models up to an amount commensurate with the noise level. For each noise setting we trained PHOENIX on 140 of these “observed” trajectories and validated on 10. The performance on the validation trajectories was used to determine the optimal value of *λ*_prior_. We then tested the trained model on 10 new test set trajectories. We display both observed and predicted test set trajectories for four arbitrary genes in both SIM350 and SIM690, across all noise settings. We display the mean squared error (MSE) between the predictions and the 10 pre-noise test set trajectories.

**Figure 3.**
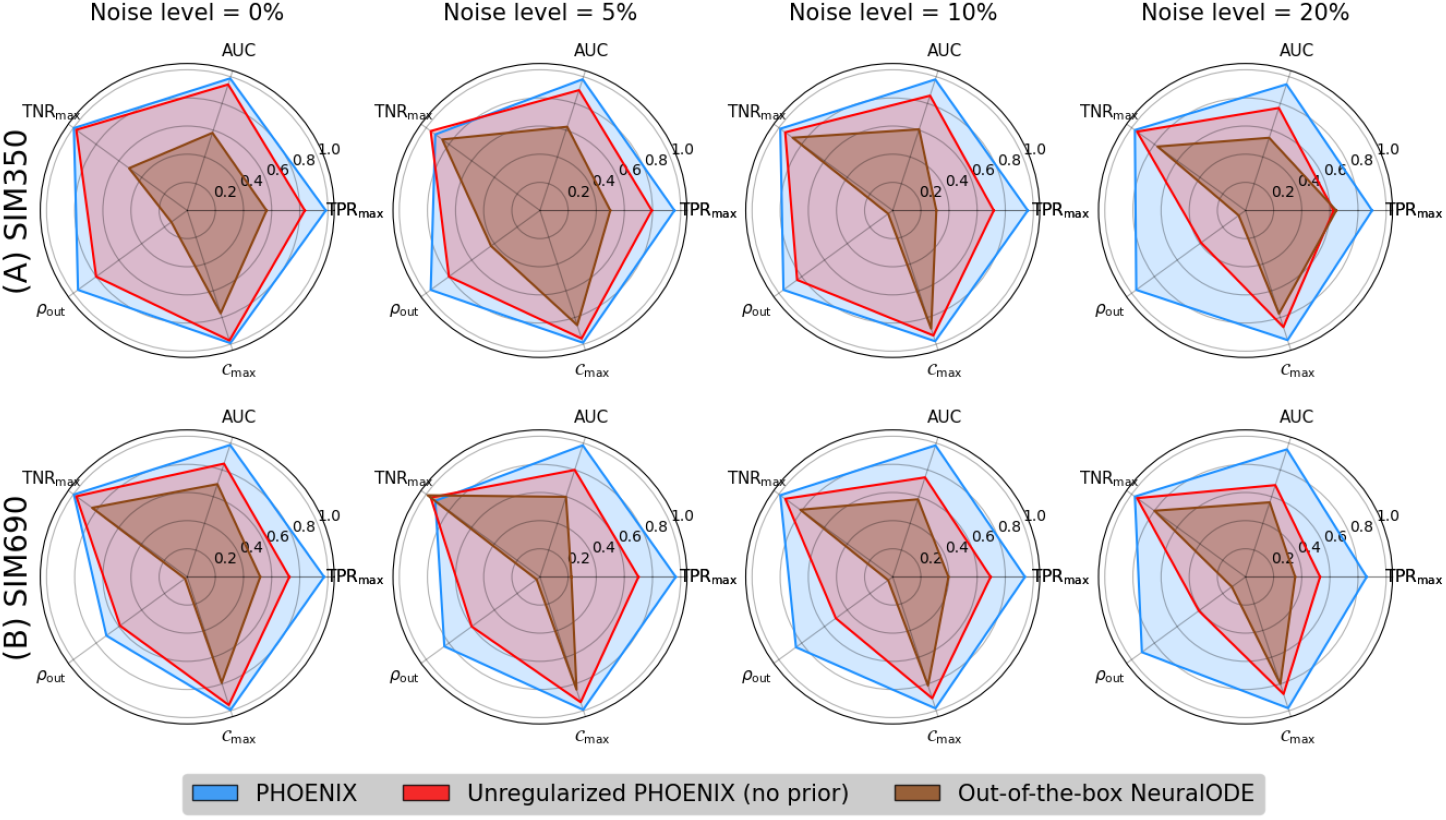
We extracted encoded GRNs from the trained PHOENIX models and the best-performing out-of-the-box NeuralODE models, for both *in silico* dynamical systems SIM350 (A) and SIM690 (B) across all noise settings (see Supp. Fig. 3 for more noise settings). We compared these GRN estimates to the corresponding ground truth GRNs used to formulate SIM350 and SIM690, and obtained AUC values as well as out-degree correlations (*ρ*_out_). We also reverse-engineered a metric (*C*_max_) to inform how sparsely PHOENIX had inferred the dynamics (see Supp. Methods 2). Furthermore, we used these *C*_max_ values to obtain optimal true positive and true negative rates (TPR_max_ and TNR_max_) that were independent of any cutoff value, allowing us to compare between “best possible” networks across all settings.

In terms of the contribution of the prior constraints to PHOENIX’s performance, we saw that PHOENIX was generally outperformed by PHX_0_, its unregularized version (Supp. Fig. 2 and Supp. Table 3). However, given that the prior can be interpreted as soft biological constraints on the estimated dynamical system [24], an important question is whether PHX_0_ (as well as OOTB models) makes accurate temporal predictions by correctly learning elements of the causal biology, or whether the lack of prior information results in an alternate learned representation of the dynamics, which - despite predicting these particular trajectories well - does not explain the true biological regulatory process.

To this end, we recognized that the parameters of a trained PHOENIX model encode an estimate of the ground-truth gene regulatory network (GRN) that causally governs the system’s evolution over time. We therefore inferred encoded GRNs from trained PHOENIX models and compared them to the ground truth networks *GRN*_350_ and *GRN*_690_ used to synthesize SIM350 and SIM690 respectively (see Supp. Methods 2). Given PHOENIX’s simple NeuralODE architecture, we were able to develop a GRN inference algorithm that could predict edge existence, direction, strength, and sign, using just model coefficients, without any need for time-consuming sensitivity analyses (unlike other approaches [2, 12]). For comparison, we wanted to extract GRNs from the most predictive OOTB models; given their black-box nature, OOTB model GRNs had to be obtained via sensitivity analyses (see Supp. Methods 3.2).

We compared inferred and ground truth GRNs in terms of several metrics, including edge recovery, out-degree correlations, and induced sparsity. We obtained near-perfect edge recovery for PHOENIX (AUC *∈* [0.96, 0.99]) as well as high out-degree correlations across all noise settings (**Figure 3** and Supp. Table 4). Most notably, we observed that PHOENIX predicted dynamics in a more robustly explainable way than PHX_0_ and the OOTB models. We measured induced sparsity by reverse engineering a metric *C*_max_ based on maximizing classification accuracy (see Supp. Methods 2), and found that PHOENIX resulted in much sparser dynamics than PHX_0_ (Supp. Table 5). To further assess this phenomenon, we computed the estimated model effect between every gene pair in SIM350, and compared these values between PHOENIX and PHX_0_. We found that the incorporation of priors helped PHOENIX identify core elements of the dynamics, and predict gene expression patterns in a biologically parsimonious manner (Supp. Fig. 4).

Since the inclusion of such static prior knowledge greatly increased the explainability of the inferred dynamics, we also investigated how explainability was affected by misspecification of the prior. In our *in silico* experiments, we had randomly corrupted (misspecified) the prior by an amount commensurate with the noise level (see Supp. Methods 4.2). We compared network representations of these misspecified prior constraints to GRNs extracted from the PHOENIX models that used these very priors. We found that PHOENIX was able to appropriately learn causal elements of the dynamics beyond what was encoded in the priors (Supp. Table 1). This suggests that even though the user-defined priors enhance explainability, PHOENIX can deviate from them when necessary, and learn regulatory interactions from just the data itself.

### PHOENIX exceeds the most optimistic performances of current black-box methods *in silico*

Having established PHOENIX models as both predictive and explainable, we compared its performance to other existing methods for gene expression ODE estimation *in silico* (Supp. Table 7). As discussed earlier, these can be placed into two groups based on the input data. The “one-step” methods estimate dynamics by directly using expression trajectories; these include RNAForecaster [6] (which is an out-of-the-box NeuralODE), and PRESCIENT [4], among others [14, 15]. PHOENIX is more similar to these methods.

“Two-step” methods such as Dynamo [3], RNA-ODE [5], and DeepVelo [12] were not directly designed for time series data, but are applicable to snapshot RNA-seq with quantified isoforms. They estimate dynamics by first reconstructing RNA velocity using inputs such as spliced and unspliced mRNA counts and then estimating a vector field mapping expression to velocity. While this comparison is not the ideal setting for such two step approaches, we did compensate for their disadvantage by providing the **ground truth** velocities as input – information that none of the one-step approaches have – into their second step (see Supp. Methods 3.1). Further, we used the validation set to optimize key hyperparameters of all the methods (Supp. Table 7, right-most column) before finally testing predictive performance on expression values from held-out test trajectories. Most of the methods also provide a means for extracting a gene network that we used to evaluate each method’s explainability (see Supp. Methods 3.3).

In these comparisons, we confirm that the “one-step” trajectory-based methods generally yield better predictions than the “two-step” velocity-based methods (although Dynamo sometimes achieved performance compared to the single-step methods), which comes as little surprise as these methods were originally designed for a slightly different setting. Overall, at reasonable noise levels, PHOENIX outperformed even the optimistic versions of the black-box methods by large margins both in terms of predicting gene expression (Supp. Table 3) and explainability (based on consistency with the ground truth network; Supp. Table 4). We found that Dynamo was the most explainable competing method in SIM350 but that, in SIM690, DeepVelo was more explainable. Finally, we found that the dynamics estimated by PHOENIX were generally much sparser than any other method and that sparsity generally decreased with noise levels (Supp. Table 5).

Further ODE estimation approaches (not included in our experiments) and their functionalities are discussed in Supp. Table 6. Code for performing such methodological benchmarks is included with the PHOENIX release [36].

### PHOENIX predicts temporal evolution of yeast cell-cycle genes in an explainable way

We tested PHOENIX using an experimental data set [37] from cell-cycle synchronized yeast cells, consisting of two technical replicates of expression values for 3551 genes across 24 time points (see Methods 2.1 for data processing).

Since there were two technical replicates (or trajectories), we used one of the two yeast replicates for training, and the other replicate for testing (not seen in any way during training). Furthermore, within the replicate for training, we used the contiguous segment of time points *t* = 45, 50, 55 minutes for validation (to choose *λ*_prior_). This scheme ensured that the train and validation sets were disjunct and that we were measuring predictive performance on a test set that was independent of the training data (see Methods 2.2). For the domain prior we used a simple adjacency-matrix-based prior model derived from TF motif enrichment analyisis in promoters of each of the 3551 genes (see Methods 2.2). We tuned the prior weight (to *λ*_prior_ = 0.05) using the validation set to induce higher explainability by promoting a biologically anchored structure to the dynamics.

PHOENIX was able to learn the temporal evolution of gene expression across the yeast cycle explaining over 69% of the variation in the test set (**Figure 4B** and **Table 1**). Notably, when we visualized the estimated dynamics by extrapolating from just initial values (expression at *t* = 0), we found that PHOENIX plausibly predicted continued periodic oscillations in gene expression, even though the training data consisted of only two full cell cycles (**Figure 4A**). The amplitude of the predicted trajectories dampened across time points, which is expected given that yeast array data tends to exhibit underdamped harmonic oscillation during cell division possibly reflecting de-synchronization of the yeast cells over time [38]. This performance on non Hill-like oscillatory dynamics is indicative of the high flexibility of PHOENIX. It inherits the universal function approximation property from NeuralODEs, allowing it to deviate from Hill-like assumptions when necessary, while still remaining explainable due to the integration of prior knowledge.

**Figure 4.**
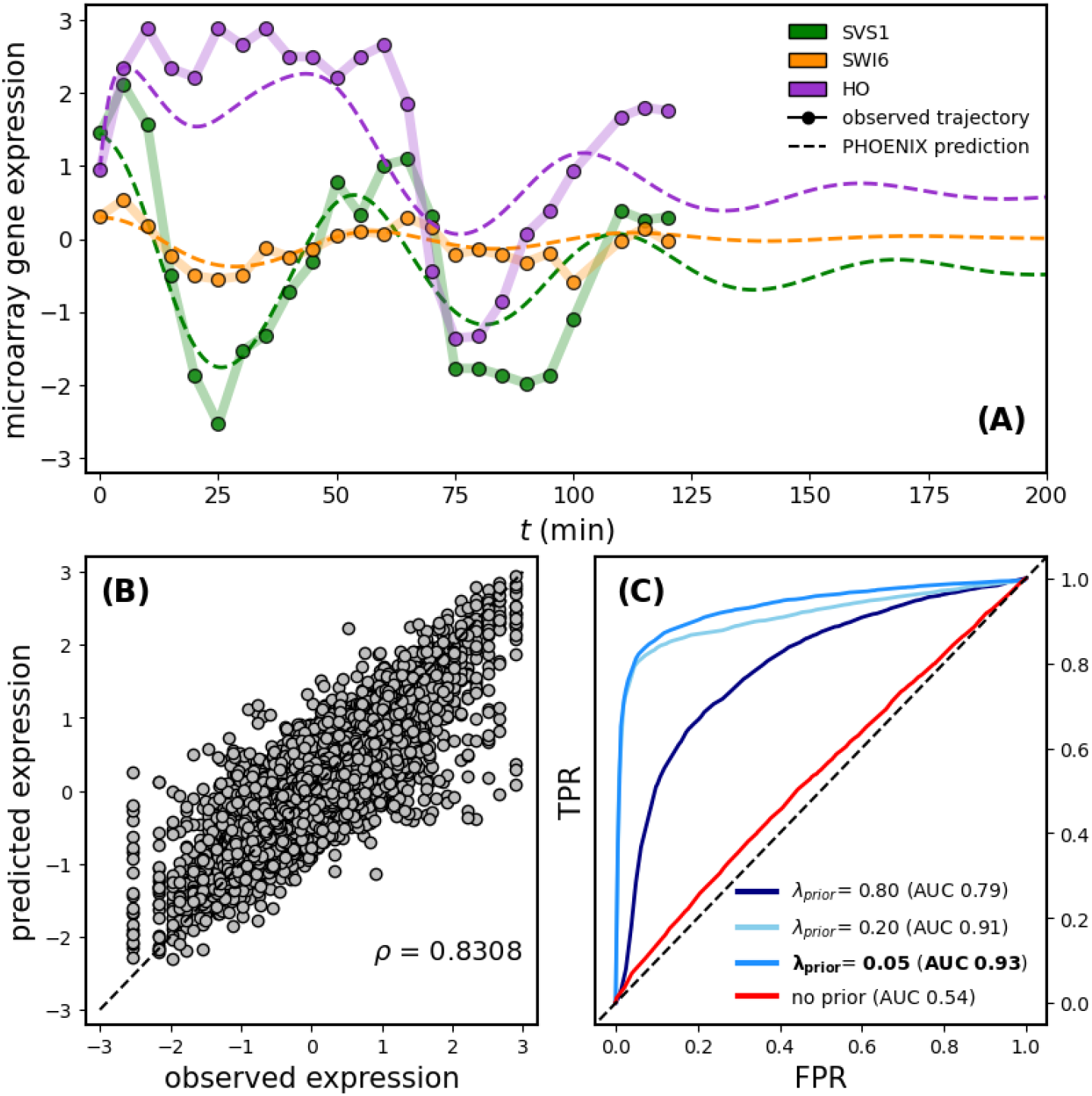
**(A)** We applied PHOENIX (*λ*_*prior*_ = 0.05) to 2 technical replicates of gene expression of 3551 genes each, collected across 24 time points in a yeast cell-cycle time course [37]. We trained on 40 transition pairs, used 3 for validation, and tested predictive accuracy on the remaining 3. We display both observed and predicted trajectories for 3 arbitrary genes, where the predicted trajectories are extrapolations into future time points based on just initial values (gene expression at *t* = 0). Additional genes plotted in Supp. Fig. 6. **(B)** We correlated observed versus predicted expression levels of all 3551 genes for the 3 expression vectors in the test set; *ρ* = 0.8308 implying *R*^2^ = 0.69. **(C)** We tested the explainability of the learned dynamics by comparing encoded GRNs retrieved from a series of trained models (of varying prior dependencies) against ChIP-chip data [39] to obtain ROC curves. The *λ*_prior_ = 0.05 model was the one chosen based on the validation set MSE (see Supp. Methods 1).

**Table 1.**
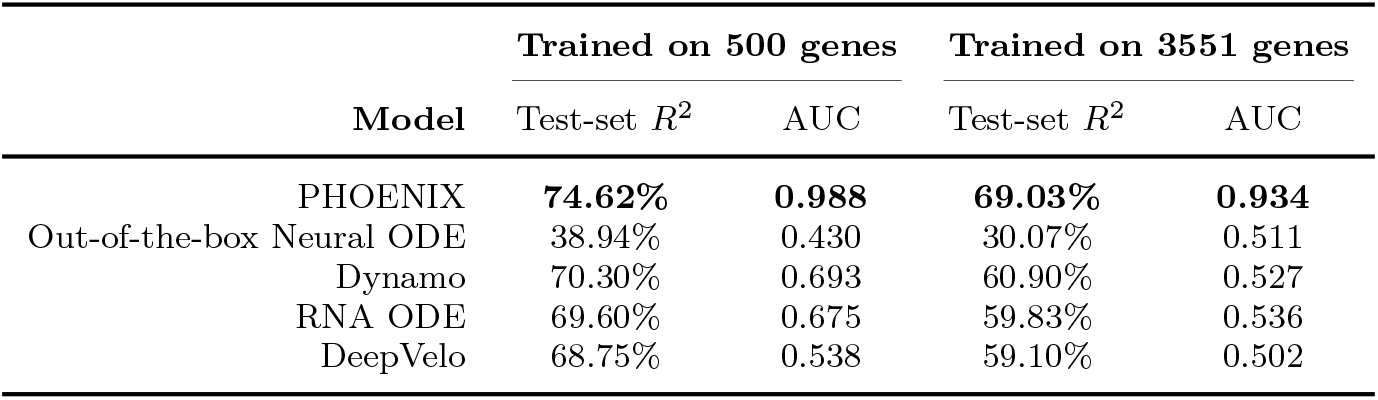
Comparison of PHOENIX with competitor methods on yeast data with all 3551 genes and separately, the 500 most variable genes. Snapshot-based methods (Dynamo, RNA-ODE, Deepvelo) require RNA velocity at every time point as an additional input [3, 5, 12]. Given that this information was not available in the data set, we estimated RNA velocity using a method of finite differences applied to smooth splines through the expression trajectories [40] (see Supp. Methods 3.1). Predictive performance is reported as the *R*^2^ on the test set. Explainabilty AUC was calculated by comparing encoded GRNs retrieved from each trained model (Supp. Methods 2 and Supp. Methods 3.3) against ChIP-chip data [39].

To test the biological explainability of the learned dynamical system, we extracted the encoded GRN from the trained PHOENIX model (with optimal *λ*_prior_ = 0.05 as determined by the validation set) and compared it to a validation network of ChIP-chip transcription factor (TF) binding data [39]. PHOENIX had very impressive accuracy in predicting TF binding (AUC = 0.93), indicating that it had learned transcription factor binding information in the process of explaining temporal patterns in expression (**Figure 4C**). In the absence of any prior knowledge (*λ*_prior_ = 0), the explainability was poor, highlighting the importance of such knowledge-based guidance in black-box models [24, 25], as well as the importance of correctly tuning *λ*_prior_.

Similar to the *in silico* experiments, we saw that PHOENIX’s ability to predict TF binding was greater than that obtained by comparing just the prior to the validation data (Supp. Table 1). This suggested that PHOENIX had used the prior knowledge of cell cycle progression as a starting point to anchor the dynamics, and then used the data itself to learn improved regulatory rules.

In order to contextualize PHOENIX’s impressive performance on this data set, we performed comparative analyses against Dynamo [3], RNA-ODE [5], DeepVelo [12], and out-of-the-box NeuralODEs. As discussed in Supp. Table 7, Dynamo, RNA-ODE, and DeepVelo are “two-step” snapshot based methods that require RNA velocity at every time point as an additional input [3, 5, 12]. Given that this information was not available in the data set, we estimated RNA velocity using a method of finite differences applied to smooth splines through the expression trajectories [40] (see Supp. Methods 3.1). Furthermore, given that it is a common approach for current methods to only consider a subset of top-*k* highly variable genes [4, 6, 7, 12, 14], we additonally performed a subsetted analysis considering only the 500 most variable genes along the trajectory.

In both sets of analysis, we found PHOENIX to be both the most predictive *R*^2^ *∈* (69%, 75%) and by far the most explainable in terms of AUC *∈* (0.93, 0.99) (**Table 1**). Dynamo, RNA-ODE, and DeepVelo were designed for a predictive setting, and we see that they do perform well on the sub-setted analysis when looking at predictive performance, albeit worse than PHOENIX. In terms of reconstruction of the underlying GRN, they however perform poorly considering the AUC, with a difference of about 0.30 or more to PHOENIX. When taken in conjunction with the poor predictive performance of out-of-the-box NeuralODEs (*R*^2^ *∈* [30%, 39%]), as well as the poor explainability of all black-box methods, this highlights the unique strength of PHOENIX to appropriately model important biological processes, such as cell cycle progression, while being highly predictive on timecourse data.

### PHOENIX infers genome-wide dynamics of breast cancer progression and identifies central pathways

Although there are several tools for inferring the dynamics of regulatory networks, most do not scale beyond a few hundreds of genes without losing explainability, falling far short of the 25,000 genes in the human genome (Supp. Table 7). Given the performance improvements we saw that were driven by PHOENIX’s use of soft constraints, we wanted to test whether PHOENIX could be extended to human-genome scale networks. Due to the dearth of longitudinal human studies with genome-wide expression measurements, we used data from a cross-sectional breast cancer study (GEO accession GSE7390 [41]) consisting of microarray expression values for 22000 genes from 198 breast cancer patients and ordered these samples in pseudotime. For consistency in pseudotime ordering, we reused a version of this data that was already preprocessed and ordered (using a random-walk-based pseudotime approach) in the PROB paper [8].

After further processing (Methods 3) obtained a single pseudotrajectory of expression values for *n*_*g*_ = 11165 genes across 186 patients, each at a distinct pseudotimepoint. To explore whether PHOENIX’s performance depends on the size of the data set, we also created pseudotrajectories for *n*_*g*_ = 500, 2000, and 4000 genes by subsetting the data set to its *n*_*g*_ most variable genes. We split up the 186 time points into contiguous intervals for training (170, 90%), validation (8, 5%), and test (8, 5%). For the domain prior network, we again used a simplistic prior model derived from a motif map of promoter targets, and tuned *λ*_prior_ using the validation set. See Methods 3.2 for further details.

For each pseudotrajectory of size *n*_*g*_, we trained a separate PHOENIX model, and measured predictive performance as the variation explained (*R*^2^) in the test trajectory by predicting the trajectory through the model from just the initial time point (see Methods 3.2). We observed encouragingly high values of *R*^2^ *∈* (92%, 97%) (**Figure 5**, top), that are, however, not what to expect given this is frozen tumor tissue. Upon further investigation we found that the high *R*^2^ was partly driven by most genes not showing enough dynamic expression, which hence are easily predicted with a constant trajectory. When we instead focused on evaluating only the top most variable genes, we observed more reasonable values of *R*^2^ (Supp. Fig. 10, blue line). We also considered alternate strategies of measuring predictive performance in this setting, but found this to oversimplify the task (Supp. Fig. 10, red line). We note here that PHOENIX’s computational cost was not excessive even when *n*_*g*_ = 11165 (see Supp. Table 10).

**Figure 5.**
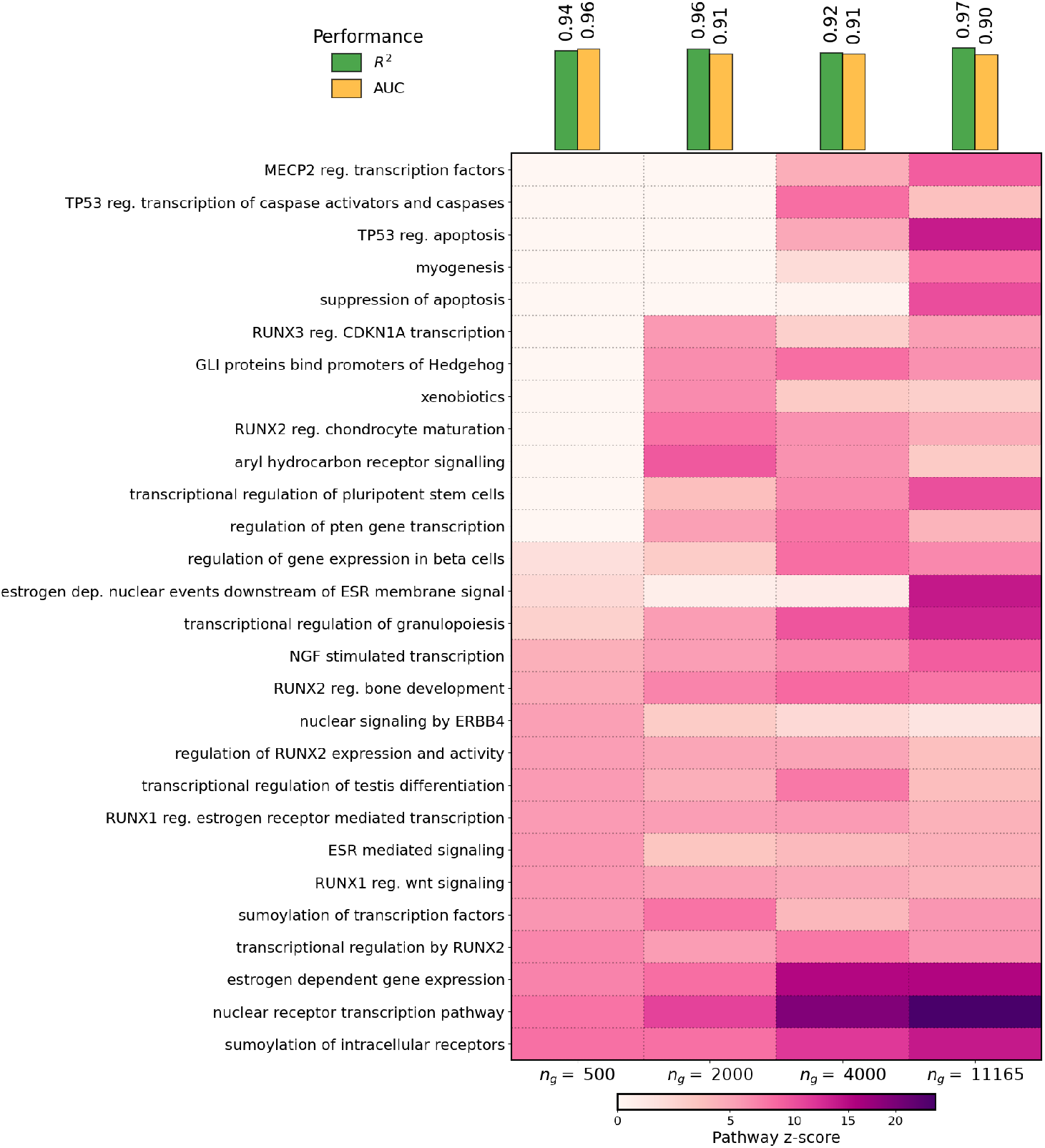
We applied PHOENIX to a pseudotrajectory of 186 breast cancer samples (ordered along subsequent “pseudotimepoints”) consisting of *n*_*g*_ = 11165 genes [41]. We split up the 186 time points into contiguous intevals for training (170, 90%), validation to tune *λ*_prior_(8, 5%), and test (8, 5%). We also repeated the analysis on smaller subsets of genes *n*_*g*_ = 500, 2000, 4000, where we subsetted the full trajectory to only the *n*_*g*_ most variable genes in the pseudotrajectory. We measured predictive performance as the *R*^2^ on the test trajectory by applying the trained model to just the first test time point (see Methods 3.2). We evaluated explainability performance as the AUC from comparing encoded GRNs from trained models against a ChIP-seq validation network [42]. Finally, we used the trained PHOENIX models to extract permutation-based influence scores for pathways in the Reactome database [52] (see Methods 3.3 and 3.4), and visualized influence scores for a collection of the most central pathways. See Supp. Table 8 and Supp. Table 9 for detailed results.

Next, we investigated PHOENIX’s ability to identify biologically relevant and actionable information regarding gene regulation in breast cancer. First, we tested the performance of the learned dynamical system to reconstruct a gene regulatory network and predict TF-gene interactions. While the ground truth GRN is unknown, we can estimate performance by comparing a validation network of experimental ChIP-chip binding information [42] to a subnetwork of the encoded GRN of a trained PHOENIX model. We found excellent alignment between the two GRNs with AUC *∈* (0.90, 0.96), even when we scaled up to *n*_*g*_ = 11165 genes (**Figure 5**). It is important to note that the PHOENIX-based concordance with experimental data was much greater than that obtained by comparing just the prior knowledge to the validation network (Supp. Table 1), indicating that PHOENIX was improving upon the GRN suggested by the prior knowledge, in addition to learning a dynamical model.

To better understand the benefits of PHOENIX’s scalability, we investigated how estimating regulatory dynamics based on a subset of only the *n*_*g*_ most variable genes can alter the perceived importance of individual genes to the regulatory system in question. We reasoned that a model trained on all assayed genes should reconstruct biological information better than those that are restricted to a subset of genes [6, 12, 14]. First, we performed a gene-level analysis by perturbing *in silico* the PHOENIX-estimated dynamical system from each value of *n*_*g*_ (500, 2000, 4000, 11165). This yielded “influence scores” representing how changes in initial (*t* = 0) expression of each gene affected subsequent (*t >* 0) predicted expression of all other genes (see Methods 3.3). As might be expected, the influence scores grew increasingly more concordant with centrality measures in the ChIP validation network, consistent with the key roles played by transcription factor genes in large GRNs (Supp. Table 10).

We observed that highly variable genes with known involvement in breast cancer (such as WT1 [43], ESR1 [44], AR [45], and FOXM1 [46]) were generally influential across all values of *n*_*g*_ (Supp. Fig. 7). It is interesting to note that both FOXM1 and AR were very influential in the *n*_*g*_ = 500 system, but their score dropped in the full genome (*n*_*g*_ = 11165) system. This is likely due to the way in which we constructed the smaller subsets of the whole genome – by selecting the most variable genes. One would expect that the most variable transcription factor genes falling within any subset would be highly correlated in expression with other genes falling in the same set and that the overall effect would be diluted by adding more - potentially uncorrelated - genes to the system. It is more interesting that genes missing in the smaller subsets (due to low expression variability) were identified as central to the dynamics in the full (*n*_*g*_ = 11165) system. Among these genes, we can find some encoding cancer-relevant transcription factors such as E2F1 [47, 48] CTCF [49], and ERG [50], and DNA methyltransferase enzymes (DNMT1 [51]).

We found that the more computationally manageable systems (*n*_*g*_ = 500, *n*_*g*_ = 2000) yielded an incomplete picture of gene-level influences since the method used in constructing these subsets hinders the mechanistic explainability of the resulting regulatory model. Certain genes exhibit relatively low variability in expression but are still central to disease-relevant genome-level dynamics; compared to methods that exclude such genes to make computation tractable [4, 6, 12], PHOENIX can correctly identify such as central because of its ability to model subtle but important genome-scale dynamics.

Finally, we performed a pathway-based functional enrichment analysis by translating these gene influence scores to pathway influence scores using permutation tests on the Reactome pathway database [52] (see Methods 3.4). We reasoned that a more complete network, to have practical advantages over smaller and more manageable models, should be able to capture a more complete picture of biological processes that are involved in the cancer pseudotime. Not surprisingly, the dynamical systems with fewer genes missed many pathways known to be associated with breast cancer that were identified as over-represented in the genome-scale (*n*_*g*_ = 11165) system (**Figure 5** and Supp. Table 8). Notably, the pathways missed in the smaller networks include apoptosis regulation (a hallmark of cancer [53]), and TP53 regulation of caspases (relevant to apoptosis control in tumors [54]), while terms for the estrogen-related signalling and GLI/Hedgehog signalling (whose role in cancer is well documented [55, 56]) would have been missed or underestimated by the smaller models.

In a parallel analysis testing for functional enrichment of GO biological process terms, we again found the smaller systems to overlook important pathways that were clearly influential in the genome-scale analysis; these included a wide array of RNA metabolism processes that are increasingly recognized as being significant to breast cancer development [57] (Supp. Fig. 8). Finally, while the GO molecular function terms are consistently dominated by binding terms, we can notice how the larger models are capable of detecting more specific terms such as BHLH TF binding and E-box binding, that are subgroups of the more TF binding term, that are known for the regulation of well known cancer-related genes such as NOTCH1 and MYC [58, 59] (Supp. Fig. 9).

These results clearly demonstrate the importance of scalable methods such as PHOENIX that can **explainably** model genome-wide dynamics. Our reduced gene sets from which we built the smaller PHOENIX models consisted of the 500, 2000, or 4000 most variable genes. These gene sets likely consist of variable genes that are correlated with each other, meaning that we are sampling only a portion of the biological processes driving the temporal changes in breast cancer; the full picture only emerges when looking at regulatory processes across the spectrum of genes that can contribute. Alternative approaches, such as concentrating on specific pathways, risk introducing self-fulfilling biases in the discovery process. Similarly, methods that use low-dimensional embedding (PCA, UMAP, etc.) to reduce the complexity of modeling dynamics risk losing valuable, biologically relevant insights. PHOENIX’s ability to scale while remaining explainable offers the best potential for the discovery of interpretable insights about the phenotypes under study.

In order to test whether the ability to remain explainable while scaling genome-wide is unique to PHOENIX, we performed comparative analyses on the full set of *n*_*g*_ = 11165 genes against Dynamo [3], RNA-ODE [5], Deep-Velo [12], and out-of-the-box NeuralODEs. Analogous to the yeast example, RNA velocity information was unavailable, and hence estimated using finite differences on smooth splines[40] for the “two-step” snapshot based methods [3, 5, 12] (see Supp. Methods 3.1). In order to better understand how well a model trained at genome-level is still able to capture biologically meaningful nuances all across the gene-network, we obtained the predicted 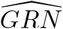 (each with *n*_*g*_ = 11165 nodes) from each trained model (Supp. Methods 2, Supp. Methods 3.3), but measured explainability in an incremental manner. For *N* ranging from 50 to 11165, we calculated the concordance between the induced subgraph of each 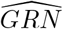 spanned by the *N* -most variable genes against the corresponding ChIP-seq validation subnetwork [42].

The results indicated that PHOENIX is on par with the existing black-box approaches in terms of test set predictive accuracy (Supp. Table 11) while offering interpretable, biologically meaningful results. This is reflected in the achieved AUC (Figure 6), where only PHOENIX is able to properly reconstruct the underlying GRN. Most black-box methods fulfil their goal of being highly predictive without offering much insight into the (interpretable) underlying processes. Notably, RNA-ODE recovers a GRN when considering only the top 100 most variable genes, which is, however, arguably not useful in practice. PHOENIX was the only approach that both scales to the full set of genes, being highly predictive, while being interpretable.

**Figure 6.**
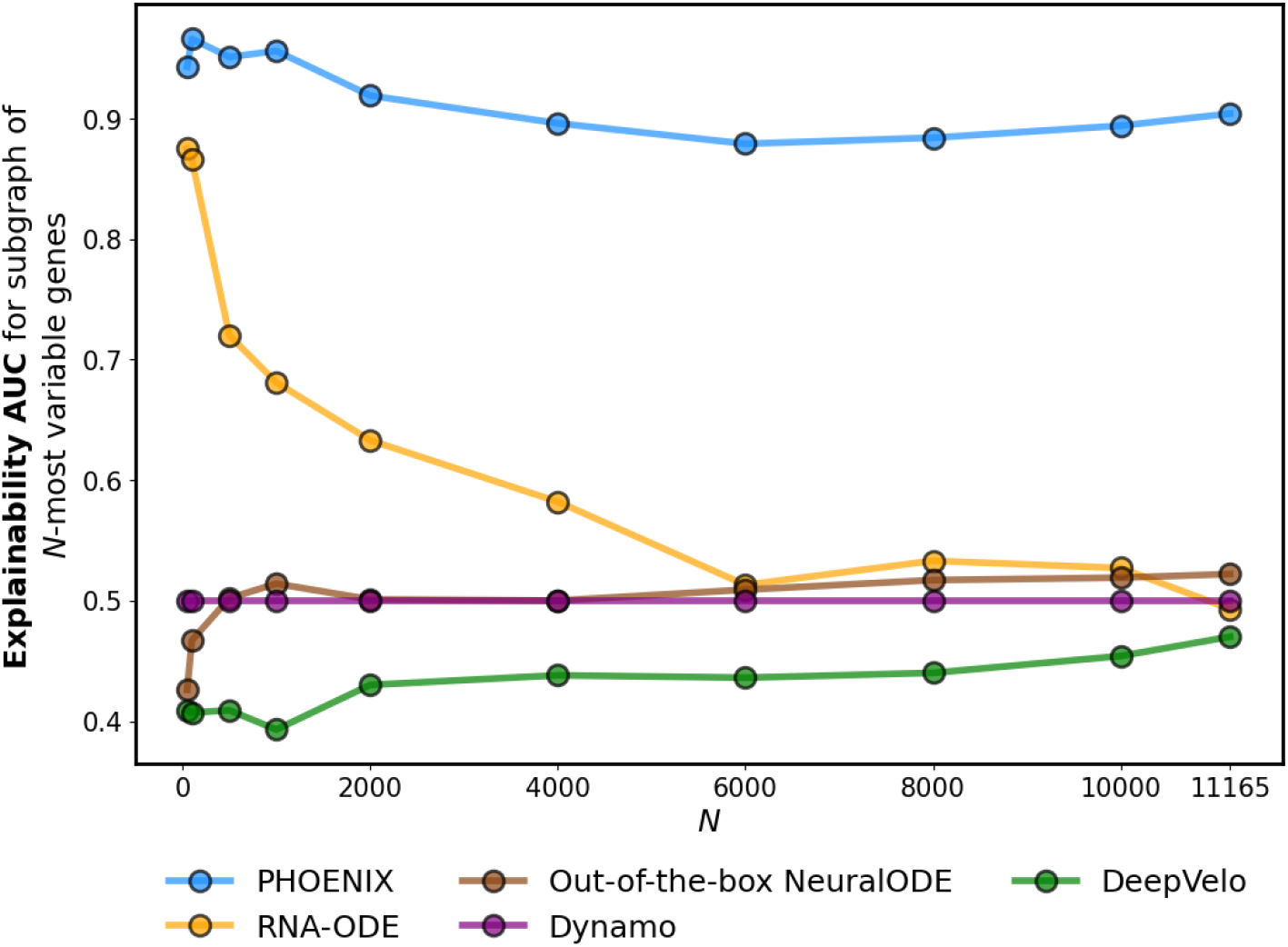
We compared the performance of PHOENIX to other methods of regulatory dynamics estimation on a pseudotrajectory of 186 breast cancer samples (ordered along subsequent “pseudotimepoints”) consisting of *n*_*g*_ = 11165 genes [41]. The data was processed for model fitting via the steps described in Methods 3.2. Snapshot-based methods (Dynamo, RNA-ODE, Deepvelo) require RNA velocity at every time point as an additional input [3, 5, 12]. Given that this information was not available in the data set, we estimated RNA velocity using a method of finite differences applied to smooth splines through the expression trajectories [40] (see Supp. Methods 3.1). Once each model was trained, we sought to measure explainability by obtaining the predicted GRN from it (Supp. Methods 2, Supp. Methods 3.3). Then, for *N* ranging from 50 to 11165, we calculated the concordance between the subgraph of each 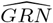 spanned by the top-*N* most variable genes in the data set against the corresponding ChIP-seq validation subnetwork [42]

### PHOENIX recovers key gene regulatory changes in Rituximab treated B cells

To challenge PHOENIX with a different data modality, we considered a longitudinal RNA-seq experiment of B-cells followed over a course of 15 hours, where cells where either treated with Rituximab or kept untreated [60]. Rituximab is used as treatment for specific leukemia, non-hodgkin lymphoma as well as rheumatoid arthritis, binding to B-cells inducing cell death through apoptosis, NK-cell mediated cytotoxicity, or Macrophage mediated phagocytosis. Both treated and untreated B-cells were sampled at *t* = 0, 1, 2, 4, 7, and 15 hours and two replicates were available for analysis.

We trained two separate PHOENIX models, one each for the treated and untreated cells, respectively. This allowed us to compare the gene regulatory changes after Rituximab treatment by examining the differences between the gene regulatory networks encoded by the two trained PHOENIX models. We split the time points in each condition into training (80%) and validation (20%). For the domain prior network, we use a TF-binding derived prior similar to the breast cancer study above (see Methods 4.2), and tuned *λ*_prior_ using the validation set. We observed that PHOENIX was able to model gene expression dynamics in both conditions well, with *R*^2^ values between 89% and 92% (Supp. Table 13). We extracted the two encoded GRNs (one each from the PHOENIX model trained on the two conditions), and subsequently sought to examine the changes of the regulatory dynamics that were visible between untreated B cells and those treated with Rituximab.

Aggregating the regulatory effects of a protein (input) across all (output) genes, we then compute log-fold changes of these regulatory potentials between the two GRNs. A closer look at the top 50 regulators in terms of changes in regulatory dynamics (Supp. Table 12), we find several key regulators of apoptosis, the main mechanism of action of Rituximab in this experiment, as neither macrophages nor NK cells are present to enable NK-mediated cytotoxicity or M-mediated phagocytosis. Among others, we find PPP2R3C, a key inhibitor of B-cell receptor induced apoptosis [61], PHLPP2, which acts by dephosphorylation of AKT family members on apoptosis [62], RAB1A, which is part of the RAS signaling cascade that acts anti-apoptotic [63], and CLUH that regulates the mTORC1 signaling pathway and hence apoptosis [64]. All of these show a more than 2x log-fold change in terms of regulatory dynamics between the two conditions. Examining the two main pathways of action of Rituximab, Apoptosis and B-cell receptor signaling, we further observe a generally strong regulatory difference of genes annotated for those pathways between treatment and control (see Supp. Fig. 11 and Supp. Fig. 12). Our analysis provides further evidence that PHOENIX reflects meaningful biological signals also on RNA-seq data, even when only few unevenly sampled measurements are available.

## Discussion

Given the importance of regulatory networks and their dynamics, there has been a tremendous interest in inferring and modeling their physical and temporal behavior. The use of NeuralODEs represents an extremely promising technology for inferring such networks, but so far, attempts to implement NeuralODE-based network modeling have encountered significant problems, not the least of which has been their inability to scale to modeling genome-wide dynamics in a biologically explainable manner.

PHOENIX represents an important new methodological extension to the NeuralODE framework that is not only scaleable to the full human genome but also biologically well interpretable and able to capture explicitly both additive as well as multiplicative ways in which transcription factors cooperate in regulating gene expression. For a simplified analysis, the underlying gene regulatory network can also be extracted from a learned model and compared with experimental evidence. An optional feature of PHOENIX that contributes significantly to its explainability is that it can be guided by (structural) domain knowledge. Notably, PHOENIX also remains flexible to deviate from domain knowledge when necessary and learn novel insights consistent with the training data.

The predictive accuracy, scalability, flexibility, and biological explainability can be attributed primarily to two things. First, our novel NeuralODE architecture includes the use of Hill-like activation functions for capturing the kinetic properties of molecular binding provides a massive advantage in terms of predictive power. Second, the introduction of soft constraints based on prior knowledge of putative network structure leads to a scalable and biologically explainable estimate of the underlying dynamics.

Using simulated data we have shown that PHOENIX outperforms other models for inferring regulatory dynamics (including other NeuralODE-based models), particularly in the presence of experimental noise. Also, an application to data from the yeast cell cycle elucidates PHOENIX’s flexibility in modeling arbitrary dynamics. More importantly, PHOENIX is the only NeuralODE method capable of extending its modeling to capture genome-scale regulatory processes, while remaining explainable. Using data from breast cancer patients organized in pseudotime we illustrate not only the ability of PHOENIX to faithfully model genome-scale networks but also demonstrate the power of extending regulatory modeling to capture seemingly subtle but biologically important regulatory processes.

One of the challenges in modeling the evolution of network processes, of course, is obtaining data sets for which temporal data are available. However, we recognize that in any data set, the individual samples represent a continuum of states between health and disease and so can use pseudotemporal ordered samples. But this too has some limitations as there is as of yet no established method for ordering bulk samples – although there have been some methods for single-cell data adapted to “bulk” tissue samples [65]. This admits an interesting possibility: one could use additional information (such as RNA velocities, provided they are reliable) from which one could infer pseudotime (or real-time) trajectories. In that sense, one may argue that PHOENIX provides a general approach to infer interpretable GRNs and ODE models from outputs of other methods (such as Dynamo [3]), which are currently less transparent. This could provide the best of both worlds with a reduced dimensional approach to temporal ordering providing input for a more complete and interpretable final model.

Although PHOENIX, in its current implementation, is designed with one “layer” of regulation to model TF-gene interactions, we recognize that there are other regulatory elements or higher order regulatory effects in the cell that contribute to the control of gene expression. Such additional effects could potentially be modeled by increasing the complexity of the NeuralODE solver by introducing additional layers to the NeuralODE framework. However, this would increase the computational requirements by PHOENIX and reduce its current advantage of being a light-weight method.

## Methods

### 1 Testing on simulated data

#### 1.1 Defining a ground truth dynamical system

We created a ground truth gene regulatory network (GRN) by sampling from *S. cerevisiae* (yeast) regulatory networks obtained from the SynTReN v1.2 supplementary data in simple interaction format (SIF) [66]. The SynTReN file provides a directional GRN containing 690 genes and 1094 edges with annotations (activating vs repressive) for edge types; we defined this GRN to be a ground-truth network *G*_690_. To obtain *G*_350_, we sampled a subnetwork of 350 genes and 590 edges from *G*_690_. We used the connectivity structure of *G*_350_ and *G*_690_, to define systems of ODEs (SIM350 and SIM690) with randomly assigned coefficients. This entire pipeline was executed using SimulatorGRN [35], a framework used extensively by the R/Bioconductor package dcanr [29]. Please see Supp. Methods 4.1 for further ODE formulation details.

#### 1.2 Simulating time series gene expression data

For each *n ∈ {*350, 690*}*, we used the ground truth dynamical system SIM*n* to generate expression vectors ***g***(*t*) *∈* R^*n*^, across time points *t*. We started by i.i.d. sampling 160 standard uniform R^*n*^ vectors to act as initial (*t* = 0) conditions. We used these initial conditions to integrate SIM*n* and obtain 160 expression trajectories across *t ∈ T* = *{*0, 2, 3, 7, 9*}* using R’s desolve package: 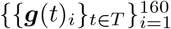. We used only five time points to emulate potential scarcities of time-series information in real data sets, while the range *t* = 0 to 9 generally covered the transition from initial to steady state. Lastly, we added Gaussian noise vectors 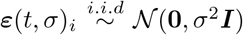 of varying *σ* to get noisy data sets: 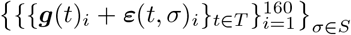. Since the average simulated expression value was *≈* 0.5, using 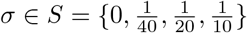 corresponded roughly to average noise levels of 0%, 5%, 10%, 20%.

#### 1.3 Model setup for training and testing

For each simulation scenario, there were 160 simulated trajectories, out of which we used 140 (88%) for training, 10 (6%) for validation (hyperparameter tuning), and 10 (6%) for testing. We provide some details on PHOENIX implementation (such as training strategy, prior incorporation, etc.) in Supp. Methods 1, and include finer technicalities (including learning rates) in our GitHub repository [36]. For prior domain knowledge model, we used the simple linear model: 𝒫^*∗*^ (***γ***_***k***_) = ***A***^***σ*%**^. ***γ***_***k***_ *−* ***γ***_***k***_, where we chose ***A***^***σ*%**^ to be noisy/corrupted versions of the adjacency matrices of ground truth networks *G*_350_ and *G*_690_ (details in Supp. Methods 4.2). We set activating edges in ***A***^***σ*%**^ to +1 and repressive edges to −1. “No interaction” was represented using 0. To validate explainability we extracted GRNs from trained models, and compared to ground truth *G*_350_ and *G*_690_ for the existence of edges, out-degree correlations, and induced sparsity (details in Supp. Methods 2).

### 2 Testing on experimental yeast cell cycle data

#### 2.1 Data processing and normalization

GPR files were downloaded from the Gene Expression Omnibus (accession **GSE4987** [37]), and consisted of two dye-swap technical replicates measured every five minutes for 120 minutes. Each of two replicates were separately ma-normalized using the maNorm() function in the marray library in R/Bioconductor [67]. The data were batch-corrected [68] using the ComBat() function in the sva library [69] and probe-sets mapping to the same gene were averaged, resulting in expression values for 5088 genes across fifty conditions. Two samples (corresponding to the 105 minute time point) were excluded for data-quality reasons, as noted in the original publication, and genes without motif information were then removed, resulting in an expression data set containing 48 samples (24 time points in each replicate) and 3551 genes.

#### 2.2 Model setup for training and testing

Given that the data set contained two technical replicates of gene expression trajectories, we used one replicate for training and the other for testing (not seen in any way during training). Furthermore, within the replicate for training, we used a the expression values of contiguous segment of time points *t* = 45, 50, 55 minutes for validation (to choose *λ*_prior_). The two remaining intervals *t ∈* [0, 40] and *t ∈* [60, 120] in this replicate were used for training. This scheme resulted in train (87%) and validation (13%) sets that were disjoint, and crucially, a test set that was independent of the training data. We provide more details on PHOENIX implementation (such as training strategy, prior incorporation, etc.) in Supp. Methods 1, and include finer technicalities (including learning rate schedule) in our GitHub repository [36].

For prior domain knowledge model we used the simple linear model: 𝒫^*∗*^ (***γ***_***k***_) = ***A***. ***γ***_***k***_ *−* ***γ***_***k***_. We based our choice of ***A*** on the regulatory network structure of a motif map, similar to that used in other methods, such as PANDA [26]. We downloaded predicted binding sites for 204 yeast transcription factors (TFs) [39]. These data include 4360 genes with tandem promoters. 3551 of these genes are also covered on the yeast cell cycle gene expression array. 105 total TFs in this data set target the promoter of one of these 3551 genes. The motif map between these 105 TFs and 3551 target genes provides the adjacency matrix ***A*** of 0s and 1s, representing whether or not a prior interaction is likely between TF and gene.

We used ChIP-chip data from Harbison et al. [39] to create a network of TF-target interactions, and used this as a validation network to test explainability. The targets of transcription factors in this ChIP-chip data set were filtered using the criterion *p <* 0.001. We calculated AUC values by comparing the encoded GRN retrieved from the trained models (see Supp. Methods 2) to the validation network.

### 3 Testing on breast cancer pseudotime data

#### 3.1 Data procurement and psuedotime ordering

The original data set comes from a cross-sectional breast cancer study (GEO accession **GSE7390** [41]) consisting of microarray expression values for 22000 genes from 198 breast cancer patients, that we sorted along a pseudotime axis. We noted that the same data set was also used in the PROB [8] paper. PROB is a GRN inference method that infers a random-walk-based pseudotime to sort cross-sectional samples and reconstruct the GRN. For consistency and convenience in pseudotime inference, we obtained the same version of this data that was already preprocessed and sorted by PROB. We normalized the expression values to be between 0 and 1. We limited our analysis to the genes that had measurable expression and appeared in our prior model and obtained a pseudotrajectory of expression values for 11165 genes across 186 patients. We also created pseudotrajectories for *n*_*g*_ = 500, 2000, and 4000 genes by subsetting to the *n*_*g*_ highest variance genes.

#### 3.2 Model setup for training and testing

We noted that the processed data set contained expression across 186 “pseudo” time points. We excised a contiguous interval of expression across 8 time points for testing (5%), and split up the remaining 178 time points into training (170, 90%) and validation for tuning *λ*_prior_ (8, 5%). To measure predictive accuracy of the trained model, we used it calculated a predicted trajectory based on just the first time point in the test set. We calculated the *R*^2^ between this prediction and the remaining 7 points of the test trajectory. Further implementation details in Supp. Methods 1 and GitHub [36].

For the prior domain knowledge model, we used the simple linear model: 𝒫^*∗*^ (***γ***_***k***_) = ***W***_**0**_ ***γ***_***k***_ *−* ***γ***_***k***_. We based our choice of ***W***_**0**_ on a motif map, similar to that used in the breast cancer analysis in OTTER [70]. The network ***W***_**0**_ is derived from the human reference genome, for the breast tissue specifically. *W*_0_ is a binary matrix with 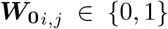 where 1 indicates a TF sequence motif in the promoter of the target gene. Sequence motif mapping was performed using the FIMO software [31] from the MEME suite [71] and the R package GenomicRanges [32].

Validation of explainability was challenging since there are only a few data sets that have ChIP-seq data for many TFs from the same cells. We used ChIP-seq data from the MCF7 cell line (breast cancer, 62 TFs) in the ReMap2018 database [42] to create a validation network of TF-target interactions. We calculated AUC values by comparing the encoded GRNs retrieved from the trained models (see Supp. Methods 2) to the validation network.

#### 3.3 Gene influence scores

Given 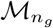 a PHOENIX model trained on the pseudotrajectory consisting of only the *n*_*g*_ most variable genes (*n*_*g*_ ∈ {500, 2000, 4000, 11165}), we performed perturbation analyses to compute gene influence scores 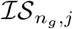. We randomly generated 200 initial (*t* = 0) expression vectors via i.i.d standard uniform sampling 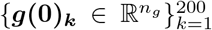. Next, for each gene *j* in 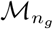, we created a perturbed version of these initial value vectors 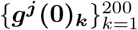, where only gene *j* was perturbed in each unperturbed vector of 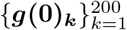. We then fed both sets of initial values into 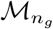 to obtain two sets of predicted trajectories 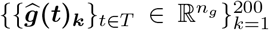 and 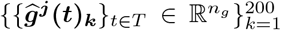 across a set of time points *T*. We calculated influence as the average absolute difference between the two sets of predictions, that represented how changes in initial (*t* = 0) expression of gene *j* affected subsequent (*t >* 0) predicted expression of all other genes in the *n*_*g*_-dimensional system:

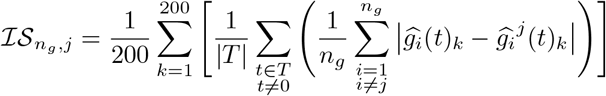

#### 3.4 Pathway influence scores

Having computed gene influence scores 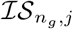 for each gene *j* in each dynamical system of dimension *n*_*g*_ genes, we translated these gene influence scores into pathway influence scores. We used the Reactome pathway data set, GO biological process terms, and GO molecular function terms from MSigDB [72], that map each biological pathway/process, to the genes that are involved in it. For each system of size *n*_*g*_, we obtained the pathway (*p*) influence scores 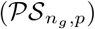 as the sum of the influence scores of all genes involved in that pathway:

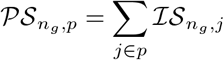

We statistically tested whether each pathway influence score is higher than expected by chance using empirical null distributions. We randomly permuted the gene influence scores across the genes to recompute “null” values 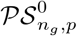. For each pathway, we performed *K* = 1000 permutations to obtain a null distribution 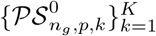 that can be compared to 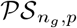. We could then compute an empirical *p*-value as 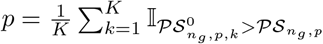, where I is the indicator function. Finally, we used the mean 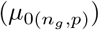 and variance 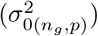 of the null distribution 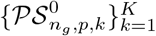 to obtain and visualize pathway *z*-scores that are comparable across pathways and subset sizes (*n*_*g*_):

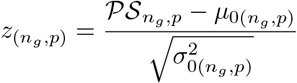

### 4 Testing on B-cell RNASeq data

#### 4.1 Data procurement and processing

RNA-seq data of B-cells has been downloaded from the Gene expression omnibus (GEO accession **GSE100441** [60]). Filtering out genes that show zero expression in all samples and genes that are not in the prior, we are left with *n* = 14691 genes with log(*FPKM* + 1) transformed gene expression values. We use the same prior as in the breast cancer experiment described above. The data spans across seven time points (t=0*h*, 1*h*, 2*h*, 4*h*, 7*h*, 15*h* after treatment start) for B cells treated with Rituximab as well as parallel untreated B cells as control. Two replicates are available for each condition.

#### 4.2 Model setup for training and GRN extraction

Given that the data set for each condition consists of 2 technical replicates of 6 time points each, we note that it contains 10 different transition pairs, where a transition pair consists of two consecutive expression vectors in the data set (***g***(*t*_*i*_), ***g***(*t*_*i*+1_)). We randomly split these 10 transition pairs into training (8, 80%) and validation (2, 20%), where the validation set was used to tune *λ*_prior_ as well as inform an early stopping criteria. Further details on the implementation are in Supp. Methods 1 and our GitHub repository [36]. For the prior domain knowledge model, we used the same motif-based prior as in our breast cancer analysis (Methods 3.2), 𝒫^*∗*^ (***γ***_***k***_) = ***W***_**0**_. ***γ***_***k***_ *−* ***γ***_***k***_.

#### 4.3 Rituximab pathway analysis

To examine whether PHOENIX recovers meaningful biology in this challenging dataset, we focused on analysing the differences between the derived GRNs of the Rituximab-treated and control group. Focusing on changes of regulators, we first aggregate the influence score *s*_*i*_ of each regulator *i* by summing the weights of outgoing edges, 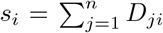, a common approach in node-centric analysis in GRNs. Note that our dynamics matrix D incorporates the scaling factor but is not normalized per gene, to enable a comparison between the dynamics of two networks (here: treated and untreated). To examine the change of regulatory influence of each protein, we then compute the log-fold change between influence scores in the two conditions as

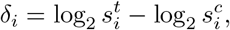

where 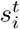 and 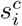 are the influence score in the GRN of the treatment respectively control group. As the primary mechanism of action of Rituximab is known—directly induced apoptosis, complement-dependent cytotoxicity, NK-mediated cytotoxicity, and macrophage-mediated phagocytosis—we focused on analzying the molecular changes within these mechanisms, with a particular focus on the top 50 most changing regulators (see Supp. Table 12). As the experiment contained only B-cells, hence NK and macrophages were not available for cell destruction, we focused on apoptosis and cytotoxicity related to B cell receptor signaling. We provide pathway maps of these pathways colored by (normalized) *δ*_*i*_ in Supp. Fig. 11 and Supp. Fig. 12, which were visualized using the Pathview package (version 1.42.0) [73] in R (version 4.3.2).

## Supporting information

Supplemental Results & Methods

## Supplementary information

The article is accompanied by an Online Supplementary Information file containing additional results and further details about the methodology. Also, all relevant code and data is available as open source with the PHOENIX release: https://github.com/QuackenbushLab/phoenix, and via Zenodo (10.5281/zenodo.10412968).

## Acknowledgments

The authors thank Marouen Ben Guebila, Dawn DeMeo, Kimberly Glass, Camila Lopes-Ramos, Panagiotis Mandros, Soel Micheletti, Enakshi Saha, and Katherine H. Shutta for thoughtful critiques and discussions. We also thank Daniel Karlsson and Olle Svanström for sharing with us their starter code [74] for basic NeuralODE training.

## Declarations

The authors declare the following:

## Availability of data and materials

All open-source datasets used in the paper have been referenced with their GEO accession ID. Furthermore, relevant data files have also been deposited on Zenodo (10.5281/zenodo.10412968). The source code is available as an open-source implementation (distributed under license **CC BY-NC 4.0**) via Github: https://github.com/QuackenbushLab/phoenix [36], and has also been deposited on Zenodo (10.5281/zenodo.10412968).

## Authors’ contributions

IH, JQ, and RB conceived of the project. RB and IH designed the modeling framework. IH carried out the bulk of the work including data processing and coding of the methodology. VF contributed to the breast cancer pathways modeling and interpretation. JF contributed the B cell modeling and interpretation. All authors contributed to the writing and editing of the manuscript.

## References

[1] Xing, J. (2022). Reconstructing data-driven governing equations for cell phenotypic transitions: integration of data science and systems biology. Physical Biology, 19(6), 061001.

[2] Hackett, S. R., Baltz, E. A., Coram, M., Wranik, B. J., Kim, G., Baker, A., … & McIsaac, R. S. (2020). Learning causal networks using inducible transcription factors and transcriptome-wide time series. Molecular systems biology, 16(3), e9174.

[3] Qiu, X., Zhang, Y., Martin-Rufino, J. D., Weng, C., Hosseinzadeh, S., Yang, D., … & Weissman, J. S. (2022). Mapping transcriptomic vector fields of single cells. Cell, 185(4), 690–711.

[4] Yeo, G. H. T., Saksena, S. D., & Gifford, D. K. (2021). Generative modeling of single-cell time series with PRESCIENT enables prediction of cell trajectories with interventions. Nature communications, 12(1), 1–12.

[5] Olteanu, M., & Stefan, R. (2020). An exponential stability test for a messenger rna–micro rna ode model. University politehnica of bucharest scientific bulletin-series a-applied mathematics and physics, 82(4), 11–16.

[6] Erbe, R., Stein-O’Brien, G., & Fertig, E. J. (2023). Transcriptomic forecasting with neural ordinary differential equations. Patterns (New York, N.Y.), 4(8), 100793. 10.1016/j.patter.2023.100793

[7] Li, Q. (2022). scTour: a deep learning architecture for robust inference and accurate prediction of cellular dynamics. bioRxiv, 2022–04

[8] Sun, X., Zhang, J., & Nie, Q. (2021). Inferring latent temporal progression and regulatory networks from cross-sectional transcriptomic data of cancer samples. PLoS computational biology, 17(3), e1008379.

[9] Mendes, P., Hoops, S., Sahle, S., Gauges, R., Dada, J., & Kummer, U. (2009). Computational modeling of biochemical networks using COPASI. Systems Biology, 17–59.

[10] Kraeutler, M. J., Soltis, A. R., & Saucerman, J. J. (2010). Modeling cardiac B-adrenergic signaling with normalized-Hill differential equations: comparison with a biochemical model. BMC systems biology, 4(1), 1–12.

[11] Alon, U. (2006). An introduction to systems biology: design principles of biological circuits. Chapman and Hall/CRC.

[12] Chen, Z., King, W. C., Hwang, A., Gerstein, M., & Zhang, J. (2022). Deep-Velo: Single-cell transcriptomic deep velocity field learning with neural ordinary differential equations. Science Advances, 8(48), eabq3745.

[13] Farrell, S., Mani, M., & Goyal, S. (2022). Inferring single-cell dynamics with structured dynamical representations of RNA velocity. bioRxiv, 2022–08

[14] Aliee, H., Richter, T., Solonin, M., Ibarra, I., Theis, F., & Kilbertus, N. (2022). Sparsity in Continuous-Depth Neural Networks. arXiv preprint arXiv:2210.14672.

[15] Monti, M., Fiorentino, J., Milanetti, E., Gosti, G., & Tartaglia, G. G. (2022). Prediction of Time Series Gene Expression and Structural Analysis of Gene Regulatory Networks Using Recurrent Neural Networks. Entropy, 24(2), 141.

[16] La Manno, G., Soldatov, R., Zeisel, A., Braun, E., Hochgerner, H., Petukhov, V., … & Kharchenko, P. V. (2018). RNA velocity of single cells. Nature, 560(7719), 494–498.

[17] Hu, Y. (2022). Modeling the gene regulatory dynamics in neural differentiation with single cell data using a machine learning approach.

[18] Mao, G., Zeng, R., Peng, J., Zuo, K., Pang, Z., & Liu, J. (2022). Reconstructing gene regulatory networks of biological function using differential equations of multilayer perceptrons. BMC Bioinformatics, 23(1), 1–17.

[19] Bergen, V., Soldatov, R. A., Kharchenko, P. V., & Theis, F. J. (2021). RNA velocity—current challenges and future perspectives. Molecular systems biology, 17(8), e10282.

[20] Bergen, V., Lange, M., Peidli, S., Wolf, F. A., & Theis, F. J. (2020). Generalizing RNA velocity to transient cell states through dynamical modeling. Nature biotechnology, 38(12), 1408–1414.

[21] Cui, H., Maan, H., & Wang, B. (2022). DeepVelo: Deep Learning extends RNA velocity to multi-lineage systems with cell-specific kinetics. bioRxiv, 2022–04.

[22] Gayoso, A., Weiler, P., Lotfollahi, M., Klein, D., Hong, J., Streets, A. M., … & Yosef, N. (2022). Deep generative modeling of transcriptional dynamics for RNA velocity analysis in single cells. bioRxiv, 2022–08

[23] Gu, Y., Blaauw, D., & Welch, J. D. (2022). Bayesian inference of rna velocity from multi-lineage single-cell data. bioRxiv, 2022 07

[24] Karniadakis, G. E., Kevrekidis, I. G., Lu, L., Perdikaris, P., Wang, S., & Yang, L. (2021). Physics-informed machine learning. Nature Reviews Physics, 3(6), 422–440.

[25] Wolpert, D. H., & Macready, W. G. (1997). No free lunch theorems for optimization. IEEE transactions on evolutionary computation, 1(1), 67–82.

[26] Glass, K., Huttenhower, C., Quackenbush, J., & Yuan, G. C. (2013). Passing messages between biological networks to refine predicted interactions. PloS one, 8(5), e64832.

[27] Chen, R. T., Rubanova, Y., Bettencourt, J., & Duvenaud, D. K. (2018). Neural ordinary differential equations. Advances in neural information processing systems, 31

[28] Chen, R. T. Q. (2021). torchdiffeq (Version 0.2.2) [Computer software]. https://github.com/rtqichen/torchdiffeq

[29] Bhuva, D. D., Cursons, J., Smyth, G. K., & Davis, M. J. (2019). Differential co-expression-based detection of conditional relationships in transcriptional data: comparative analysis and application to breast cancer. Genome biology, 20(1), 1–21

[30] Gesztelyi, R., Zsuga, J., Kemeny-Beke, A., Varga, B., Juhasz, B., & Tosaki, A. (2012). The Hill equation and the origin of quantitative pharmacology. Archive for history of exact sciences, 66(4), 427–438.

[31] Grant, C. E., Bailey, T. L., & Noble, W. S. (2011). FIMO: scanning for occurrences of a given motif. Bioinformatics, 27(7), 1017–1018.

[32] Lawrence, M., Huber, W., Pag`es, H., Aboyoun, P., Carlson, M., Gentleman, R., … & Carey, V. J. (2013). Software for computing and annotating genomic ranges. PLoS computational biology, 9(8), e1003118.

[33] Aliee, H., Theis, F. J., & Kilbertus, N. (2021). Beyond Predictions in Neural ODEs: Identification and Interventions. arXiv preprint arXiv:2106.12430.

[34] Cheng, S., & Sabes, P. N. (2006). Modeling sensorimotor learning with linear dynamical systems. Neural computation, 18(4), 760–793.

[35] Bhuva, D. D. (2017). SimulatorGRN [Computer software]. https://github.com/DavisLaboratory/SimulatorGRN

[36] Hossain, I. (2022). PHOENIX package [Computer software]. https://github.com/QuackenbushLab/phoenix

[37] Pramila, T., Wu, W., Miles, S., Noble, W. S., & Breeden, L. L. (2006). The Forkhead transcription factor Hcm1 regulates chromosome segregation genes and fills the S-phase gap in the transcriptional circuitry of the cell cycle. Genes & development, 20(16), 2266–2278.

[38] Sirovich, L. (2020). A novel analysis of gene array data: yeast cell cycle. Biology Methods and Protocols, 5(1), bpaa018.

[39] Harbison, C. T., Gordon, D. B., Lee, T. I., Rinaldi, N. J., Macisaac, K. D., Danford, T. W., … & Young, R. A. (2004). Transcriptional regulatory code of a eukaryotic genome. Nature, 431(7004), 99–104.

[40] Ahnert, K., & Abel, M. (2007). Numerical differentiation of experimental data: local versus global methods. Computer Physics Communications, 177(10), 764–774.

[41] Desmedt, C., Piette, F., Loi, S., Wang, Y., Lallemand, F., Haibe-Kains, B., … & TRANSBIG Consortium. (2007). Strong time dependence of the 76-gene prognostic signature for node-negative breast cancer patients in the TRANSBIG multicenter independent validation series. Clinical cancer research, 13(11), 3207–3214.

[42] Chèneby, J., Gheorghe, M., Artufel, M., Mathelier, A., & Ballester, B. (2018). ReMap 2018: an updated atlas of regulatory regions from an integrative analysis of DNA-binding ChIP-seq experiments. Nucleic acids research, 46(D1), D267–D275.

[43] Artibani, M., Sims, A. H., Slight, J., Aitken, S., Thornburn, A., Muir, M., … & Hohenstein, P. (2017). WT1 expression in breast cancer disrupts the epithelial/mesenchymal balance of tumour cells and correlates with the metabolic response to docetaxel. Scientific reports, 7(1), 1–15.

[44] Brett, J. O., Spring, L. M., Bardia, A., & Wander, S. A. (2021). ESR1 mutation as an emerging clinical biomarker in metastatic hormone receptor-positive breast cancer. Breast Cancer Research, 23(1), 1–15.

[45] Kensler, K. H., Regan, M. M., Heng, Y. J., Baker, G. M., Pyle, M. E., Schnitt, S. J., … & Tamimi, R. M. (2019). Prognostic and predictive value of androgen receptor expression in postmenopausal women with estrogen receptor-positive breast cancer: results from the Breast International Group Trial 1–98. Breast Cancer Research, 21(1), 1–11.

[46] Lu, X. F., Zeng, D., Liang, W. Q., Chen, C. F., Sun, S. M., & Lin, H. Y. (2018). FoxM1 is a promising candidate target in the treatment of breast cancer. Oncotarget, 9(1), 842.

[47] Mandigo, A. C., Yuan, W., Xu, K., Gallagher, P., Pang, A., Guan, Y. F., … & Knudsen, K. E. (2021). RB/E2F1 as a Master Regulator of Cancer Cell Metabolism in Advanced DiseaseRB/E2F1 Regulates Cell Metabolism in Advanced Disease. Cancer discovery, 11(9), 2334–2353.

[48] Chen, H. Z., Tsai, S. Y., & Leone, G. (2009). Emerging roles of E2Fs in cancer: an exit from cell cycle control. Nature reviews cancer, 9(11), 785–797.

[49] Fang, C., Wang, Z., Han, C., Safgren, S. L., Helmin, K. A., Adelman, E. R., … & Zang, C. (2020). Cancer-specific CTCF binding facilitates oncogenic transcriptional dysregulation. Genome biology, 21, 1–30.

[50] Adamo, P., Ladomery, M. (2016). The oncogene ERG: a key factor in prostate cancer. Oncogene 35, 403–414.

[51] Pathania, R., Ramachandran, S., Elangovan, S., Padia, R., Yang, P., Cinghu, S., … & Thangaraju, M. (2015). DNMT1 is essential for mammary and cancer stem cell maintenance and tumorigenesis. Nature communications, 6(1), 6910.

[52] Gillespie, M., Jassal, B., Stephan, R., Milacic, M., Rothfels, K., Senff-Ribeiro, A., … & D’Eustachio, P. (2022). The reactome pathway knowledgebase 2022. Nucleic acids research, 50(D1), D687–D692.

[53] Hanahan, D., & Weinberg, R. A. (2011). Hallmarks of cancer: the next generation. cell, 144(5), 646–674.

[54] Okal, A., Matissek, K. J., Matissek, S. J., Price, R., Salama, M. E., Janát-Amsbury, M. M., & Lim, C. S. (2014). Re-engineered p53 activates apoptosis in vivo and causes primary tumor regression in a dominant negative breast cancer xenograft model. Gene therapy, 21(10), 903–912.

[55] Le Romancer, M., Poulard, C., Cohen, P., Sentis, S., Renoir, J. M., & Corbo, L. (2011). Cracking the estrogen receptor’s posttranslational code in breast tumors. Endocrine reviews, 32(5), 597–622.

[56] Grund-Gröschke, S., Stockmaier, G. & Aberger, F. (2019). Hedgehog/GLI signaling in tumor immunity - new therapeutic opportunities and clinical implications. Cell Commun Signal 17, 172.

[57] Wang, X., & Yang, D. (2021). The regulation of RNA metabolism in hormone signaling and breast cancer. Molecular and cellular endocrinology, 529, 111221.

[58] Gallo, C., Fragliasso, V., Donati, B. et al. (2018). The bHLH transcription factor DEC1 promotes thyroid cancer aggressiveness by the interplay with NOTCH1. Cell Death Dis 9, 871.

[59] Madden, S.K., de Araujo, A.D., Gerhardt, M. et al. (2021). Taking the Myc out of cancer: toward therapeutic strategies to directly inhibit c-Myc. Mol Cancer 20, 3.

[60] Mias G.I., Brooks, L.R. (2018). Integrated Transcriptomic and Proteomic Dynamics of Rituximab Treatment in Primary B Cells. GEO data deposit, GSE100441.

[61] Xing, Y., Igarashi, H., Wang, X., Sakaguchi, N. (2005). Protein phosphatase subunit G5PR is needed for inhibition of B cell receptor-induced apoptosis. Journal of Experimental Medicine, 202(5), 707–719.

[62] Gao, T., Furnari, F., Newton, A. C. (2005). PHLPP: a phosphatase that directly dephosphorylates Akt, promotes apoptosis, and suppresses tumor growth. Molecular Cell, 18(1), 13–24.

[63] Downward, J. (1998) Ras signalling and apoptosis. Current Opinion in Genetics & Development, 8(1), 49–54.

[64] Pla-Martín, D., Schatton, D., Wiederstein, J.L., Marx, MC, Khiati, S., Krüger, M., Rugarli, E. (2020). CLUH granules coordinate translation of mitochondrial proteins with mTORC1 signaling and mitophagy. The EMBO Journal, 39(9), e102731.

[65] Campbell, K. R., & Yau, C. (2018). Uncovering pseudotemporal trajectories with covariates from single cell and bulk expression data. Nature communications, 9(1), 2442.

[66] Van den Bulcke, T., Van Leemput, K., Naudts, B., van Remortel, P., Ma, H., Verschoren, A., … & Marchal, K. (2006). SynTReN: a generator of synthetic gene expression data for design and analysis of structure learning algorithms. BMC bioinformatics, 7(1), 1–12.

[67] Yang, Y. H., & Paquet, A. C. (2005). Preprocessing two-color spotted arrays. In Bioinformatics and Computational Biology Solutions Using R and Bioconductor (pp. 49–69). Springer, New York, NY.

[68] Johnson, W. E., Li, C., & Rabinovic, A. (2007). Adjusting batch effects in microarray expression data using empirical Bayes methods. Biostatistics, 8(1), 118–127.

[69] Leek, J. T., Johnson, W. E., Parker, H. S., Jaffe, A. E., & Storey, J. D. (2012). The sva package for removing batch effects and other unwanted variation in high-throughput experiments. Bioinformatics, 28(6), 882–883.

[70] Weighill, D., Guebila, M. B., Lopes-Ramos, C., Glass, K., Quackenbush, J., Platig, J., & Burkholz, R. (2021, May). Gene regulatory network inference as relaxed graph matching. In Proceedings of the AAAI Conference on Artificial Intelligence (Vol. 35, No. 11, pp. 10263–10272).

[71] Bailey, T. L., Boden, M., Buske, F. A., Frith, M., Grant, C. E., Clementi, L., … & Noble, W. S. (2009). MEME SUITE: tools for motif discovery and searching. Nucleic acids research, 37(suppl 2), W202–W208.

[72] Liberzon, A., Birger, C., Thorvaldsdóttir, H., Ghandi, M., Mesirov, J. P., & Tamayo, P. (2015). The molecular signatures database hallmark gene set collection. Cell systems, 1(6), 417–425.

[73] Luo, W., Brouwer, C. (2013). Pathview: an R/Bioconductor package for pathway-based data integration and visualization. Bioinformatics 29 (14), 1830–1831.

[74] Karlsson, D., & Svanström, O. (2019). Modelling Dynamical Systems Using Neural Ordinary Differential Equations. [master’s thesis], Chalmers University of Technology

