## Supplemental Results & Methods for "Biologically informed NeuralODEs for genome-wide regulatory dynamics"

### Online supplement accompanying: "Biologically informed neural ordinary differential equations for genome-wide regulatory dynamics"

Intekhab Hossain[1*], Viola Fanfani[1], Jonas Fischer[1], John Quackenbush[1] and Rebekka Burkholz[2]

[1*]Department of Biostatistics, Harvard T.H. Chan School of Public Health, Boston, MA, USA.
[2]Helmholtz Center for Information Security (CISPA), Saarbrücken, Germany.

Contributing authors:;
;
;

**This is the supplementary information file accompanying the paper.** This document is divided into a **Supplemental Methods** and a **Supplemental Results** section. For further queries, please feel free to reach out to ``

- Availability of data and materials: All open-source datasets used in the paper have been referenced with their GEO accession ID. Furthermore, relevant data files have also been deposited on Zenodo (`10.5281/zenodo.10412968`). The source code is available as an open-source implementation (distributed under license **CC BY-NC 4.0**) via Github: https://github.com/QuackenbushLab/phoenix [1], and has also been deposited on Zenodo (`10.5281/zenodo.10412968`).

### Supplemental results

#### Supp. Results 1 PHOENIX *in silico* predictive accuracy

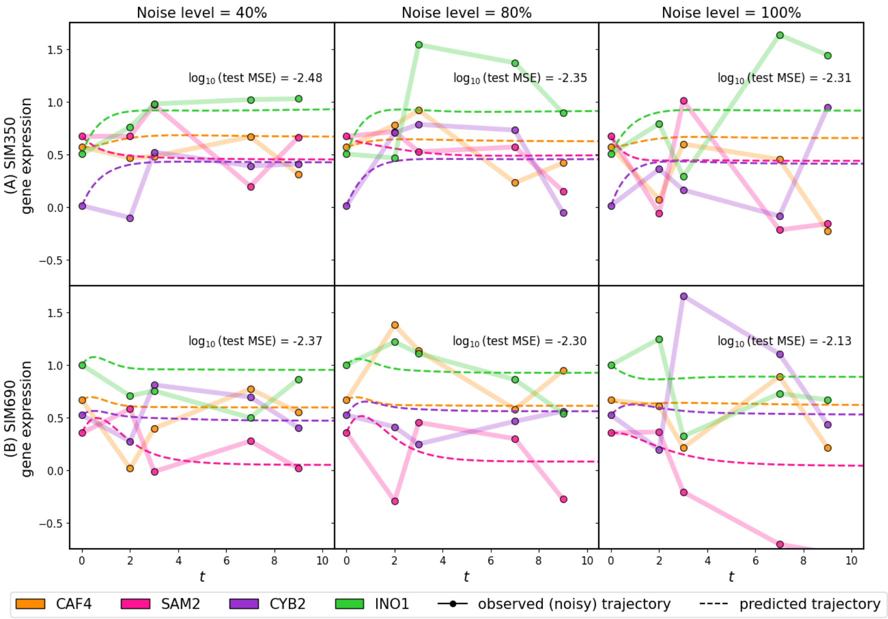

**Supp. Fig. 1** We applied PHOENIX to simulated gene expression data originating from two different *in silico* dynamical systems SIM350 (A) and SIM690 (B) that simulate the temporal expression of 350 and 690 genes respectively. Each simulated trajectory consisted of five time points ($t = 0, 2, 3, 7, 9$) and was subjected to varying levels of Gaussian noise ($\frac{\text{noise } \sigma}{\text{mean}}$ = 40%, 80%, 100%). **These results are provided in addition to the noise settings of 0%, 5%, 10%, and 20% that have been shown in Figure 2 of the main paper**. Since PHOENIX uses a user-defined prior network model as a regularizer, we also corrupted the prior models up to an amount commensurate with the noise level. For each noise setting we trained PHOENIX on 140 of these "observed" trajectories and validated on 10. The performance on the validation trajectories was used to determine the optimal value of $\lambda_{\text{prior}}$. We then tested the trained model (with the optimal choice of $\lambda_{\text{prior}}$) on 10 new test set trajectories. We display both observed and predicted test set trajectories for four arbitrary genes in both SIM350 and SIM690, across all noise settings. We display the mean squared error (MSE) between the predictions and the 10 pre-noise test set trajectories.

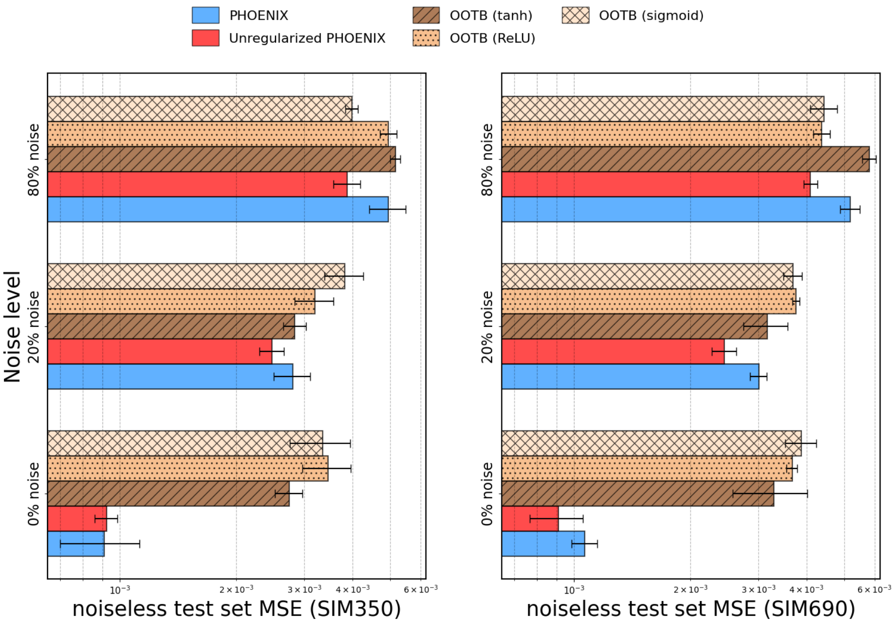

**Supp. Fig. 2** We measured the marginal contribution of PHOENIX's architecture and incorporation of prior information by comparing against the baseline contributions of out-of-the-box NeuralODE models with three different activation functions, across both *in silico* dynamical systems SIM350 and SIM690, after subjecting the data to different amounts of noise. We display the performance of the out-of-the-box models, as well as PHOENIX ($\lambda_{\text{prior}}$ tuned using the validation set) and its unregularized version ($\lambda_{\text{prior}} = 0$), in terms of how well held out time points from noiseless test set trajectories could be predicted after training on trajectories from different noise settings. The entire procedure was repeated five times to generate average mean-squared error (MSE) values and error bars.

### Supp. Results 2 PHOENIX *in silico* explainability: unabridged version

**Supp. Table 1** Assessing the contribution of priors to PHOENIX explainability, as well as the effect of prior misspecification, in both *in silico* experiments and real experimental data

| | Genes | Noise | PHOENIX | Prior constraints |
|---|---|---|---|---|
| SIM350[1] | 350 | 0% | 0.987 | 1.000 |
| | 350 | 5% | 0.980 | 0.970 |
| | 350 | 10% | 0.979 | 0.950 |
| | 350 | 20% | 0.943 | 0.900 |
| | 350 | 40% | 0.761 | 0.800 |
| | 350 | 80% | 0.538 | 0.600 |
| | 350 | 100% | 0.520 | 0.500 |
| SIM690[1] | 690 | 0% | 0.986 | 1.000 |
| | 690 | 5% | 0.984 | 0.975 |
| | 690 | 10% | 0.983 | 0.950 |
| | 690 | 20% | 0.958 | 0.900 |
| | 690 | 40% | 0.737 | 0.800 |
| | 690 | 80% | 0.531 | 0.600 |
| | 690 | 100% | 0.529 | 0.500 |
| Yeast[2] | 3551 | - | 0.934 | 0.790 |
| Breast[2] | 500 | - | 0.958 | 0.820 |
| | 2000 | - | 0.906 | 0.840 |
| | 4000 | - | 0.909 | 0.810 |
| | 11165 | - | 0.904 | 0.810 |

[1]For *in silico* experiments (SIM350, SIM690) the prior knowledge model was corrupted by an amount commensurate with the noise level (see Supp. Methods 4.2). Then for each scenario, a network representation of the misspecified prior model was checked for how well it aligned with the ground truth GRN in terms of AUC. This AUC was compared to that obtained by aligning the ground truth GRN against a network extracted from a PHOENIX model trained using the misspecified prior in question.

[2]This same approach of aligning a "ground truth" GRN against extracted and prior networks was repeated for the real data sets (yeast cell cycle and breast cancer). But this time the prior networks came from the motiff-prior models used in model-fitting (see Methods 2.2 and 3.2), and the "ground-truth" GRNs were experimentally verified ChIP-Seq networks describing transcription factor binding [2, 3].

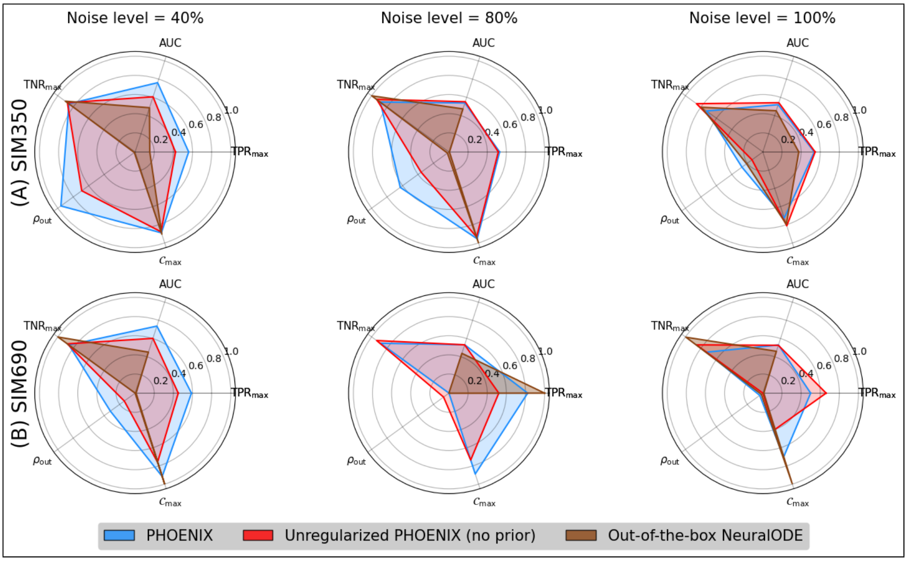

**Supp. Fig. 3** We extracted encoded GRNs from the trained PHOENIX models and the best-performing out-of-the-box NeuralODE models, for both *in silico* dynamical systems SIM350 (A) and SIM690 (B) across all noise settings. Here we display noise settings 40%, 80% and 100%. **These results are provided in addition to the noise settings of 0%, 5%, 10%, and 20% that have been shown in Figure 3 of the main paper**.We compared these GRN estimates to the corresponding ground truth GRNs used to formulate SIM350 and SIM690, and obtained AUC values as well as out-degree correlations ($\rho_{\text{out}}$). We also reverse-engineered a metric ($\mathcal{C}_{\max}$) to inform how sparsely PHOENIX had inferred the dynamics (see Supp. Methods 2). Furthermore, we used these $\mathcal{C}_{\max}$ values to obtain optimal true positive and true negative rates ($\text{TPR}_{\max}$ and $\text{TNR}_{\max}$) that were independent of any cutoff value, allowing us to compare between "best possible" networks across all settings.

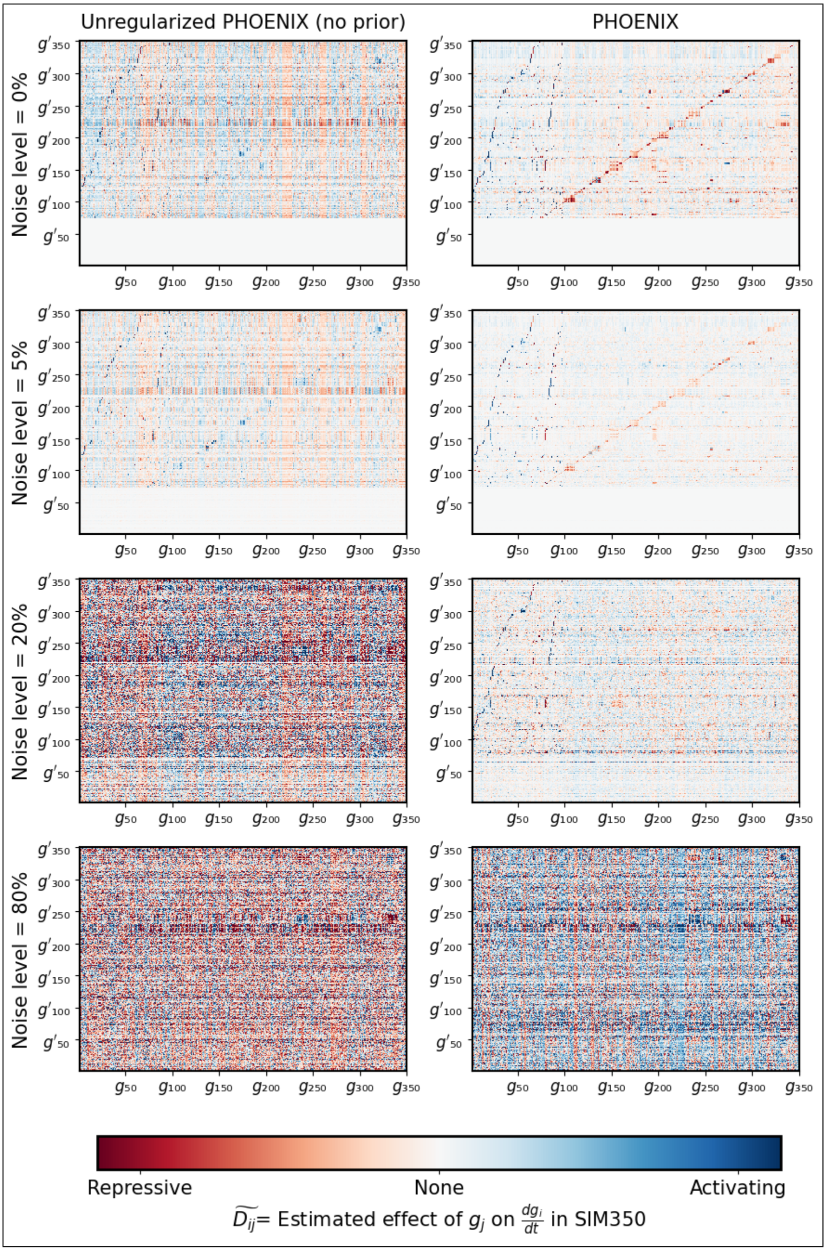

**Supp. Fig. 4** We investigated *how* PHOENIX used prior information to sparsify the learned dynamics by computing the encoded 350 × 350 dynamics matrices $\widetilde{\boldsymbol{D}}$ (see Supp. Methods 2), across multiple noise settings in SIM350. We display $\widetilde{\boldsymbol{D}}$ for both PHOENIX as well as its unregularized ($\lambda_{\text{prior}} = 0$) version as heatmaps of effect size. Unlike its unregularized counterpart, PHOENIX identifies core elements of the dynamics even at relatively high noise levels.

### Supp. Results 3 Exploring aspects of the PHOENIX architecture

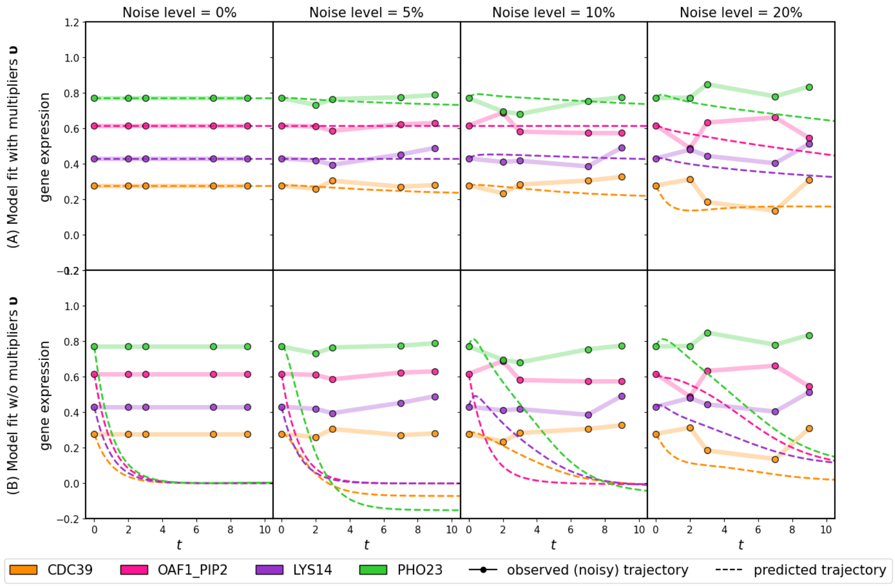

**Supp. Fig. 5** PHOENIX uses gene-specific multipliers $\boldsymbol{\upsilon} \in \mathbb{R}^n$ (see Model Formulation in main paper) to simplify the representation of steady state for genes without upstream transcription factors $\left(\frac{dg_i(t)}{dt} = 0, \forall t\right)$. In order to explore the contribution of these gene-specific multipliers towards the PHOENIX model, we tested PHOENIX with (A) and without (B) the multipliers using gene expression data originating from the *in silico* dynamical system SIM690. We used SIM690 to simulate the temporal gene expression patterns for its 690 genes. Each simulated trajectory consisted of five time points ($t = 0, 2, 3, 7, 9$) and was subjected to varying levels of Gaussian noise ($\frac{\text{noise } \sigma}{\text{mean}}$ = 0%, 5%, 10%, and 20% ). We also corrupted the user-defined prior models up to an amount commensurate with the noise level. We display across all noise settings both observed and predicted trajectories for four arbitrary genes that had true flat trajectories i.e $\left(\frac{dg_i(t)}{dt} = 0, \forall t\right)$ in SIM690.

**Supp. Table 2** In order to incorporate prior domain knowledge, PHOENIX uses an adjacency matrix $\boldsymbol{A}$ of likely network structure based on user-defined biological insights (including experimentally validated interactions, motif map of promoter targets, etc.). Here, $\boldsymbol{A}_{ij} \in \{+1, -1, 0\}$ representing an activating, repressive, or no prior interaction, respectively. But for real world scenario, the signs (activating/repressive) of prior interactions are often unknown. So we wanted to check whether formulating $\boldsymbol{A}$ simply based on *prior interaction existence*, $\boldsymbol{A}_{ij} \in \{+1, 0\}$, would suffice. Below we show the results of testing the importance of providing the correct prior signs to PHOENIX using our *in silico* setup where the ground truth signs were always known. We tested multiple forms of regularized PHOENIX models and report results both in terms predictive accuracy (MSE on test set) and explainability (AUC from comparing inferred dynamics to ground truth GRNs).

| | | Predictive performance (MSE) | | | Explainability (AUC) | | |
|---|---|---|---|---|---|---|---|
| | Noise | $PHX_0$[1] | $PHX_+$[2] | PHX[3] | $PHX_0$[1] | $PHX_+$[2] | PHX[3] |
| SIM350 | 0% | 0.0009 | 0.0015 | 0.0009 | 0.94 | 0.97 | 0.99 |
| | 5% | 0.0010 | 0.0012 | 0.0011 | 0.90 | 0.97 | 0.98 |
| | 10% | 0.0019 | 0.0019 | 0.0014 | 0.86 | 0.94 | 0.98 |
| | 20% | 0.0025 | 0.0029 | 0.0028 | 0.77 | 0.92 | 0.94 |
| | 40% | 0.0031 | 0.0039 | 0.0037 | 0.61 | 0.75 | 0.76 |
| | 80% | 0.0039 | 0.0049 | 0.0050 | 0.55 | 0.54 | 0.54 |
| | 100% | 0.0045 | 0.0055 | 0.0059 | 0.54 | 0.53 | 0.52 |
| SIM690 | 0% | 0.0009 | 0.0012 | 0.0011 | 0.85 | 0.98 | 0.99 |
| | 5% | 0.0018 | 0.0016 | 0.0012 | 0.80 | 0.97 | 0.98 |
| | 10% | 0.0024 | 0.0021 | 0.0019 | 0.74 | 0.96 | 0.98 |
| | 20% | 0.0025 | 0.0033 | 0.0030 | 0.69 | 0.91 | 0.95 |
| | 40% | 0.0032 | 0.0044 | 0.0042 | 0.60 | 0.73 | 0.74 |
| | 80% | 0.0041 | 0.0053 | 0.0052 | 0.53 | 0.52 | 0.53 |
| | 100% | 0.0042 | 0.0067 | 0.0070 | 0.53 | 0.52 | 0.53 |

[1] $PHX_0$ = Unregularized PHOENIX where $\lambda_{\text{prior}} = 0$ (ignoring the network prior altogether).

[2] $PHX_+$ = Regularized PHOENIX where $\lambda_{\text{prior}}$ was chosen based on the validation set, but both activating and repressive edges in the prior were set to +1. "No prior interaction" was kept to 0. This emulated a setting where we didn't have prior knowledge of these signs (either activating vs repressive prior edge).

[3] PHX = Regularized PHOENIX where $\lambda_{\text{prior}}$ was chosen based on the validation set. Each entry in the prior was correctly set to +1 or -1, depending on whether the prior edge was activating or repressive. "No prior interaction" was set to 0.

### Supp. Results 4 Comparing PHOENIX to black-box models

**Supp. Table 3** Benchmarking PHOENIX against other methods *in silico*, in terms of trajectory-recovery performance (performance metric is MSE on test set)

| | | Trajectory based | | | | Velocity based | | |
|---|---|---|---|---|---|---|---|---|
| | Noise | PHX[1] | $PHX_0$[2] | OOTB[3] | PRESC[4] | Dynamo[5] | RNODE[6] | DeepVelo[7] |
| SIM350 | 0% | 0.0009 | 0.0009 | 0.0027 | 0.0046 | 0.0032 | 0.0052 | 0.0062 |
| | 5% | 0.0011 | 0.0010 | 0.0029 | 0.0043 | 0.0034 | 0.0074 | 0.0083 |
| | 10% | 0.0014 | 0.0019 | 0.0029 | 0.0045 | 0.0040 | 0.0084 | 0.0086 |
| | 20% | 0.0028 | 0.0025 | 0.0028 | 0.0048 | 0.0052 | 0.0104 | 0.0122 |
| | 40% | 0.0037 | 0.0031 | 0.0042 | 0.0061 | 0.0066 | 0.0098 | 0.0093 |
| | 80% | 0.0050 | 0.0039 | 0.0052 | 0.0066 | 0.0074 | 0.0090 | 0.0108 |
| | 100% | 0.0059 | 0.0045 | 0.0061 | 0.0065 | 0.0080 | 0.0095 | 0.0151 |
| SIM690 | 0% | 0.0011 | 0.0009 | 0.0033 | 0.0046 | 0.0042 | 0.0076 | 0.0075 |
| | 5% | 0.0012 | 0.0018 | 0.0029 | 0.0051 | 0.0057 | 0.0084 | 0.0074 |
| | 10% | 0.0019 | 0.0024 | 0.0035 | 0.0057 | 0.0063 | 0.0083 | 0.0086 |
| | 20% | 0.0030 | 0.0025 | 0.0032 | 0.0056 | 0.0061 | 0.0109 | 0.0099 |
| | 40% | 0.0042 | 0.0032 | 0.0046 | 0.0055 | 0.0077 | 0.0100 | 0.0100 |
| | 80% | 0.0052 | 0.0041 | 0.0058 | 0.0126 | 0.0107 | 0.0146 | 0.0103 |
| | 100% | 0.0070 | 0.0042 | 0.0071 | 0.0133 | 0.0099 | 0.0183 | 0.0108 |

[1]PHX = Regularized PHOENIX where $\lambda_{\text{prior}}$ was chosen based on the validation set.

[2]$PHX_0$ = Unregularized PHOENIX where $\lambda_{\text{prior}} = 0$ (ignoring the network prior).

[3]OOTB = Out-of-the-box NeuralODE, resembling how plain NeuralODEs are typically used for this problem [4, 5]. We note that the method RNAForecaster [4] uses an OOTB approach with ReLU activation, but the results here are for the *tanh* activation function that had better performance than both sigmoid and ReLU on the validation set.

[4]PRESC = PRESCIENT [6], where we used the validation set to optimize the value of $k_{dim}$, which is the number of neurons use in PRESCIENT's hidden layer.

[5]For Dynamo [7] we used the validation set to optimize the sparsity regularization penalty ($\lambda$), as well as the number of kernel basis functions use to approximate the vector field ($M$)

[6]RNODE = RNA-ODE [8], where we used the validation set to optimize the number of trees used in the random forest function

[7]In DeepVelo [9] the encoder and the decoder consisted of four dense layers (size $4p$ for the intermediate layers and size $p$ for the latent layer) with ReLU activation. We used the validation set to optimize $p$.

**Supp. Table 4** Benchmarking PHOENIX against other methods *in silico*, in terms of explainability. A GRN describing the inferred dynamics was extracted for each fitted model, and was compared to the ground truth GRN to calculate an AUC

| | | Trajectory based | | | | Velocity based | | |
|---|---|---|---|---|---|---|---|---|
| | Noise | PHX[1] | $PHX_0$[1] | OOTB[2] | PRESC[3] | Dynamo[4] | RNODE[5] | DeepVelo[6] |
| SIM350 | 0% | 0.987 | 0.942 | 0.580 | N/A | 0.784 | 0.683 | 0.768 |
| | 5% | 0.980 | 0.899 | 0.625 | N/A | 0.776 | 0.659 | 0.744 |
| | 10% | 0.979 | 0.858 | 0.606 | N/A | 0.762 | 0.593 | 0.704 |
| | 20% | 0.943 | 0.766 | 0.543 | N/A | 0.741 | 0.599 | 0.732 |
| | 40% | 0.761 | 0.606 | 0.486 | N/A | 0.717 | 0.596 | 0.751 |
| | 80% | 0.538 | 0.552 | 0.473 | N/A | 0.667 | 0.607 | 0.724 |
| | 100% | 0.520 | 0.541 | 0.452 | N/A | 0.646 | 0.606 | 0.701 |
| SIM690 | 0% | 0.986 | 0.847 | 0.694 | N/A | 0.733 | 0.730 | 0.822 |
| | 5% | 0.984 | 0.800 | 0.599 | N/A | 0.710 | 0.634 | 0.768 |
| | 10% | 0.983 | 0.744 | 0.580 | N/A | 0.707 | 0.614 | 0.768 |
| | 20% | 0.953 | 0.686 | 0.557 | N/A | 0.698 | 0.523 | 0.784 |
| | 40% | 0.737 | 0.602 | 0.450 | N/A | 0.652 | 0.455 | 0.729 |
| | 80% | 0.531 | 0.531 | 0.437 | N/A | 0.644 | 0.476 | 0.732 |
| | 100% | 0.529 | 0.525 | 0.460 | N/A | 0.625 | 0.395 | 0.729 |

[1]The architecture of PHOENIX allowed GRNs to be extracted from both regularized and unregularized versions using a very simple and efficient algorithm (see Supp. Methods 2)

[2]GRNs were extracted from out-of-the-box NeuralODEs, using sensitivity analyses (see Supp. Methods 3.2)

[3]PRESCIENT does not provide a straightforward means for extracting a GRN describing the dynamics [6]

[4]Dynamo provides a function that calculates the Jacobian matrix of the fitted model at any data point [7]. We used this function to calculate an average Jacobian matrix across several randomly generated data points and estimate a GRN (see Supp. Methods 3.3)

[5]RNA-ODE provides a function for GRN inference by ranking regulatory links using estimated effect sizes [8] (see Supp. Methods 3.3)

[6]DeepVelo provides a function for estimating gene correlation networks from simulated retrograde trajectories [9]. We generated $n = 200$ retrograde trajectories using 200 randomly generated initial conditions (see Supp. Methods 3.3)

**Supp. Table 5** Benchmarking PHOENIX against other methods *in silico*, in terms of sparsity of inferred dynamics. Sparsity was calculated here as the average out-degree of the extracted GRN from each of the inferred dynamical systems

| | | Trajectory based | | | | Velocity based | | |
|---|---|---|---|---|---|---|---|---|
| | Noise | PHX | $PHX_0$ | OOTB | PRESC[1] | Dynamo | RNA-ODE | DeepVelo |
| SIM350 | 0% | 3 | 10 | 81 | N/A | 94 | 97 | 37 |
| | 5% | 4 | 15 | 50 | N/A | 65 | 48 | 38 |
| | 10% | 8 | 23 | 41 | N/A | 113 | 23 | 40 |
| | 20% | 11 | 45 | 80 | N/A | 109 | 75 | 82 |
| | 40% | 39 | 43 | 34 | N/A | 178 | 76 | 64 |
| | 80% | 16 | 23 | 1 | N/A | 265 | 73 | 35 |
| | 100% | 96 | 66 | 72 | N/A | 272 | 78 | 66 |
| SIM690 | 0% | 4 | 29 | 144 | N/A | 218 | 147 | 79 |
| | 5% | 5 | 44 | 109 | N/A | 205 | 54 | 85 |
| | 10% | 12 | 64 | 130 | N/A | 171 | 48 | 108 |
| | 20% | 13 | 86 | 138 | N/A | 254 | 127 | 149 |
| | 40% | 58 | 166 | 2 | N/A | 73 | 43 | 170 |
| | 80% | 77 | 183 | 689 | N/A | 402 | 35 | 131 |
| | 100% | 205 | 413 | 4 | N/A | 501 | 21 | 63 |

[1]PRESCIENT [6] does not provide a straightforward means for extracting a GRN that describes the dynamics

**Supp. Table 6** Details about additional black-box methods for estimating gene expression dynamics that we excluded from our *in silico* benchmarking experiments. We provide details about each method, and some reasoning behind its exclusion

| Method | Approach for estimating dynamics | Notes on exclusion |
|---|---|---|
| PROB[10] | Uses a local Euler approximation to calculate RNA velocity $d\boldsymbol{x}/dt$. Then describes this RNA velocity as a function of gene-expression through a linear model with quadratic interaction terms between all pairs of genes. Bayesian Lasso is used to induce sparsity. | The linear model is too simplistic, and may not accurately reflect complex regulatory patterns. Also, the method seeks to directly estimate pairwise gene effects ($\mathcal{O}(n^2)$ parameters), and hence has not been shown to scale beyond 100 genes. |
| LatentVelo[11] | Embeds spliced and unspliced RNA counts into a low dimensional latent space using a variational autoencoder (VAE), and then infers dynamics in this latent space with a NeuralODE, with soft constraints on the interplay between latent spliced and latent unspliced counts. | Dynamics can only be inferred in a low-dimensional latent space, making it difficult to derive interpretable insights and compare explainability. We already benchmarked against a VAE-based model (DeepVelo [9]) that - unlike LatentVelo - provides a GRN interpreting the inferred dynamics. |
| scTour[12] | Embeds spliced RNA counts into a low dimensional latent space using a variational autoencodeer, and then infers dynamics in this latent space with a NeuralODE. | Same reasoning as above. |
| PBA[13] | Uses spectral-graph theory to solve multi-dimensional Fokker-Plank equations on an empirical grid formed by expression values. It assumes that velocity fields are gradients of a potential landscape in gene expression space $\boldsymbol{J} = -\nabla F$. | PBA ostensibly ignores oscillatory gene expression dynamics (the cell cycle) [13], and hence may not be flexible enough. Also, we already benchmarked against another method (PRESCIENT [6]), that assumes dynamics to be the gradient of a scalar potential. |
| PathReg [14] | Uses an out-of-the-box NeuralODE with an Exponential Linear Unit (ELU) activation function and enforces both weight and feature sparsity at the path-level via a differentiable $L_0$-based regularizer in the loss function. A matrix of non-negative stochastic gates regularizes the probability of any path throughout the entire network contributing to a given output. This constrains the number of input-output paths. | PathReg was primarily developed to induce sparsity in NeuralODEs, and is similar in principle to the out-of-the-box models that we already benchmarked. Consequently, is has only been demonstrated to scale up to 529 highly variable dimensions/genes [14]. |

**Supp. Table 7** Qualitative comparison of black-box methods for estimating gene expression dynamics that we benchmarked against PHOENIX *in silico*. We provide details about inputs, learning algorithms, and key performance metrics

| | | | Scalability | Explainability | | Notes |
|---|---|---|---|---|---|---|
| | **Method** | **Description of approach for estimating dynamical system** | Demonstrated performance on $> 10^4$ genes without dimension reduction | Can extract GRN that describes dynamics | Flexibly incorporates prior/domain knowledge | ✳ Hyperparameters we optimized using val. set<br>✂ GRN extraction details |
| Velocity based (two step) | Dynamo [7] | Can use multiple modalities of seq data to estimate RNA velocity using both deterministic and stochastic approaches, then fits a sparse vector field mapping expression to velocity using Gaussian kernel regression | ✘ (Demonstrated on $< 10^4$ genes using dimension reduction via PCA/UMAP) | ✓ | ✘ | ✳ Control points ($M$) and sparsity parameter ($\lambda$)<br>✂ Monte Carlo approach to estimate GRN using Jacobians provided by tool (see Supp. Methods 3.3) |
| | RNA-ODE [8] | Uses scRNA transcriptome and RNA velocity to fit random forests mapping expression to velocity | ✘ (Demonstrated on 3001 genes) | ✓ | ✘ | ✳ Number of trees (nTrees)<br>✂ Tool provides GRN (see Supp. Methods 3.3) |
| | DeepVelo [9] | Uses mRNA counts and corresponding and RNA velocity (from scVelo [15]) to fit variational autoencoders mapping expression to velocity | ✘ (Demonstrated on 3000 highly variable genes) | ✓ | ✘ | ✳ Number of neurons<br>✂ Tool provides gene correlation matrix by simulating retrograde trajectories (see Supp. Methods 3.3) |
| Traj. based (one step) | PRESCIENT [6] | Uses time-series scRNA-seq and cell-growth rate data to learn a potential function $\Psi$ with a neural network. Final drift model is obtained using automatic differentiation $\mu = -\nabla\Psi$ | ✘ (Demonstrated on 2500 highly variable genes or on top$K$ PC) | ✘ | ✘ | ✳ Number of neurons inside each hidden layer ($k_{\text{dim}}$) (see Supp. Methods 3.3) |
| | Out-of-the-box NeuralODE [4, 5] | Uses time-series expression and NeuralODE out-of-the-box, with traditional activation functions | ✘ (Demo. on 2000 high expressed genes) | ✓ | ✘ | ✂ Monte Carlo approach using sensitivity analysis (see Supp. Methods 3.2) |
| | PHOENIX | Our method | ✓ | ✓ | ✓ | ✳ Prior weight ($\lambda_{\text{prior}}$)<br>✂ Algo. (Supp. Methods 2) |

### Supp. Results 5 Applying PHOENIX to yeast cell-cycle data

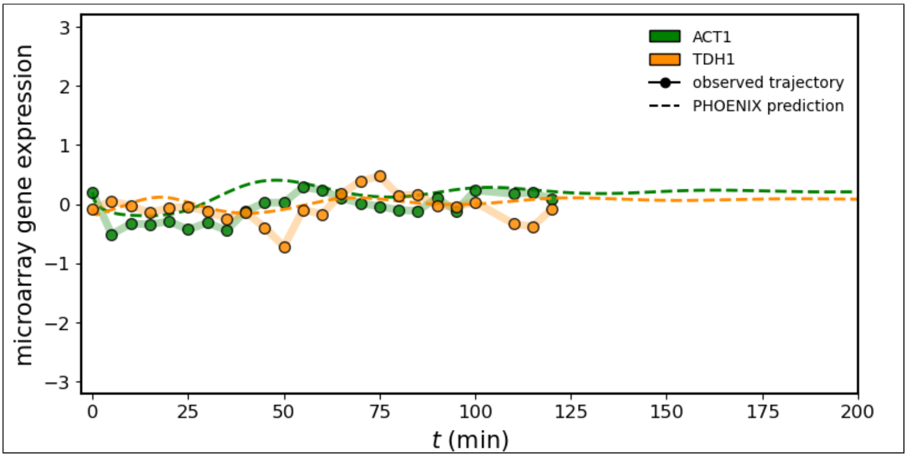

**Supp. Fig. 6** We applied PHOENIX ($\lambda_{prior} = 0.05$) to 2 technical replicates of gene expression of 3551 genes each, collected across 24 time points in a yeast cell-cycle time course [16]. We trained on 40 transition pairs, used 3 for validation, and tested predictive accuracy on the remaining 3. We display both observed and predicted trajectories for ACT1 and TDH1, where the predicted trajectories are extrapolations into future time points based on just initial values (gene expression at $t = 0$).

### Supp. Results 6 Applying PHOENIX to breast cancer data

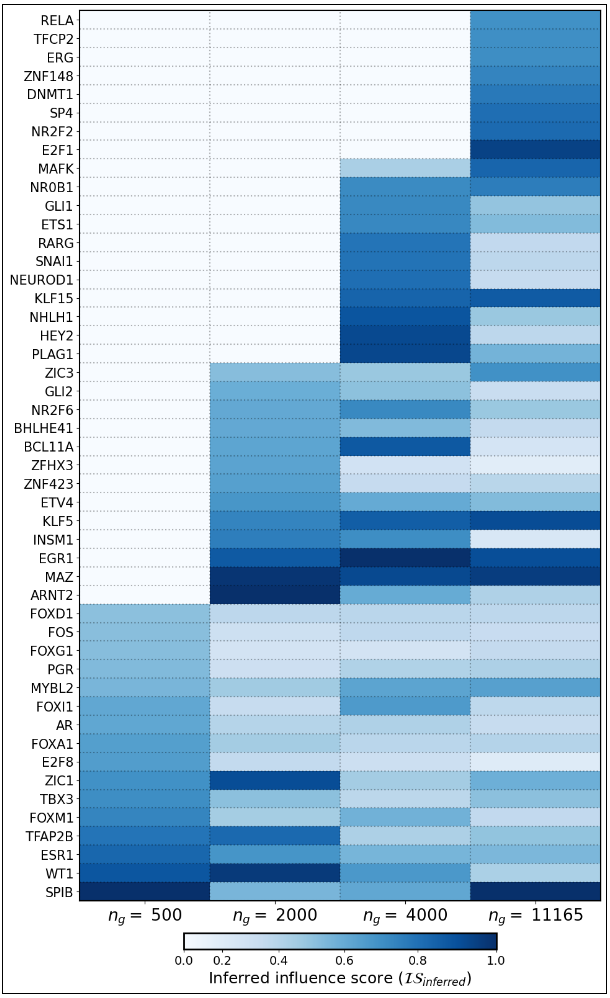

**Supp. Fig. 7** We applied PHOENIX to a pseudotrajectory of 186 breast cancer samples (ordered along subsequent "pseudotimepoints") consisting of $n_g = 11165$ genes [17]. We also repeated the analysis on smaller subsets of genes $n_g = 500, 2000, 4000$, where we subsetted the full trajectory to only the $n_g$ most variable genes in the pseudotrajectory. We used the trained PHOENIX models to extract influence scores for individual genes in the estimated system (see Methods 3.3), and visualized influence scores for the most central genes across different values of $n_g$. For visualization purposes, the influence scores are normalized within each column (each value of $n_g$) to be between 0 and 1. Genes that are excluded from the subset of $n_g$ most variable genes were assigned $\mathcal{IS}_{inferred} = 0$.

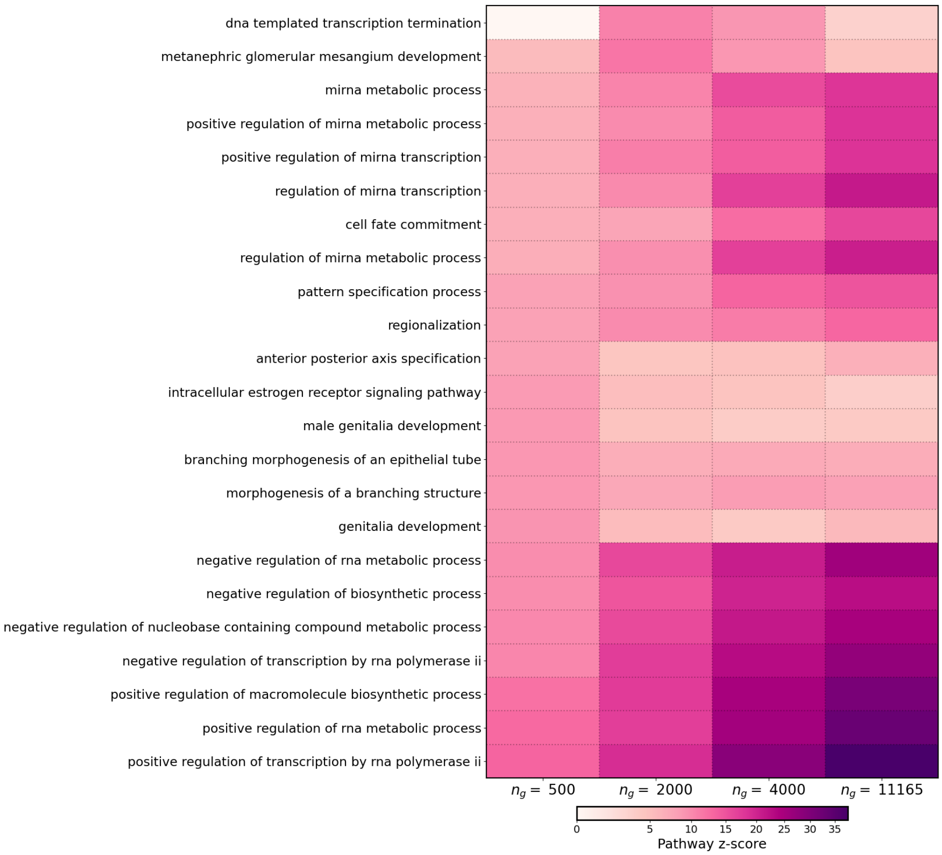

**Supp. Fig. 8** We applied PHOENIX to a pseudotrajectory of 186 breast cancer samples (ordered along subsequent "pseudotimepoints") consisting of $n_g = 11165$ genes [17]. We also repeated the analysis on smaller subsets of genes $n_g = 500, 2000, 4000$, where we subsetted the full trajectory to only the $n_g$ most variable genes in the pseudotrajectory. We used the trained PHOENIX models to extract influence scores for pathways in the **Gene Ontology (biological process)** database (see Methods 3.4), and visualized influence scores for the most central pathways across different values of $n_g$.

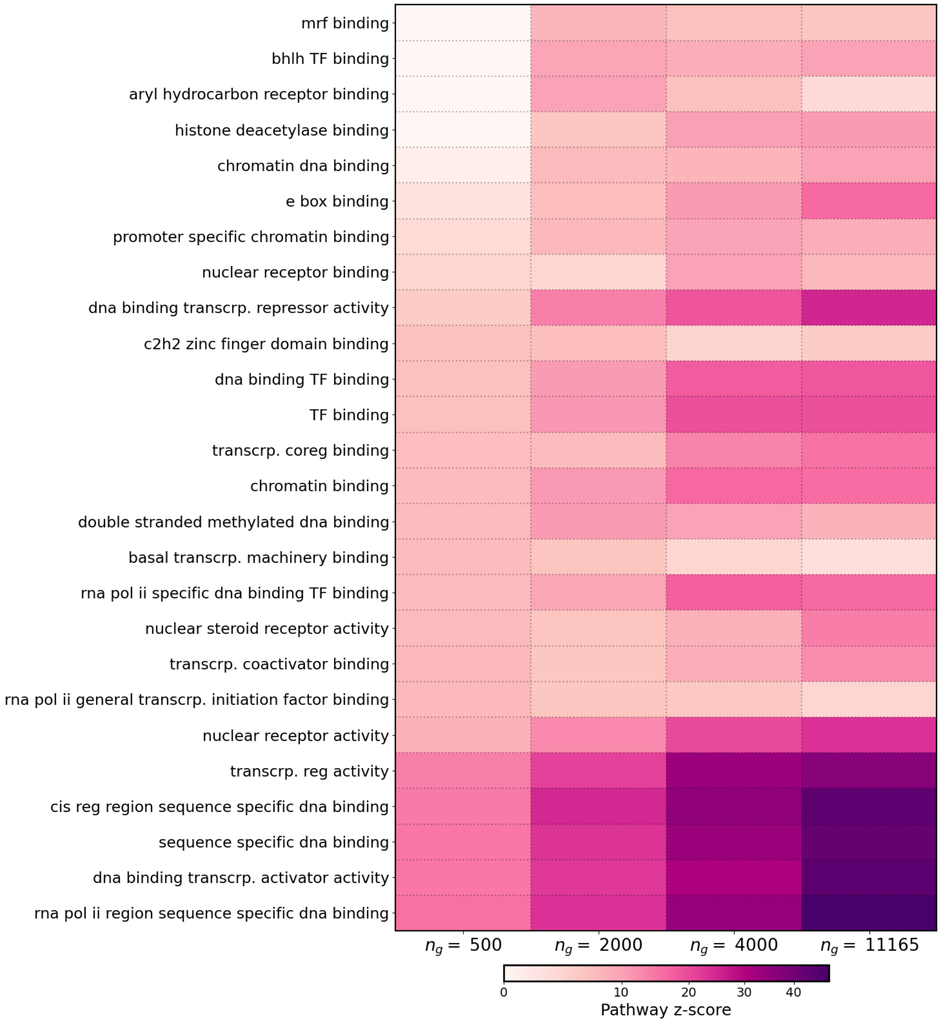

**Supp. Fig. 9** We applied PHOENIX to a pseudotrajectory of 186 breast cancer samples (ordered along subsequent “pseudotimepoints”) consisting of $n_g = 11165$ genes [17]. We also repeated the analysis on smaller subsets of genes $n_g = 500, 2000, 4000$, where we subsetted the full trajectory to only the $n_g$ most variable genes in the pseudotrajectory. We used the trained PHOENIX models to extract influence scores for pathways in the **Gene Ontology (molecular function)** database (see Methods 3.4), and visualized influence scores for the most central pathways across different values of $n_g$.

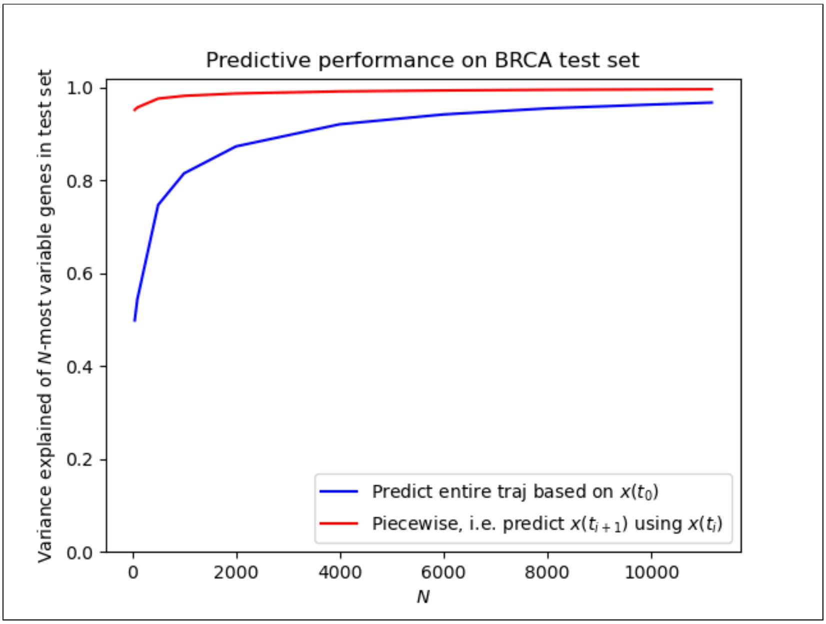

**Supp. Fig. 10** For the breast cancer data, we depict the $R^2$ performance (y-axis) of a PHOENIX model trained on all genes, when considering only the $N$ most variable genes (x-axis) for evaluation. We further evaluate the impact of predicting the entire trajectory based on the initial gene expression value ($t_0$, blue) versus predicting each expression value based on its immediately previous time point (red).

**Supp. Table 8** Detailed results of permutation tests (see Methods 3.4) used to obtain pathway influence scores from trained PHOENIX models and the **Reactome pathway database** [18]. The mean ($\mu_0$) and standard deviation ($\sigma_0$) of each permutation test null distribution, over $K = 1000$ permutations, are tabulated for each subset ($n_g$) in question, and the corresponding $z$-scores are visualized in **Figure 5**. Missing values indicate that the **all** genes involved in that pathway were excluded from that particular subset based on low variability in expression. Reactome accession IDs for all the pathways below have been provided in Supp. Table 9 for reproducibility.

| **Reactome pathway** | $n_g = 500$ genes | | | $n_g = 2000$ genes | | | $n_g = 4000$ genes | | | $n_g = 11165$ genes | | |
|---|---|---|---|---|---|---|---|---|---|---|---|---|
| | $z$ | $\mu_0$ | $\sigma_0$ | $z$ | $\mu_0$ | $\sigma_0$ | $z$ | $\mu_0$ | $\sigma_0$ | $z$ | $\mu_0$ | $\sigma_0$ |
| sumoylation of intracellular receptors | 8.001 | 0.008 | 0.007 | 8.055 | 0.003 | 0.002 | 11.678 | 0.009 | 0.003 | 13.834 | 0.003 | <9e-4 |
| nuclear receptor transcription pathway | 7.81 | 0.013 | 0.009 | 10.956 | 0.007 | 0.002 | 19.355 | 0.017 | 0.004 | 24.588 | 0.005 | <9e-4 |
| estrogen dependent gene expression | 6.927 | 0.03 | 0.014 | 8.144 | 0.011 | 0.003 | 15.315 | 0.015 | 0.005 | 15.5 | 0.008 | 0.001 |
| transcriptional regulation by RUNX2 | 6.751 | 0.009 | 0.007 | 5.59 | 0.01 | 0.003 | 7.645 | 0.018 | 0.005 | 6 | 0.01 | 0.001 |
| sumoylation of transcription factors | 5.967 | 0.002 | 0.004 | 7.791 | 0.002 | 0.001 | 3.754 | 0.004 | 0.002 | 5.944 | 0.002 | <9e-4 |
| RUNX1 reg. wnt signaling | 5.811 | 0.002 | 0.004 | 5.429 | 0.001 | 0.001 | 4.787 | 0.001 | 0.001 | 4.105 | <9e-4 | <9e-4 |
| ESR mediated signaling | 5.8 | 0.041 | 0.015 | 2.85 | 0.019 | 0.004 | 3.6 | 0.029 | 0.006 | 4.237 | 0.014 | 0.001 |
| RUNX1 reg. estrogen receptor mediated transcrip. | 5.692 | 0.003 | 0.004 | 5.518 | 0.001 | 0.001 | 5.649 | 0.001 | 0.001 | 4.138 | <9e-4 | <9e-4 |
| transcriptional regulation of testis differentiation | 5.618 | 0.004 | 0.004 | 4.317 | 0.003 | 0.001 | 7.503 | 0.006 | 0.003 | 3.375 | 0.001 | <9e-4 |
| regulation of RUNX2 expression and activity | 5.459 | 0.005 | 0.005 | 5.002 | 0.004 | 0.002 | 5.037 | 0.008 | 0.003 | 3.227 | 0.006 | 0.001 |
| nuclear signaling by ERBB4 | 5.371 | 0.008 | 0.007 | 2.436 | 0.004 | 0.002 | 1.46 | 0.01 | 0.004 | 0.731 | 0.003 | <9e-4 |
| RUNX2 reg. bone development | 4.671 | 0.002 | 0.004 | 6.85 | 0.003 | 0.002 | 8.44 | 0.006 | 0.003 | 7.674 | 0.003 | <9e-4 |
| NGF stimulated transcription | 4.238 | 0.007 | 0.006 | 5.484 | 0.007 | 0.002 | 6.593 | 0.01 | 0.004 | 9.107 | 0.003 | <9e-4 |
| transcriptional regulation of granulopoiesis | 1.948 | 0.002 | 0.004 | 5.556 | 0.003 | 0.001 | 9.688 | 0.008 | 0.003 | 13.092 | 0.003 | <9e-4 |
| estrogen dep. nucl. events (downstr. ESR signal) | 1.445 | 0.01 | 0.008 | 0.257 | 0.004 | 0.002 | 0.437 | 0.006 | 0.003 | 13.961 | 0.002 | <9e-4 |
| regulation of gene expression in beta cells | 1.065 | 0.002 | 0.004 | 2.342 | 0.001 | <9e-4 | 8.107 | 0.004 | 0.002 | 6.665 | 0.001 | <9e-4 |
| regulation of pten gene transcription | -0.307 | 0.003 | 0.004 | 5.367 | 0.001 | 0.001 | 7.716 | 0.009 | 0.004 | 4.028 | 0.005 | 0.001 |
| transcriptional regulation of pluripotent stem cells | -0.474 | 0.003 | 0.004 | 3.309 | 0.003 | 0.001 | 6.545 | 0.007 | 0.003 | 10.134 | 0.002 | <9e-4 |
| aryl hydrocarbon receptor signalling | | | | 9.454 | 0.001 | 0.001 | 6.065 | 0.001 | 0.001 | 2.482 | 0.001 | <9e-4 |
| RUNX2 reg. chondrocyte maturation | | | | 7.833 | 0.001 | 0.001 | 6.172 | 0.001 | 0.001 | 4.482 | <9e-4 | <9e-4 |
| Xenobiotics | | | | 6.516 | 0.001 | 0.001 | 2.456 | 0.005 | 0.003 | 2.145 | 0.001 | <9e-4 |
| GLI proteins bind promoters of Hedgehog | | | | 6.378 | 0.001 | 0.001 | 8.154 | 0.003 | 0.002 | 6.22 | <9e-4 | <9e-4 |
| RUNX3 reg. CDKN1A transcription | | | | 5.776 | 0.001 | 0.001 | 2.117 | 0.001 | 0.001 | 5.43 | 0.001 | <9e-4 |
| suppression of apoptosis | | | | -0.04 | 0.001 | 0.001 | 0.094 | 0.001 | 0.001 | 10.147 | 0.001 | <9e-4 |
| myogenesis | | | | -0.223 | 0.001 | 0.001 | 1.283 | 0.004 | 0.002 | 7.772 | 0.002 | <9e-4 |
| TP53 reg. apoptosis | | | | -0.295 | 0.002 | 0.001 | 4.796 | 0.004 | 0.002 | 13.769 | 0.001 | <9e-4 |
| TP53 reg. transcrip. of caspase activators | | | | | | | 8.13 | 0.001 | 0.001 | 3.15 | 0.001 | <9e-4 |
| MECP2 reg. transcription factors | | | | | | | 4.4 | 0.001 | 0.001 | 9.114 | <9e-4 | <9e-4 |

**Supp. Table 9** Accession IDs for Reactom pathways described in **Figure 5** (main paper) and Supp. Table 8. We shortened the names of pathways to fit them into tables and figures throughout the paper and supplement. Hence we're providing the accession IDs here.

| **Pathway** | **Reactome Stable Identifier** |
|---|---|
| Sumoylation of intracellular receptors | R-HSA-4090294 |
| Nuclear receptor transcription pathway | R-HSA-383280 |
| Estrogen dependent gene expression | R-HSA-9018519 |
| Transcriptional regulation by RUNX2 | R-HSA-8878166 |
| Sumoylation of transcription factors | R-HSA-3232118 |
| RUNX1 reg. wnt signaling | R-HSA-195721 |
| ESR mediated signaling | R-HSA-8939211 |
| RUNX1 reg. estrogen receptor mediated transcrip. | R-HSA-8931987 |
| Transcriptional regulation of testis differentiation | R-HSA-9690406 |
| Regulation of RUNX2 expression and activity | R-HSA-8939902 |
| Nuclear signaling by ERBB4 | R-HSA-1251985 |
| RUNX2 regulates bone development (corrected) | R-HSA-8941326 |
| NGF stimulated transcription | R-HSA-9031628 |
| Transcriptional regulation of granulopoiesis | R-HSA-9616222 |
| Estrogen dep. nucl. events (downstr. ESR signal) | R-HSA-9634638 |
| Regulation of gene expression in beta cells | R-HSA-210745 |
| Regulation of pten gene transcription | R-HSA-8943724 |
| Transcriptional regulation of pluripotent stem cells | R-HSA-452723 |
| Aryl hydrocarbon receptor signaling | R-HSA-8937144 |
| RUNX2 reg. chondrocyte maturation | R-HSA-8941284 |
| Xenobiotics | R-HSA-211981 |
| GLI proteins bind promoters of Hedgehog | R-HSA-5635851 |
| RUNX3 reg. CDKN1A transcription | R-HSA-8941855 |
| Suppression of apoptosis | R-HSA-9635465 |
| Myogenesis | R-HSA-525793 |
| TP53 reg. apoptosis | R-HSA-5633008 |
| TP53 reg. transcrip. of caspase activators | R-HSA-6803207 |
| MECP2 reg. transcription factors | R-HSA-9022707 |

**Supp. Table 10** Additional performance metrics for PHOENIX on breast cancer data

| Number of genes ($n_g$)[1] | Runtime (hrs)[2] | Concordance with influential ChIP genes[3] |
|---|---|---|
| 500 | 0.10 | 11.48% |
| 2000 | 0.14 | 18.03% |
| 4000 | 0.50 | 31.15% |
| 11165 | 2.52 | 80.33% |

[1]The full data set consisted of $n_g = 11165$ genes [17]. We also fit PHOENIX to smaller subsets of genes $n_g = 500, 2000, 4000$, where we subsetted the full data set to only the $n_g$ most variable genes in the pseudotrajectory.

[2]Runtime for a single run when using an AWS c5.4xlarge instance ($0.68/hour).

[3]For each $n_g$, we computed inferred influence scores for individual genes based on perturbation analyses on the fitted PHOENIX model (see Methods 3.3). We also computed harmonic centralities of the genes based on a validation network describing ChIP-binding data [3]. We then did a binary assignment, where the top 10% ($10\% \times n_g$) genes with highest inferred influence were labelled "predicted influential" and the remaining ($90\% \times n_g$) were "predicted non-influential." We measured concordance as the **sensitivity** $\frac{TP}{TP+FN}$ with which these predicted labels recovered the "truly influential" genes (those genes with non-zero harmonic centrality in the ChIP validation network).

**Supp. Table 11** We compared the performance of PHOENIX to other methods of regulatory dynamics estimation on a pseudotrajectory of 186 breast cancer samples (ordered along subsequent "pseudotimepoints") consisting of $n_g = 11165$ genes [17]. The data was processed for model fitting via the steps described in Methods 3.2. Snapshot-based methods (Dynamo, RNA-ODE, Deepvelo) require RNA velocity at every time point as an additional input [7–9]. Given that this information was not available in the data set, we estimated RNA velocity using a method of finite differences applied to smooth splines through the expression trajectories [19] (see Supp. Methods 3.1). Once each model was trained, we measured predictive accuracy in a step-wise fashion; for $N$ ranging from 50 to 11165, we calculated the **test set MSEs** (as described in Methods 3.2) of only the top-$N$ most varying genes in the dataset.

| $N$[1] | Dynamo | RNA-ODE | PHOENIX | DeepVelo | OOTB[2] |
|---|---|---|---|---|---|
| 50 | 1.61E-02 | 1.60E-02 | **1.57E-02** | 1.68E-02 | 1.59E-02 |
| 100 | 1.22E-02 | 1.21E-02 | **1.18E-02** | 1.20E-02 | 1.20E-02 |
| 500 | 5.76E-03 | 5.73E-03 | 5.62E-03 | 5.81E-03 | **5.54E-03** |
| 1000 | 4.04E-03 | 4.02E-03 | 3.95E-03 | 4.95E-03 | **3.90E-03** |
| 2000 | 2.78E-03 | 2.77E-03 | **2.72E-03** | 2.83E-03 | 2.79E-03 |
| 4000 | 1.79E-03 | 1.78E-03 | **1.75E-03** | **1.75E-03** | 1.77E-03 |
| 6000 | 1.32E-03 | 1.32E-03 | **1.29E-03** | 1.30E-03 | 1.30E-03 |
| 8000 | 1.05E-03 | 1.04E-03 | **1.02E-03** | 1.07E-03 | 1.03E-03 |
| 10000 | 8.58E-04 | 8.56E-04 | **8.39E-04** | 8.44E-04 | 8.43E-04 |
| 11165 | 7.73E-04 | 7.70E-04 | **7.55E-04** | 7.76E-04 | 7.59E-04 |

[1]$N$ = Number of most highly variable genes being considered in the MSE calculation.

[2]OOTB = Out-of-the-box NeuralODE, resembling how plain NeuralODEs are typically used for this problem [4, 5].

### Supp. Results 7 PHOENIX on B-cell RNASeq

**Supp. Table 12** Top 50 most changing regulators discovered by PHOENIX in B cells after Rituximab treatment as measured by log-fold change.

| Gene Symbol | log-fold change | Gene Symbol | log-fold change |
|---|---|---|---|
| PNRC2 | 2.56873290336475 | PPP2R3C | 2.47535285573376 |
| PCDHAC2 | 2.4183531041133 | PSD | -2.33736834391816 |
| SDF2L1 | -2.32960158182622 | PHLPP2 | -2.31911018375466 |
| TTC27 | 2.31393595893796 | SLFN12 | 2.31373381881861 |
| HMOX2 | 2.29844754939876 | ARPC5L | 2.28379510626964 |
| EXOSC9 | 2.27076773960713 | CCT6A | 2.26711838782967 |
| CLUH | -2.26276854159785 | NACC2 | -2.25002672061429 |
| CCDC51 | 2.23323817802628 | ACAA2 | 2.23302148320103 |
| RNF10 | 2.2271584454618 | CCNT1 | 2.22571736590982 |
| GRAP | -2.2202087315781 | TIMM8B | -2.20914724136027 |
| GOLGA4 | 2.20154634635607 | DTNBP1 | 2.16880863093615 |
| SETBP1 | -2.16869943368544 | GNG10 | -2.16748492186343 |
| ASL | 2.16732773542853 | HNRNPA1L2 | -2.16278244412603 |
| MRPS5 | -2.15142782602203 | COA6 | 2.15119270022043 |
| NOL8 | -2.14932979658522 | NKAP | 2.14646756016082 |
| SMYD4 | 2.14241657408929 | TMEM181 | 2.14176930774647 |
| PPM1K | 2.14122503727424 | TMEM43 | -2.13977794610878 |
| ZC3H4 | 2.13435569243602 | UPRT | -2.12251533684904 |
| TMEM9B | -2.12219325230208 | WIPI1 | -2.11544001910886 |
| ATP1A1 | -2.10439941185222 | ISCU | -2.10262448389575 |
| TOX4 | 2.10142807192934 | ABI2 | -2.09719154828222 |
| AGGF1 | 2.09364544925298 | KTI12 | -2.09231153185146 |
| CHRDL2 | -2.08727487121291 | HACL1 | -2.0871444586354 |
| CNTN4 | -2.08664483437801 | MAML1 | -2.08413955022521 |
| RAB1A | -2.08379529871721 | MFNG | 2.07967407563063 |

**Supp. Table 13** Training results from PHOENIX models applied to the B-cell RNASeq datasets. We fitted two seperate PHOENIX models, one for each condition of untreated and treated with Rituximab.

| Condition | validation set MSE | training set $R^2$ |
|---|---|---|
| Untreated | 0.0015 | 91.77% |
| Treated | 0.0021 | 89.58% |

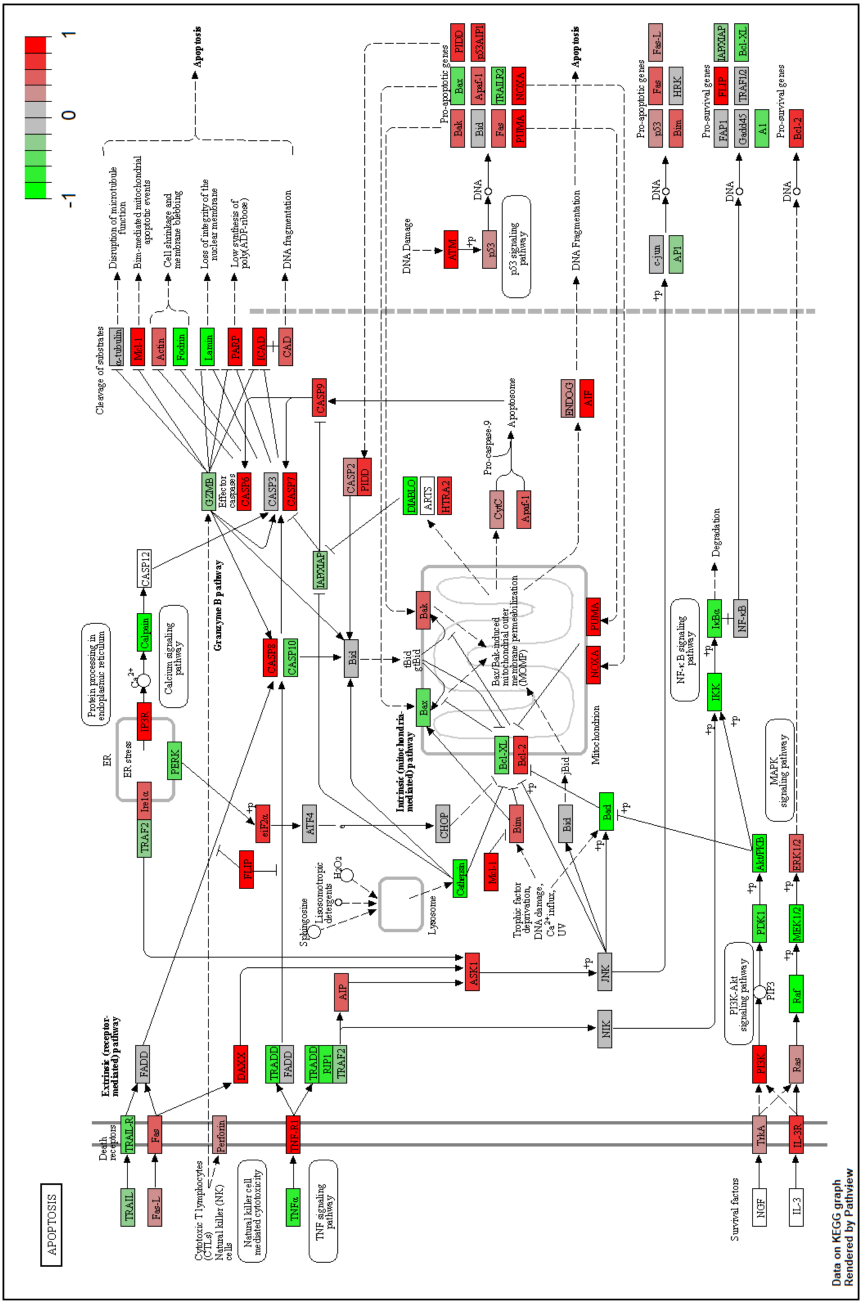

**Supp. Fig. 11** Regulatory changes discovered by PHOENIX in B cells after Rituximab treatment. All genes annotated in the KEGG Apoptosis pathway are colored by log-fold change between treatment and control, the range of values has been normalized to $[-1, 1]$. Positive values/red color shades mean that these genes show higher regulatory influence in Rituximab-treated B cells compared to control.

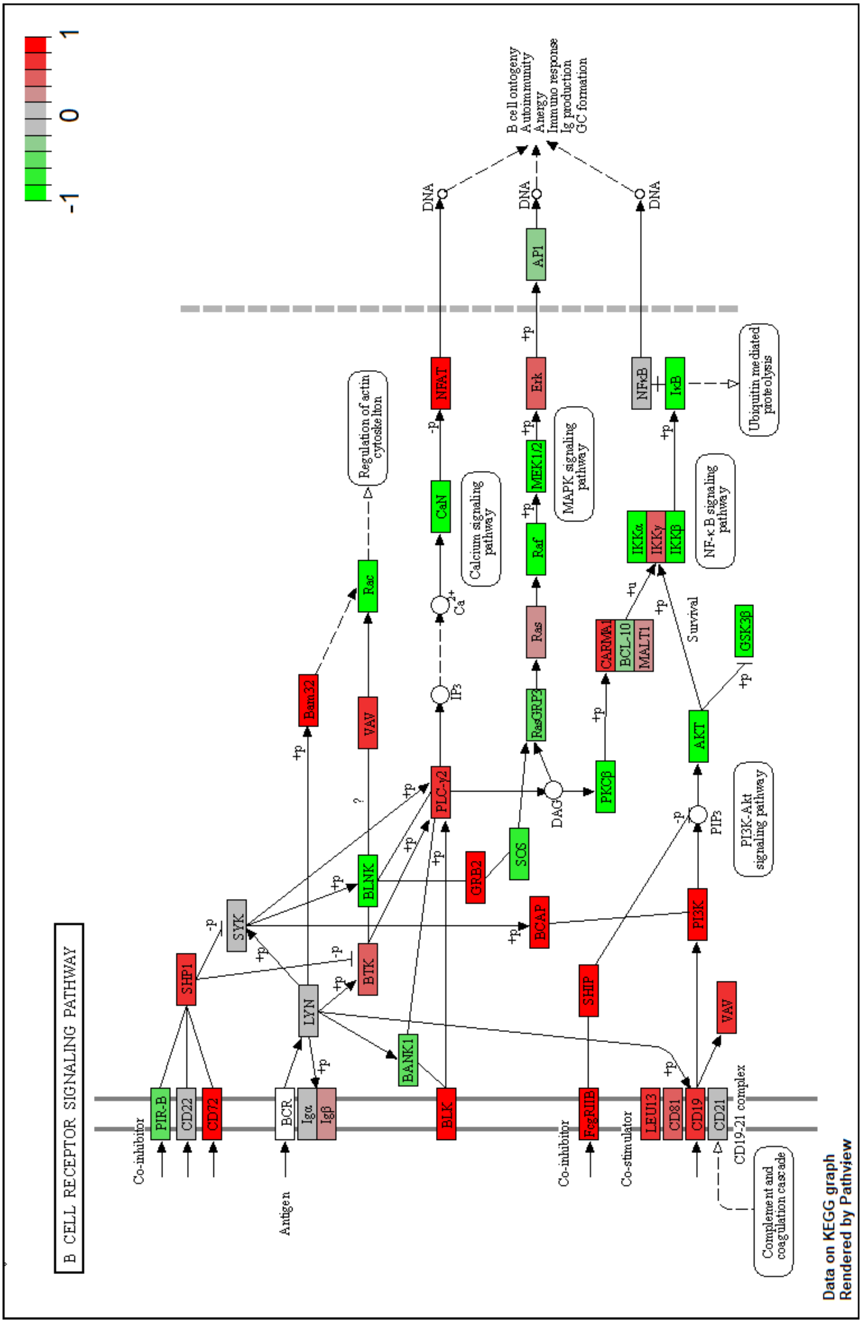

**Supp. Fig. 12** Regulatory changes discovered by PHOENIX in B cells after Rituximab treatment. All genes annotated in the KEGG B cell receptor signaling pathway are colored by log-fold change between treatment and control, the range of values has been normalized to $[-1, 1]$. Positive values/red color shades mean that these genes show higher regulatory influence in Rituximab-treated B cells compared to control.

### Supplemental methods

### Supp. Methods 1 PyTorch: details

#### Supp. Methods 1.1 NN architecture and forward()

Both $NN_{sums}$ and $NN_{prods}$ are fully connected single layer NNs with Hill-like activations that take $n$ inputs (for $n$ genes) and produce $m$ outputs each.

```
(NN_sums): Sequential(
    (activation_0): phi_Sigma()
    (sum_combos): Linear(in_features=n,out_features=m, bias = True))

(NN_prods): Sequential(
    (activation_0): phi_Pi()
    (prod_combos): Linear(in_features=n,out_features=m, bias = True))
```

We use $m << n$ allowing us to approximate lower dimensional sets of additive and multiplicative gene combinations. The choice of $m$ is determined by:

1. We surmised that $m$ should roughly scale with $n$
2. Given a value of $n$, we selected a plausible range of values for $m$, and picked $m$ based on the predictive performance on the validation set. This allows for a means to balance between fitting accuracy and risk of overfitting.

This lead to the following size $m$ in each experiment, where $m$ roughly scaled with the dimensionality $n$ of the problem,

- Simulations: For ($n = 350, 690$ genes) we use $m = 40, 50$, respectively
- Yeast cell cycle: There were $n = 3551$ genes, and we used $m = 120$
- Breast cancer: For subset sizes $n_g = 500, 2000, 4000, 11165$ genes we used $m = 40, 100, 120, 300$ respectively
- B-cell RNASeq: There were $n = 14691$ genes, and we found used $m = 200$ to give the best validation performance.

These $2m$ outputs are then fed into a final fully connected single layer $NN_{combine}$, and gene-specific parameters ($\boldsymbol{v}$/ `u_vector` $\in \mathbb{R}^n$) are applied to create the `final` estimations for the $n$ local derivatives.

```
(NN_combine): Sequential(
    (final_outputs): Linear(in_features=2*m,out_features=n))

def forward(self, t, input):
    c_Sigma = self.NN_sums(input)
    c_Pi = torch.exp(self.NN_prods(input))
    joint = self.NN_combine(torch.cat((c_Sigma, c_Pi),dim=-1)
    final = torch.relu(u_vector)*(joint - y)
    return(final)
```

#### Supp. Methods 1.2 Predictive performance: training, testing, and validation (choosing $\boldsymbol{\lambda}$)

We re-shaped the simulated data sets to be amenable with the `torchdiffeq` package in `PyTorch` [20], which contains the base library used for NeuralODEs. Regardless of whether the data was split based on trajectories (*in silico* experiments) or transition pairs (real data applications), *at any given training step* the data was fed into the PHOENIX model in pairwise-vector form. For instance, in the *in silico* experiments, each training trajectory consisted of 5 time-points $\in \{0, 2, 3, 7, 9\}$, and was subsequently fed to the model in the form of its 4 constituent transition pairs $(t_i, t_{i+1}) \in \{(0, 2), (2, 3), (3, 7), (7, 9)\}$, where a transition pair consists of two consecutive expression vectors in the trajectory $(\boldsymbol{g}(t_i),\ \boldsymbol{g}(t_{i+1}))$. With each transition pair fed to the model, it learned an approximation of the derivative that described the transition of $\boldsymbol{g}(t_i)$ to $\boldsymbol{g}(t_{i+1})$. We initialized each of $\text{NN}_{sums}$, $\text{NN}_{prods}$, and $\text{NN}_{combine}$ with a sparse initialization scheme that set 95% of the weight parameters to be 0. We initialized the $\boldsymbol{v}_i$s with i.i.d standard uniform values.

We used `dopri5` (included with `torchdiffeq`) as the ODESolver within PHOENIX's NeuralODE engine, and the Adam optimizer (also included with `torchdiffeq`) to optimize NN parameters. Broadly speaking, `torchdiffeq` learns by performing back propagation through the ODESolver via adjoint sensitivity analysis [21]. Next, we use a `simulated_batch` of expression values $\{\boldsymbol{\gamma_k}\}_{k=1}^{K}$ to incorporate a structural domain knowledge inspired prior model. Since the $\boldsymbol{\gamma_k}$s were simulated expression values, we also refer to them as "ghost" expression values.

$$\mathcal{P}^*\left(\boldsymbol{\gamma_k}\right) + \boldsymbol{\gamma_k} = \boldsymbol{A} \cdot \boldsymbol{\gamma_k}$$

So, given the user-supplied `prior_matrix` $\boldsymbol{A}$, we pre-calculated `prior_output`, $\mathcal{P}^*\left(\boldsymbol{\gamma_k}\right) + \boldsymbol{\gamma_k}$, using `torch.matmul()`.

```
simulated_batch = torch.rand(10000,1,prior_mat.shape[0])
prior_output = torch.matmul(batch_for_prior,prior_mat)
```

We tied this into model training using a modified loss function `composed_loss` with weight ($\lambda$). We obtained model `predictions` using the `odeint()` function in `torchdiffeq`, as shown in the pseudocode below:

```
def training_step(...., training_batch, target, simulated_batch,
                    prior_output, lambda, time_pts):
    predictions =  odeint(....,training_batch, time_pts)
    loss_data = torch.mean((predictions - target)**2)
    model_sim_output = forward(time_pts,simulated_batch) +
                                                  simulated_batch
    loss_prior = torch.mean((model_sim_output - prior_output)**2)
```

```
    composed_loss = lambda * loss_data +
                    (1- lambda) * loss_prior
    composed_loss.backward()
    opt.step()
    return [loss_data, loss_prior]
```

We trained for up to 200 epochs on an AWS c5.9xlarge instance, where each epoch consisted of the entire training set being fed to the model in the form of constituent transition pairs. We used `torch.optim` function `ReduceLROnPlateau()` to reduce the learning rate by 10% every 3 epochs, unless the validation set performance showed reasonable improvement.

```
scheduler = optim.lr_scheduler.ReduceLROnPlateau(opt, mode='min',
            factor=0.9, patience=3, threshold=1e-05,
            threshold_mode='abs', eps=1e-09)
```

Training terminated if validation set performance failed to improve in 10 consecutive epochs. We repeated this entire pipeline for a grid of $\lambda \in \{0.1, 0.2, 0.5, 0.8, 0.9, 0.99, 0.999, 1\}$, and used the validation mean squared error (MSE) at model termination to decide on an optimal $\lambda$. For this final model, we evaluated final predictive performance using test set MSE.

For even further details (exact learning rates, `torchdiffeq` details, ODE-Solver details, optimizer details, etc.) we refer the reader to our GitHub repository [1], where the entire code base is made available.

### Supp. Methods 2 Explainability performance: GRN inference

#### Supp. Methods 2.1 Algorithm for efficiently retrieving encoded GRN from trained PHOENIX model

We start with PHOENIX's prediction for the local derivative given a gene expression vector $\boldsymbol{g}(t) \in \mathbb{R}^n$ in an $n$-gene system:

$$\widehat{\frac{d\boldsymbol{g}(t)}{dt}} = \text{ReLU}(\boldsymbol{\upsilon}) \odot \Big[\boldsymbol{W}_\cup \{\boldsymbol{c}_{\boldsymbol{\Sigma}}(\boldsymbol{g}(t)) \oplus \boldsymbol{c}_{\boldsymbol{\Pi}}(\boldsymbol{g}(t))\} - \boldsymbol{g}(t)\Big], \quad \text{where}$$

$$\boldsymbol{c}_{\boldsymbol{\Sigma}}(\boldsymbol{g}(t)) = \boldsymbol{W}_{\boldsymbol{\Sigma}} \phi_\Sigma(\boldsymbol{g}(t)) + \boldsymbol{b}_{\boldsymbol{\Sigma}} \quad \text{and} \quad \boldsymbol{c}_{\boldsymbol{\Pi}}(\boldsymbol{g}(t)) = \exp \circ (\boldsymbol{W}_{\boldsymbol{\Pi}} \phi_\Pi(\boldsymbol{g}(t)) + \boldsymbol{b}_{\boldsymbol{\Pi}})$$

We observed that a trained PHOENIX model encodes interactions *between* genes primarily within the gene-specific multipliers $\boldsymbol{\upsilon} \in \mathbb{R}^n$, and the weight parameters from its neural network blocks $\boldsymbol{W}_{\boldsymbol{\Pi}}, \boldsymbol{W}_{\boldsymbol{\Sigma}} \in \mathbb{R}^{m \times n}$ and $\boldsymbol{W}_\cup \in \mathbb{R}^{n \times 2m}$. This inspired an efficient means of projecting the estimated dynamical system down to a gene regulatory network (GRN) $\widehat{G_n}$.

We first calculated a matrix $\boldsymbol{D} \in \mathbb{R}^{n \times n}$, where $\boldsymbol{D}_{ij}$ approximated the *absolute contribution* of gene $j$ to the derivative of gene $i$'s expression:

$$\boldsymbol{D} = \boldsymbol{W}_\cup \begin{bmatrix} \boldsymbol{W}_{\boldsymbol{\Sigma}} \\ \boldsymbol{W}_{\boldsymbol{\Pi}} \end{bmatrix}$$

We applied the gene-specific multipliers $\boldsymbol{\upsilon}$, before adapting the marginal attribution approach described by Hackett *et al.* [22]. This resulted in the **dynamics matrix** $\widetilde{\boldsymbol{D}}$ where $\widetilde{\boldsymbol{D}_{ij}}$ was scaled according to the *relative contribution* of gene $j$ to the rate of change in gene $i$'s expression:

$$\widetilde{\boldsymbol{D}_{ij}} = \frac{\boldsymbol{\upsilon}_i \boldsymbol{D}_{ij}}{\sum_{j'=1}^{n} |\boldsymbol{\upsilon}_i \boldsymbol{D}_{ij'}|}$$

Finally, we subjected $\widetilde{\boldsymbol{D}}$ to a cut-off value $v_{\mathcal{C}}$ based on an appropriate percentile $\mathcal{C}$; we used $\mathcal{C} = 0.995$, meaning $v_{\mathcal{C}} = 99.5^{th}$ percentile of $\{| \widetilde{\boldsymbol{D}_{ij}} |\}_{\forall i,j}$. Specifically, we decided edge **existence** and **strength** in $\widehat{G_n}$ as follows:

$$\textbf{No edge}\text{: gene } j \overset{0}{\to} \text{gene } i \iff | \widetilde{\boldsymbol{D}_{ij}} | \leq v_{\mathcal{C}}$$

$$\textbf{Activating edge}\text{: gene } j \overset{\widetilde{\boldsymbol{D}_{ij}}}{\to} \text{gene } i \iff \widetilde{\boldsymbol{D}_{ij}} > v_{\mathcal{C}}$$

$$\textbf{Repressive edge}\text{: gene } j \overset{\widetilde{\boldsymbol{D}_{ij}}}{\to} \text{gene } i \iff \widetilde{\boldsymbol{D}_{ij}} < -v_{\mathcal{C}}$$

##### Supp. Methods 2.2 Evaluation of explainability

We compared the estimated $\widehat{G_n}$ to the corresponding validation network in terms of out-degree correlation and edge-existence, calculating recovery AUC, true positive rate (classification TPR), and true negative rate (classification TNR). For our *in silico* experiments, we reverse-engineered a method to inform how sparsely PHOENIX had inferred the dynamics.

Since the ground truth graphs $G_{350}$ and $G_{690}$ were known, we found the value of $\mathcal{C} \in (0, 1)$, the aforementioned percentile cutoff of $\mid \widetilde{\boldsymbol{D}_{ij}} \mid$ values, that maximized the balanced classification accuracy ($\frac{\text{TPR}+\text{TNR}}{2}$), and used this $\mathcal{C}_{\max}$ as a measure of how sparsely the dynamics were inferred. Since $\mathcal{C}_{\max}$ reflects how well the trained PHOENIX model discriminates between true and non interactions, a low value of $\mathcal{C}_{\max}$ corresponded to dense dynamics, while a high value corresponded to sparser dynamics. When comparing explainability across different PHOENIX fits, we used $\mathcal{C}_{\max}$ to obtain corresponding values $\text{TPR}_{\max}$ and $\text{TNR}_{\max}$. This eliminated the dependence on any arbitrary cutoff and allowed us to compare networks in terms of best possible TPR and TNR.

#### Supp. Methods 3 Benchmark experiments against existing methods

##### Supp. Methods 3.1 Data sets used for benchmarking

For comparison to OOTB models, we used the same simulated data sets from SIM350 and SIM690 that were used for the PHOENIX experiments, with the same train-val-test split. For PRESCIENT [6], we found it to work only with consecutive integer time points, and so we generated 160 noisy trajectories from the ground-truth SIM350 and SIM690 at $t \in T = \{0, 1, 2, 3, 4, 5, 6, 7, 8, 9\}$. Each of the 1600 simulated vectors in $\{\{\boldsymbol{g}(t)_k \in \mathbb{R}^n\}_{t \in T}\}_{k=1}^{160}$ could be conceptualized as a **single cell**, with `cell_id` $= (t, k)$, at time $t$ with expression values for $n$ genes. We partitioned the cells via the same train-val-test split as the PHOENIX runs, and used this time-labelled "single cell training data" as PRESCIENT's input. We set all `cell_type` to `regular` in the meta data.

For "two-step" methods such as Dynamo [7], RNA-ODE [8], and DeepVelo [9], we know that they estimate dynamics by first reconstructing RNA velocity using inputs such as spliced and unspliced mRNA counts (step 1) and then estimating a vector field mapping expression to velocity (step 2). To avoid the need for uncommon input data types and to also emulate the **theoretically optimal performance** in step 1, we used the noiseless ground truth velocities $\frac{d\boldsymbol{g}(t)}{dt}$ as input into step 2 directly, since these true velocities could be retrieved from our simulator based on the random seeds used. Subsequently, we only tested step 2 (with the same train, validation, and test data sets used in PHOENIX and the corresponding ground truth velocities) and obtained an optimistic estimate of each method's performance.

As just discussed above, Dynamo, RNA-ODE, and DeepVelo are "two-step" snapshot based methods that require RNA velocity at every time point as an additional input [7–9]. Given that this information was not available in either the yeast or the breast cancer data set, we estimated RNA velocity using a method of finite differences applied to smooth splines through the expression trajectories [19], in order to apply these methods to those datasets. Specifically, for each gene in the dataset, we used the local estimator of scatterplot smoothing function `loess()` in `R` to fit a smooth spline through its expression trajectory (degree = 2, smoothing parameter $\alpha = 2$). Then we simply calculated the local derivative w.r.t. time on this smooth spline using `diff(smoooth_y)/diff(t)` in `R`. Since this approach calculates a backward finite difference, we manually copied the derivative at $t_2$ on to $t_1$.

#### Supp. Methods 3.2 Out-of-the-box NeuralODE models

We tried to emulate how one might typically use OOTB NeuralODE models for the purpose of predicting gene expression dynamics [4]. Given a gene regulatory network of $n$ genes, we assume that the gene expression of all genes $g_j(t)$ can have an effect on a specific $g_i(t)$: $\frac{d\boldsymbol{g}(t)}{dt} = f_{reg}\Big(\boldsymbol{g}(t)\Big) - \boldsymbol{g}(t)$, where $\boldsymbol{g}(t) = \{g_i(t)\}_{i=1}^n$ and $f_{reg} : \mathbb{R}^n \to \mathbb{R}^n$. We approximated $f_{reg}$ with just an out-of-the-box neural network ($\text{NN}_{OOTB}$) with parameters $\boldsymbol{\theta}_{OOTB}$, and ReLU, tanh, or sigmoid activation functions:

$$\frac{d\boldsymbol{g}(t)}{dt} \approx \text{NN}_{OOTB}\Big(\boldsymbol{g}(t), \boldsymbol{\theta}_{OOTB}\Big) - \boldsymbol{g}(t)$$

For fair comparison, we created $\text{NN}_{OOTB}$ such that it contained a similar number of hidden layers and trainable parameters as PHOENIX. Specifically, we designed (`basic_act_funcs` used were Sigmoid, tanh, and ReLU) as follows:

```
(NN_OOTB): Sequential(
    (activation_0): basic_act_func()
    (layer_1): Linear(in_features=n, out_features=2*m, bias=True)
    (activation_1): basic_act_func()
    (layer_out): Linear(in_features=2*m, out_features=n), bias=True)

def forward(self, t, input):
    res = self.NN_OOTB(input)
    return(res - y)
```

We chose $m = 40$ for $n = 350$ (SIM350), and $m = 50$ for $n = 690$ (SIM690). We used the same basic learning strategy as the experiments with PHOENIX. These details as well as other technicalities are on our GitHub repository [1].

To evaluate explainability, we bootstrapped the trained $\text{NN}_{OOTB}$ to estimate the encoded GRN. We randomly generated 100 input expression vectors via i.i.d standard uniform sampling $\{\boldsymbol{b_k} \in \mathbb{R}^n\}_{k=1}^{100}$. Next, for each gene $j$, we created a perturbed version of these input vectors $\{\boldsymbol{b_k^j}\}_{k=1}^{100}$, where only gene $j$ was perturbed in each vector. We then fed both sets of input vectors into the trained $\text{NN}_{OOTB}$ to obtain corresponding perturbed output $\{\widehat{\boldsymbol{o}}_{\boldsymbol{k}}^{\boldsymbol{j}} \in \mathbb{R}^{n_g}\}_{k=1}^{200}$ and unperturbed output $\{\widehat{\boldsymbol{o}}_{\boldsymbol{k}} \in \mathbb{R}^{n_g}\}_{k=1}^{200}$. Next, for each perturbed gene $j$ in the input, we measured how much the perturbed output of every other gene $i$ changed, *as a proportion* of its unperturbed output. This generally yielded better results in favor of the OOTB models than if we proceeded without taking proportions, or normalized based on input perturbation.

$$\boldsymbol{\Delta}_{ij} = \frac{1}{200}\sum_{k=1}^{100}\left|\frac{\widehat{\boldsymbol{o}}_{\boldsymbol{k}i}^{\boldsymbol{j}} - \widehat{\boldsymbol{o}}_i}{\widehat{\boldsymbol{o}}_i}\right|$$

We used $\boldsymbol{\Delta}$ to create the normalized effects matrix $\widetilde{\boldsymbol{E}}$ such that $\widetilde{\boldsymbol{E}}_{ij}$ reflected the *relative contribution* of gene $j$'s input to the proportion change in gene $i$'s output: $\widetilde{\boldsymbol{E}}_{ij} = \frac{\boldsymbol{\Delta}_{ij}}{\sum_{j=1}^{n}\boldsymbol{\Delta}_{ij}}$. Finally, by thresholding the values of $\widetilde{\boldsymbol{E}}$ using an optimal cut off $\mathcal{C}_{\max}$ (see Supp. Methods 2.2; this is the the percentile cut off of $\widetilde{\boldsymbol{E}}_{ij}$ values that maximizes $\frac{TPR+TNR}{2}$), we obtained adjacency matrices describing $\widehat{G_{350}}$ and $\widehat{G_{690}}$ that approximated the ground truth $G_{350}$ and $G_{690}$, respectively.

#### Supp. Methods 3.3 Other benchmarked methods

We provide some detail here, but source code (and even more technicalities) can be found on our GitHub repository [1], where this entire benchmarking pipeline is made available. For assessing explainability in each case, we only explain up to how a matrix $\widetilde{\boldsymbol{E}}$ was obtained. We then processed $\widetilde{\boldsymbol{E}}$ in the same way as described at the end of Supp. Methods 3.2 to obtain $\widehat{G_n}$. We calculated $\widehat{G_n}$'s average out-degree and obtained AUC by comparing to ground truth $G_n$.

##### Supp. Methods 3.3.1 PRESCIENT

PRESCIENT [6] uses time-series scRNA-seq and cell-growth rate data to learn a scalar-valued potential function $\Psi(\boldsymbol{g}(t))$ with a neural network. A final drift model is obtained using automatic differentiation $\frac{\boldsymbol{g}(t)}{dt} = -\nabla\Psi(\boldsymbol{g}(t))$. We found that PRESCIENT's implementation could train only with consecutive integer time points, and so we generated 160 noisy trajectories from the ground-truth SIM350 and SIM690 at $t \in T = \{0, 1, 2, 3, 4, 5, 6, 7, 8, 9\}$. Each of the 1600 simulated vectors in $\{\{\boldsymbol{g}(t)_k \in \mathbb{R}^n\}_{t\in T}\}_{k=1}^{160}$ could be conceptualized as a **single cell**, with `cell_id` $= (t, k)$, at time $t$ with expression values for $n$ genes. We partitioned the cells via the same train-val-test split as the PHOENIX runs, and used this time-labelled "single cell training data" as PRESCIENT's input. PRESCIENT requires a meta data file mapping each `cell_id` to its timepoint $\in T = \{0, 1, 2, 3, 4, 5, 6, 7, 8, 9\}$. We also set the `cell_type` column in the meta-data file to be `regular` for all cells. PRESCIENT models can incorporate proliferation during training via computing a ”growth weight” per cell using either lineage tracing data or KEGG gene signatures. In the absence of such data in our simulation, we set this weight to be 1 for all cells. We used the `prescient` package in `Python` to train a model with a single hidden layer of width $k_{dim}$ followed by a softplus activation function. We trained for 2600 epochs and predicted trajectories using PRESCIENT's `net._drift()` function (that returns $\frac{\boldsymbol{g}(t)}{dt}$) along with `solve_ivp()` from `scipy.integrate`. We optimized $k_{dim}$ based on validation trajectory prediction. We did not find a straightforward functionality in the package to extract a GRN. Because the PRESCIENT implementation was limited to consecutive integer (or equally spaced) time points, this was an impediment to benchmarking PRESCIENT on the yeast and breast cancer datasets.

#### Supp. Methods 3.3.2 Dynamo

Given the input data (simulated gene expression and the corresponding ground truth RNA velocities), Dynamo [7] fits a sparse vector field mapping gene expression to RNA velocity using Gaussian kernel regression. Using the training trajectories from the input data, we created an `vf.SvcVectorField()` object (from the `dynamo` package in `Python`). The trained model (mapping expression to velocity) was used along with `solve_ivp()` from `scipy.integrate` to predict trajectories. Cross-validation was used (based on best MSE in validation trajectory prediction) to choose hyper-parameters $M$ (number of control points) and sparsity parameter $\lambda$. MSE was calculated for final model on the test trajectories.

Although `dynamo` provides a straighforward Jacobian function `get_Jacobian()`, the extraction of a GRN using this function has to be done carefully. This is because there is a possibility that the Jacobian is close to zero for both low and high regulator levels; the absolute value of a Jacobian is significantly larger than 0 only within a small range of expression values. Hence we used the following strategy to estimate a GRN:

- We sampled random expression vectors $\boldsymbol{g}_k$ from $\mathcal{N}(0.5, 0.25)$. Since high and low expression values corresponded to 1 and 0 in our simulations, sampling from this Gaussian centered around 0.5 made it less likely that we would be picking very high or low regulator levels.
- To further prevent the Jacobians from being too small, we did sampling in two phases:
    1. We first sampled 5000 random expression vectors $\{\boldsymbol{g}_k \in \mathbb{R}^n\}_{k=1}^{5000}$ from $\mathcal{N}(0.5, 0.25)$, and calculated the 5000 corresponding Jacobian matrices $\{\boldsymbol{J}_k \in \mathbb{R}^{n\times n}\}_{k=1}^{5000}$ using the `get_Jacobian()` function. We computed $\{\mid \boldsymbol{J}_k \mid_{\mathcal{F}} \in \mathbb{R}\}_{k=1}^{5000}$, the Frobenius norms for all the Jacobians. Then we chose $f_{\text{cutoff}} = 95^{\text{th}}$ percentile across the 5000 values in $\{\mid \boldsymbol{J}_k \mid_{\mathcal{F}}\}_{k=1}^{5000}$ as a filtration criterion for the next step.

2. We sampled a **new batch** of 10000 random expression vectors $\{\boldsymbol{g}_l\}_{l=1}^{10000}$ from $\mathcal{N}(0.5, 0.25)$ and computed $\{\boldsymbol{J}_l\}_{l=1}^{10000}$. We **discarded** any $\boldsymbol{J}_l$ that was "not large enough," that is if $|\ \boldsymbol{J}_k\ |_{\mathcal{F}} < f_{\text{cutoff}}$, and kept sampling until we had 10000 $\boldsymbol{J}_l$s that satisfied this criterion.

- We used the final list of 10000 Jacobian matrices to compute an average Jacobian $\widetilde{\boldsymbol{E}}$.

##### Supp. Methods 3.3.3 RNA-ODE

Given the input data (simulated gene expression and the corresponding ground truth RNA velocities), RNA-ODE [8] fits black-box random forests mapping expression to velocity. We obtained `Python` source code from `https://github.com/RuishanLiu/VelocytoAnalysis`, to build random forest regressors. We predicted trajectories using the `predict()` function of the random forest along with `solve_ivp()` from `scipy.integrate`. We optimized the number of trees in the forest (`n_estimators`) based on validation trajectory prediction. The source code also has a `GET_GRN()` function to easily estimate a $\widetilde{\boldsymbol{E}}$ based on the fitted model using a GENIE3-like approach.

##### Supp. Methods 3.3.4 DeepVelo

Given the input data (simulated gene expression and the corresponding ground truth RNA velocities), DeepVelo [9] fits a black-box autoencoder mapping expression to velocity. We obtained `Python` source code from `https://github.com/gersteinlab/DeepVelo`, to build a 4 layer encoder and 4 layer decoder with $\ell_1$ regularization. We used size $4p$ for the intermediate layers and size $p$ for the latent layers. We optimized $p$ (and hence the number of neurons in each layer of the autoencoder) based on validation trajectory prediction. Further details can be found on our GitHub repository [1]. We predicted trajectories using the auto-encoder's `predict()` function along with `solve_ivp()` from `scipy.integrate`. The source code also has functionalities to estimate $\widetilde{\boldsymbol{E}}$ a gene-correlation matrix of cells, based on simulating "retrograde trajectories."

### Supp. Methods 4 Creating *in silico* data

#### Supp. Methods 4.1 Ground truth system using SimulatorGRN

We created a ground truth gene regulatory network (GRN) by sampling from S. cerevisiae (yeast) regulatory networks obtained from the SynTReN v1.2 supplementary data in simple interaction format (SIF) [23]. The SynTReN file provides a directional GRN containing 690 genes and 1094 edges with annotations (activating vs repressive) for edge types; we defined this GRN to be ground truth network $G_{690}$. To obtain $G_{350}$, we used `SimulatorGRN` [24] in `R` to sample a subnetwork of 350 genes and 590 edges from $G_{690}$, using the `sampleGraph()` function. Next, to each edge (for instance from gene $A$ to gene $B$), we assigned randomly generated $EC_{50}^{AB} \in (0.4, 0.6)$ and $\eta^{AB} \in (1.39, 1.8)$ values using the `randomizePrams()` function from `SimulatorGRN`; lists of edges with $EC_{50}^{AB}, \eta^{AB}$ values are provided with the PHOENIX release [1]. Each $EC_{50}^{AB}, \eta^{AB}$ pair defines the relationship between a regulator $A$ and its target $B$. The `simulationGRN()` function then used the edges in $G_{350}$ and $G_{690}$ to define systems (SIM350 and SIM690) of normalised-Hill ODEs [25], where the activation of B by a single regulator A was modelled as:

$$\frac{dB}{dt} = f_{act}(A, EC_{50}^{AB}, \eta^{AB}) - B = \frac{\beta A^{\eta^{AB}}}{\beta - 1 + A^{\eta^{AB}}} - B,$$

$$\text{where } \beta = \frac{EC_{50}^{AB} - 1}{2EC_{50}^{AB} - 1}$$

Repression of $B$ by $A$ was simply modelled using $1 - f_{act}(A, EC_{50}^{AB}, \eta^{AB})$. Co-regulation by $A_1$ and $A_2$ was modelled as either a logical `AND`:

$$f_{act}(A_1, EC_{50}^{A_1B}, \eta^{A_1B}) \times f_{act}(A_2, EC_{50}^{A_2B}, \eta^{A_2B}),$$

or a logical `OR` gate:

$$f_{act}(A_1, EC_{50}^{A_1B}, \eta^{A_1B}) + f_{act}(A_2, EC_{50}^{A_2B}, \eta^{A_2B}) \\ - f_{act}(A_1, EC_{50}^{A_1B}, \eta^{A_1B}) \times f_{act}(A_2, EC_{50}^{A_2B}, \eta^{A_2B})$$

where we randomly assigned 90% of co-regulations as logical `AND`s, and the other 10% as logical `OR`s. Proceeding in this manner, `simulationGRN()` used the network structures to define dynamical systems SIM350 and SIM690, containing an ODE for each gene in each of $G_{350}$ and $G_{690}$ respectively. For any gene $Z$ that has no upstream regulators, we assign $dZ/dt = 0$. We simulated expression trajectories from these ODE systems using `R`'s `desolve` package.

#### Supp. Methods 4.2 Creating corrupted/misspecified prior models

For each noise level $\sigma_{\%} \in \{0\%, 5\%, 10\%, 20\%, 40\%, 80\%, 100\%\}$ in our *in silico* experiments, we created a shuffled version of $G_{350}$ (and similarly $G_{690}$) where we shuffled $\sigma_{\%}$ of the edges by relocating those edges to new randomly chosen origin and destination genes within the network. This yielded the shuffled network $G_{350}^{\sigma\%}$ (and similarly $G_{690}^{\sigma\%}$) with corresponding adjacency matrix $\boldsymbol{A}^{\boldsymbol{\sigma\%}}$. We set activating edges in $\boldsymbol{A}^{\boldsymbol{\sigma\%}}$ to +1 and repressive edges to -1, and defined the simple linear prior domain knowledge model: $\mathcal{P}^*(\boldsymbol{\gamma_k}) = \boldsymbol{A}^{\boldsymbol{\sigma\%}} \cdot \boldsymbol{\gamma_k} - \boldsymbol{\gamma_k}$.

### References

[1] Hossain, I. (2022). PHOENIX package [Computer software]. https://github.com/QuackenbushLab/phoenix

[2] Harbison, C. T., Gordon, D. B., Lee, T. I., Rinaldi, N. J., Macisaac, K. D., Danford, T. W., ... & Young, R. A. (2004). Transcriptional regulatory code of a eukaryotic genome. Nature, 431(7004), 99-104.

[3] Chèneby, J., Gheorghe, M., Artufel, M., Mathelier, A., & Ballester, B. (2018). ReMap 2018: an updated atlas of regulatory regions from an integrative analysis of DNA-binding ChIP-seq experiments. Nucleic acids research, 46(D1), D267-D275.

[4] Erbe, R., Stein-O'Brien, G., & Fertig, E. J. (2023). Transcriptomic forecasting with neural ordinary differential equations. Patterns (New York, N.Y.), 4(8), 100793. https://doi.org/10.1016/j.patter.2023.100793

[5] Hu, Y. (2022). Modeling the gene regulatory dynamics in neural differentiation with single cell data using a machine learning approach.

[6] Yeo, G. H. T., Saksena, S. D., & Gifford, D. K. (2021). Generative modeling of single-cell time series with PRESCIENT enables prediction of cell trajectories with interventions. Nature communications, 12(1), 1-12.

[7] Qiu, X., Zhang, Y., Martin-Rufino, J. D., Weng, C., Hosseinzadeh, S., Yang, D., ... & Weissman, J. S. (2022). Mapping transcriptomic vector fields of single cells. Cell, 185(4), 690-711.

[8] Olteanu, M., & Stefan, R. (2020). An exponential stability test for a messenger rna–micro rna ode model. University politehnica of bucharest scientific bulletin-series a-applied mathematics and physics, 82(4), 11-16.

[9] Chen, Z., King, W. C., Hwang, A., Gerstein, M., & Zhang, J. (2022). DeepVelo: Single-cell transcriptomic deep velocity field learning with neural ordinary differential equations. Science Advances, 8(48), eabq3745.

[10] Sun, X., Zhang, J., & Nie, Q. (2021). Inferring latent temporal progression and regulatory networks from cross-sectional transcriptomic data of cancer samples. PLoS computational biology, 17(3), e1008379.

[11] Farrell, S., Mani, M., & Goyal, S. (2022). Inferring single-cell dynamics with structured dynamical representations of RNA velocity. bioRxiv, 2022-08

[12] Li, Q. (2022). scTour: a deep learning architecture for robust inference and accurate prediction of cellular dynamics. bioRxiv, 2022-04

[13] Weinreb, C., Wolock, S., Tusi, B. K., Socolovsky, M., & Klein, A. M. (2018). Fundamental limits on dynamic inference from single-cell snapshots. Proceedings of the National Academy of Sciences, 115(10), E2467-E2476.

[14] Aliee, H., Richter, T., Solonin, M., Ibarra, I., Theis, F., & Kilbertus, N. (2022). Sparsity in Continuous-Depth Neural Networks. arXiv preprint arXiv:2210.14672.

[15] Bergen, V., Lange, M., Peidli, S., Wolf, F. A., & Theis, F. J. (2020). Generalizing RNA velocity to transient cell states through dynamical modeling. Nature biotechnology, 38(12), 1408-1414.

[16] Pramila, T., Wu, W., Miles, S., Noble, W. S., & Breeden, L. L. (2006). The Forkhead transcription factor Hcm1 regulates chromosome segregation genes and fills the S-phase gap in the transcriptional circuitry of the cell cycle. Genes & development, 20(16), 2266-2278.

[17] Desmedt, C., Piette, F., Loi, S., Wang, Y., Lallemand, F., Haibe-Kains, B., ... & TRANSBIG Consortium. (2007). Strong time dependence of the 76-gene prognostic signature for node-negative breast cancer patients in the TRANSBIG multicenter independent validation series. Clinical cancer research, 13(11), 3207-3214.

[18] Gillespie, M., Jassal, B., Stephan, R., Milacic, M., Rothfels, K., Senff-Ribeiro, A., ... & D'Eustachio, P. (2022). The reactome pathway knowledgebase 2022. Nucleic acids research, 50(D1), D687-D692.

[19] Ahnert, K., & Abel, M. (2007). Numerical differentiation of experimental data: local versus global methods. Computer Physics Communications, 177(10), 764-774.

[20] Chen, R. T. Q. (2021). torchdiffeq (Version 0.2.2) [Computer software]. `https://github.com/rtqichen/torchdiffeq`

[21] Chen, R. T., Rubanova, Y., Bettencourt, J., & Duvenaud, D. K. (2018). Neural ordinary differential equations. Advances in neural information processing systems, 31

[22] Hackett, S. R., Baltz, E. A., Coram, M., Wranik, B. J., Kim, G., Baker, A., ... & McIsaac, R. S. (2020). Learning causal networks using inducible transcription factors and transcriptome-wide time series. Molecular systems biology, 16(3), e9174.

[23] Van den Bulcke, T., Van Leemput, K., Naudts, B., van Remortel, P., Ma, H., Verschoren, A., ... & Marchal, K. (2006). SynTReN: a generator of synthetic gene expression data for design and analysis of structure learning

algorithms. BMC bioinformatics, 7(1), 1-12.

[24] Bhuva, D. D. (2017). SimulatorGRN [Computer software]. https://github.com/DavisLaboratory/SimulatorGRN

[25] Bhuva, D. D., Cursons, J., Smyth, G. K., & Davis, M. J. (2019). Differential co-expression-based detection of conditional relationships in transcriptional data: comparative analysis and application to breast cancer. Genome biology, 20(1), 1-21
